# Spontaneous social communication in laboratory mice - placing ultrasonic vocalizations in their behavioral context

**DOI:** 10.1101/2020.07.09.195362

**Authors:** Elodie Ey, Fabrice de Chaumont, Thomas Bourgeron

**Affiliations:** Human Genetics and Cognitive Functions, Institut Pasteur, UMR 3571 CNRS, Université de Paris, Paris, France

**Keywords:** Ultrasonic vocalizations, mouse, rodents, behavior, acoustic analyses, Shank3, autism, animal models, phenotyping

## Abstract

In their natural habitat, mice interact and communicate to regulate major functions, such as reproduction, group coordination, and protection. Nevertheless, little is currently known about their spontaneous emission of ultrasonic vocalizations (USVs), despite their broad use as a phenotypic marker in mouse models of neuropsychiatric disorders. Here, we investigated mouse spontaneous communication by coupling automatic recording, segmentation, and analysis of USVs to the tracking of complex behaviors. We continuously recorded undisturbed same-sex pairs of C57BL/6J males and females at 5 weeks and 3 and 7 months of age over three days. Males emitted only a few short USVs, mainly when isolated from their conspecific, whereas females emitted a high number of USVs, especially when engaged in intense dynamic social interactions. The context-specific use of call types and acoustic variations emerged with increasing age. The emission of USVs also reflected a high level of excitement in social interactions. Finally, mice lacking Shank3, a synaptic protein associated with autism, displayed atypical USV usage and acoustic structure, which did not appear in classical protocols, highlighting the importance of studying spontaneous communication. The methods are freely available for the research community (https://usv.pasteur.cloud).

## Introduction

Social communication regulates major biological functions under strong selective pressure, such as finding reproductive partners, raising progeny, group coordination for territory advertisement, and protection from predators (Bradbury and Vehrencamp, 2011). It is not yet clear whether the mechanisms underlying these functions are conserved during evolution, but an increasing number of species has been studied to identify genes and brain circuits related to social interaction and communication (e.g., (Arriaga et al., 2012; Chen and Hong, 2018; Kelley et al., 2020; Tu et al., 2019)). Such knowledge on the phylogeny and ontogeny of social communication abilities is also clinically important to better understand the causes underlying neuropsychiatric conditions. Indeed, several genes associated with autism have been mutated in animal models and lead to atypical social interactions (e.g., mice: (Crawley, 2012; Ey et al., 2011); rats: (Modi et al., 2018); drosophila: (Coll-Tané et al., 2019); monkeys: (Tu et al., 2019)), suggesting that at least certain social brain circuits are conserved during evolution. Hereafter, we will focus on mice, due to their broad use as models for neuropsychiatric disorders.

Mice are social animals, naturally living in *demes*, with a single dominant male, occasionally a few subordinate males, and several females occupying contiguous nests, but only a fraction of them reproduce (Palanza et al., 2005). This social organization leads mice to use tactile, olfactory, visual, and vocal (mostly in the ultrasonic range) communication in same-sex social interactions, male-female socio-sexual interactions, and mother-infant relationships (Brennan and Kendrick, 2006; Latham and Mason, 2004; Portfors, 2007). Tactile, visual, and olfactory cues are investigated through the observation of contacts, body postures, and marking behavior (e.g., (de Chaumont et al., 2019; Mathis et al., 2018)). In addition, mice emit ultrasonic vocalizations (USVs) that are interpreted as a proxy for vocal communication (Portfors, 2007; Sewell, 1970; Zippelius and Schleidt, 1956). USVs may also represent emotional reflectors of social interactions, reflecting the positive or negative valence of the interaction and the excitement of the interacting mice. Our understanding of mouse USVs nevertheless is still poor compared to our knowledge of vocal communication in other species, such as birds or primates (reviewed in (Naguib et al., 2009); for non-human primates in (Fischer and Hammerschmidt, 2010)).

Mouse USVs are generally investigated in contexts in which the motivation of the animals is controlled, through either social deprivation or the introduction of a sexual component to trigger a maximum quantity of USVs (reviewed in (Portfors, 2007)). During development, mouse pups isolated from the dam and littermates emit USVs as distress calls that trigger approach and retrieval by the mother (Sewell, 1970; Zippelius and Schleidt, 1956). In juveniles and adults, socially-deprived mice are highly motivated to interact with conspecifics; this maximizes the quantity of USVs emitted, which may be used to regulate close contacts and hierarchy (Ey et al., 2018; Moles et al., 2007). Finally, an estrous female, or at least its odor, stimulates adult males to vocalize (Chabout et al., 2015; Holy and Guo, 2005). Such male USVs are suspected to facilitate close body contact with the female (Hammerschmidt et al., 2009; Pomerantz et al., 1983); reviewed in (Egnor and Seagraves, 2016).

The contexts and functions of USVs have been studied in diverse conditions. USVs are mostly emitted during close contacts and approach behaviors in male-male (Ferhat et al., 2015a) and female-female (Ferhat et al., 2016a) interactions (including when one is socially deprived). Specific phases of courtship interactions trigger different call types (Hanson and Hurley, 2012; Nyby, 1983). Triangulation has been used to determine the emitter of the USVs (Warren et al., 2018a). In a group of four adult mice (two males and two females, aged 3-5 months) interacting for five hours after at least two weeks of social deprivation, both the females and males vocalized, more specifically during chasing (Neunuebel et al., 2015). The lowest call rates occurred when the animals were isolated from one another, whereas the highest call rates occurred when the mice were sniffing each other’s ano-genital region. Simple vocalizations (complex ones with frequency jumps or high modulations were discarded) are emitted in different proportions depending on the behavioral context (Sangiamo et al., 2020). In paired interactions, USVs trigger no behavioral variations in female-female interactions, but males accelerate when the females accelerate while vocalizing in male-female interactions (Warren et al., 2018b, 2020).

An ethological approach to mouse social communication should provide behavioral observations in contexts closer to natural conditions in which the animals should also be less stressed (*Peromyscus californicus*, *P. boylii*: (Briggs and Kalcounis-Rueppell, 2011; Kalcounis-Rueppell et al., 2006, 2010); *Mus musculus*: (Hoier et al., 2016a)). Here, we provide a detailed description of the spontaneous behaviors (see **Table I** in the **Methods** section) that are synchronized with USVs during an undisturbed period of three days and nights (**Figure 1a**) at three different ages (5 weeks and 3 and 7 months). We focused on same-sex interactions to avoid mixing sexual motivation in the factors that influenced vocal behavior. In addition, we investigated whether spontaneous USVs are perturbed in mice lacking Shank3, a glutamatergic synaptic scaffolding protein that we previously associated with autism (Durand et al., 2007; Leblond et al., 2014). We used Live Mouse Tracker (LMT) to provide a detailed behavioral description over long-term experiments (**Figure 1b-d**) (de Chaumont et al., 2019) and deliver the complete pipeline of recording and analyses, as well as an online application to automatically segment and analyze the USVs (https://usv.pasteur.cloud; **Figure 1e-f** and **Methods: Motivation to create a new recording and analysis pipeline)**. This first in-depth investigation of spontaneous mouse social communication and the resources presented here will be helpful for the community to design more sensitive tests to exploit the natural abilities of mice (Gerlai and Clayton, 1999; Warburton et al., 1988), as well as to detect and interpret new behavioral phenotypes in mouse models of neuropsychiatric disorders.

**Table I:** Definitions of behavioral events extracted by Live Mouse Tracker.

| Behavioral events |  |
| --- | --- |
| Name | Description |
| ***Stop isolated*** | The mouse is stopped (speed lower than 5 pixels per frame) and not in contact with any other mouse. |
| ***Move isolated*** | The mouse is moving at a speed higher than 5 pixels per frame and is not in contact with any other mouse. |
| ***Break contact*** | The mouse breaks a contact with another mouse and moves away from this mouse. |
| ***Get away*** | The mouse moves away from the other mouse (any distance) |
| ***Social approach*** | The mouse is approaching another one, i.e. the distance between the two animals shortens, the speed of the mouse is higher than the speed of the approached mouse, and the distance between the two animals is shorter than two mean body lengths (of the approached animal). This approach does not necessarily lead to a contact. |
| ***Approach reared mouse*** | The mouse is approaching another one, i.e. the distance between the two animals shortens, the speed of the mouse is higher than the speed of the approached mouse, and the distance between the two animals is shorter than two mean body lengths (of the approached animal). The approached mouse is rearing, i.e. the slope between the tail basis and the nose point. This approach does not necessarily lead to a contact. |
| ***Approach contact*** | The mouse is approaching another one and makes contact with it. |
| ***Contact*** | The mouse is in contact (i.e. the two masks have one common pixel) with the other mouse. |
| ***Oral-oral contact*** | The mouse is sniffing the oral region of another mouse, i.e. the two nose points are less than 15 pixels (26 mm) from one another. |
| ***Oral-genital contact*** | The mouse is sniffing the ano-genital region of another mouse, i.e. the nose point of the first mouse is within 15 pixels (26 mm) from the tail point of the other mouse. |
| ***Side by side contact*** | The side of a mouse is within 30 pixels (52 mm) from the side of the other mouse; both animals are oriented in the same direction. |
| ***Side by side contact (opposite way)*** | The side of a mouse is within 30 pixels (52 mm) from the side of the other mouse; both animals are oriented in opposite directions. |
| ***Sequence oral oral - oral genital*** | The mouse first sniffs the nose of the other mouse and then sniffs the ano-genital region of this mouse. |
| ***Sequence oral genital - oral oral*** | The mouse first sniffs the ano-genital region of the other mouse and then sniffs the nose of this mouse. |
| ***Follow zone isolated*** | The mouse is walking alone directly in the path of another mouse. Both mice move at a speed higher than 5 pixels per frame, with an angle between their direction vectors smaller than 45°, and the distance between their centers of mass shorter than two mean body lengths. |
| ***Train2*** | The mouse is following another mouse (speed higher than 5 pixels per frame) while sniffing its ano-genital region. |
| ***Longchase*** | A chase sequence involving "Contact", "FollowZone Isolated", "Train2" and "Break contact" as a sequence. |

**Figure 1.**
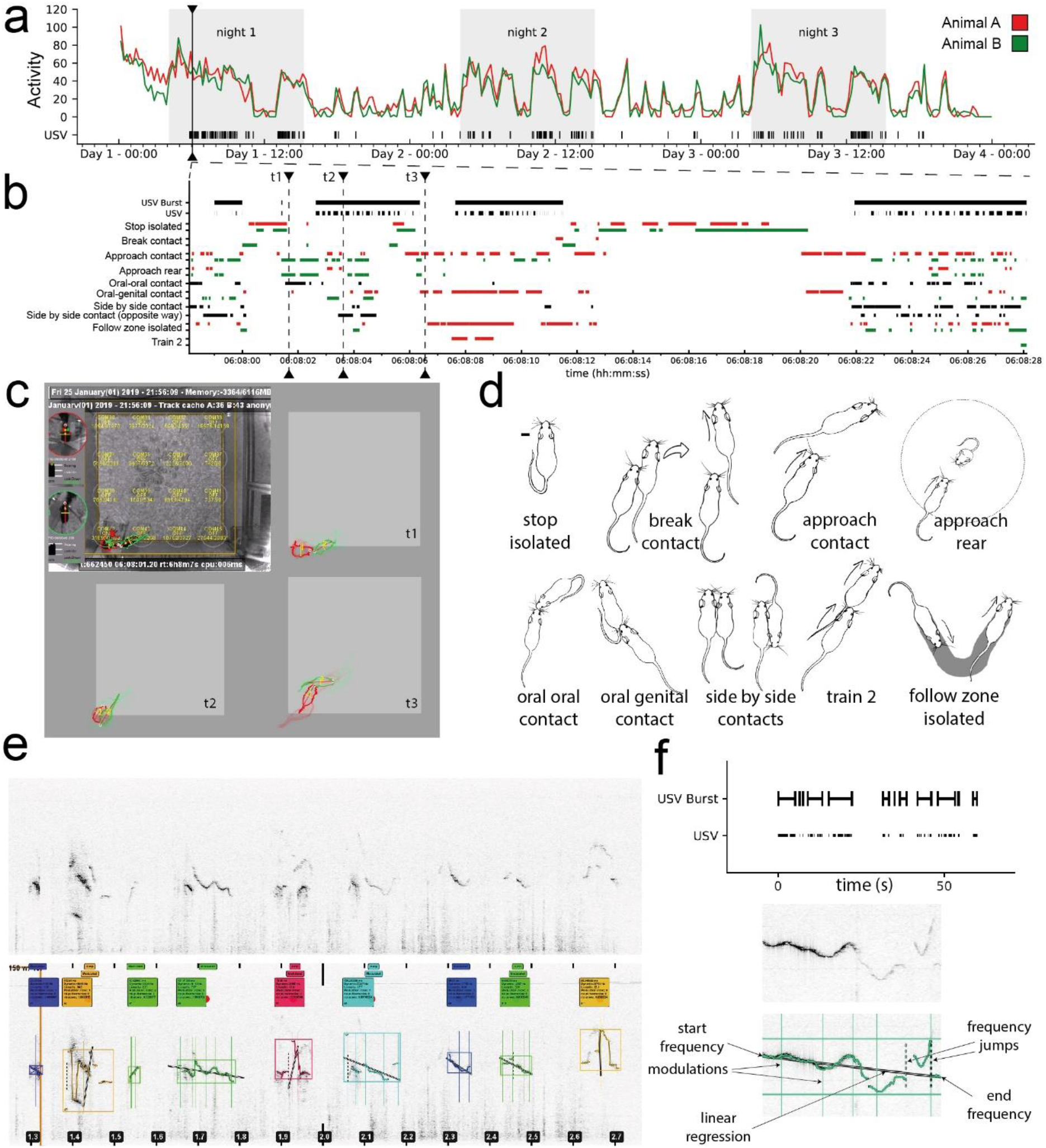
Overview of the study of spontaneous USVs in same-sex pairs of mice. (a) Example of the activity timeline of a pair of three-month-old C57BL/6J female mice recorded for three days, displaying the distance travelled per 15-min time bin (m) and the emission of USVs. The black arrow indicates the timepoint at which the timeline is zoomed in (b). (b) Timeline of 30 s of the experiment (day 1, 6 a.m.), with USVs, USV bursts, and the major events. (c) Set-up used to record the spontaneous USVs and behaviors of a same-sex pair of mice using Live Mouse Tracker (LMT) and examples of time points (t1, t2, t3) displayed on the timeline in (b). (d) Examples of behaviors automatically extracted using LMT; see definitions in (de Chaumont et al., 2019) and **Methods**. (e) Original spectrogram (upper panel) and segmented USVs within this spectrogram (lower panel) in a USV burst. (f) Temporal organization of USVs in USV bursts with inter-burst intervals longer than 1 s (upper panel), and acoustic variables extracted from each USV (lower panel).

## Results

### A platform to analyze USVs and their synchronization with behaviors

We needed to automatically detect USVs within background noise (i.e., animals moving in the bedding, continuous electronic or ventilation noises) and to extract their acoustic features, all these aspects being synchronized with behavioral monitoring. In our system, sounds were recorded using Avisoft SASLab Pro Recorder and synchronized with behavioral monitoring using LMT (**Supplementary methods - Wave record and synchronization; Figure 1a-d**). Mechanical noise (e.g., amplitudes at continuous frequencies from electronic devices or ventilation) was removed (**Supplementary methods - filtering mechanical noise**) and the USVs extracted (**Figure 1e**; **Supplementary methods - USV detection / USV detection validation**). The USVs were then synchronized with behaviors and the acoustic characteristics (**Figure 1f**) extracted and stored as metadata in the database (**Supplementary methods - USV audio feature extraction**). We provide a library in Python to access and query these data. We developed an online application that allows the segmentation and analysis of wave files without software installation (http://usv.pasteur.cloud). Annotated spectrograms are generated online to allow visual inspection of the results. We also provide online examples.

### Pairs of male mice emit few USVs and not preferentially during social investigations

Throughout the 72 hours of continuous observation, pairs of familiar male mice emitted USVs mainly during their active nocturnal hours (**Supplementary Figure S1**). Over the three days, male mice emitted a low and constant number of USVs and USV bursts (i.e., sequences of USVs separated by intervals of less than 1 s) at each age relative to female mice (**Figure 2a**). No significant effect of age was detected (paired Wilcoxon test: all p-values > 0.05). USV rates (i.e., the number of USVs emitted in one behavioral context divided by the total duration of this behavioral context) emitted by pairs of males remained low in all behavioral contexts (**Figure 2b**, upper panel).

**Figure 2.**
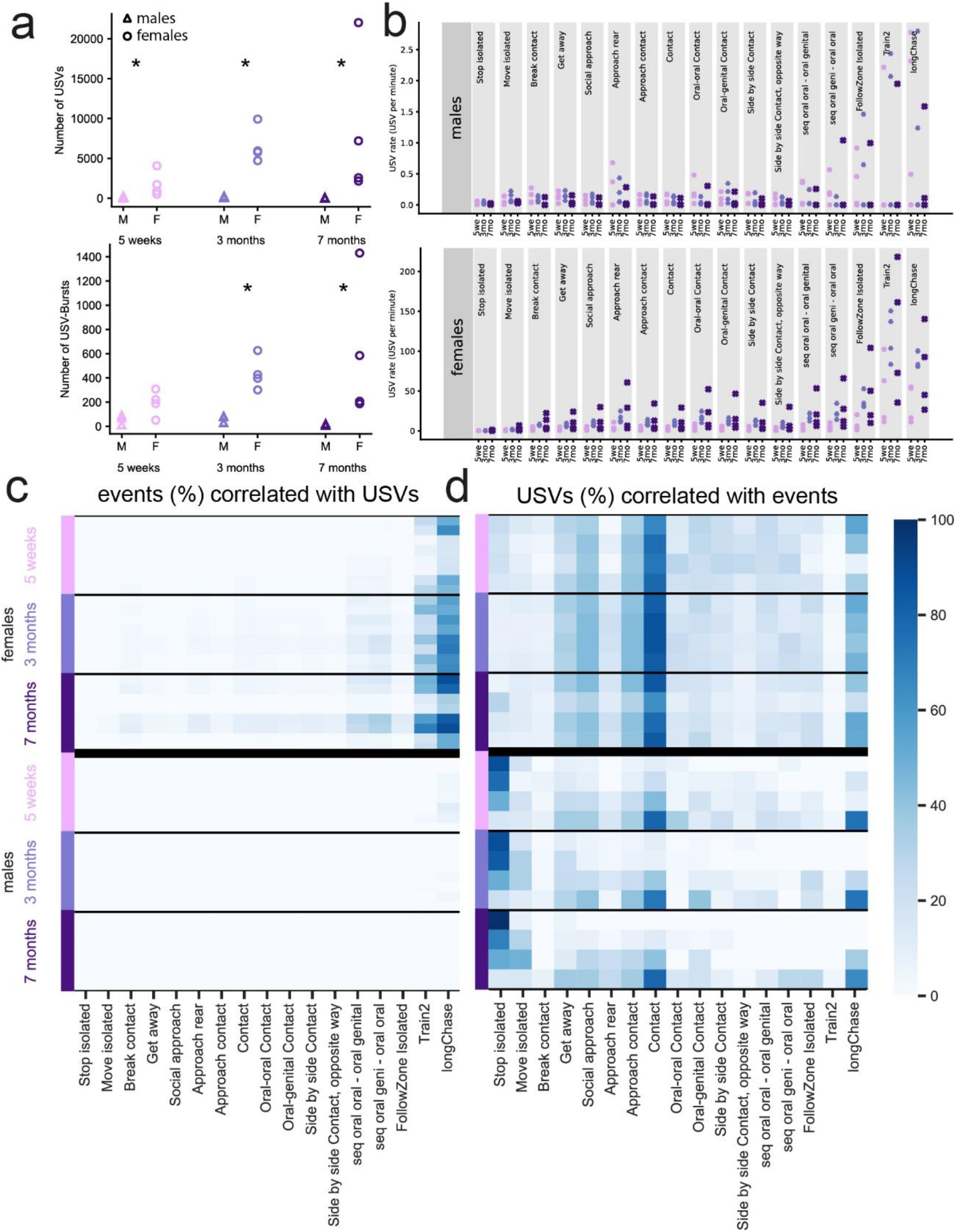
Relationships between USVs and behaviors in WT same-sex pairs. (a) Total number of USVs (upper panel) and USV bursts (lower panel) recorded over three days in each age class for both males (M) and females (F). (b) Rate of emission of USVs for various behaviors by males (upper panel) and females (lower panel) aged five weeks, three months, or seven months. (c) Proportion of the frames of the various behavioral events overlapping with USVs for each age and sex. The behavioral event is the reference. For dark blue, the figure reads as if 80% of the duration of Train2 was accompanied by USVs. (d) Proportion of the frames of the USV events overlapping with various behavioral events for each age and sex. The USV event is the reference. For dark blue, the figure reads as if 80% of the USV duration was accompanied by stop isolated (for males) or contact (for females).

Less than 10% of the behavioral events overlapped with the USVs (**Figure 2c**). However, most of the few USVs emitted by males overlapped with stop isolated (**Figure 2d**). Given the low number of USVs emitted by males during same-sex interactions, we investigated female USVs in more detail.

### Female mice emit spontaneous USVs during dynamic social interactions at an increasing rate with age

As for males, pairs of female mice emitted USVs principally during their active nocturnal hours (**Supplementary Figure S2**). In contrast to males, the number of USVs emitted tended to increase with age (paired Wilcoxon test between 5 weeks and 3 months: U = 0, p = 0.125; **Figure 2a**, upper panel). At five weeks of age, females emitted USVs at the highest rates during Train2 and longChase behaviors and the lowest during stop isolated events (**Figure 2b**, lower panel). From three months of age on, this pattern was complemented by high USV rates during approach reared mouse behavior and sequences of oral-genital sniffing (**Figure 2b**, lower panel).

In contrast to males, most USVs were emitted during contact, approach contact, social approach, and longChase events (**Figure 2d**). Up to 80% of Train2 and longChase behavioral events were accompanied by USVs (**Figure 2c**). Overall, female mice emitted USVs at the highest rates during intense dynamic (i.e., involving movements) social interactions. Interestingly, USVs and USV bursts appeared to be emitted once direct social interactions had already begun or during direct social interactions and preceded the termination of such social interactions. This temporal organization was not clearly established in immature mice (5 weeks of age; **Supplementary Figure S3**). Overall, these results suggest specificity in the contexts of emission of USVs in female mice, which we then explored.

### Acoustic traits are context-specific in mature female mice

We investigated whether the acoustic features (duration, frequency characteristics, frequency range, modulations, harshness, and slope; see definitions in **Methods**) of the USVs were distinctive according to the context of emission at different ages. First, acoustic variables varied continuously with increasing age across all contexts. USVs became longer, more modulated, and showed lower frequency characteristics with increasing age (**Figure 3a** for duration, frequencyTV and min frequency and **Supplementary Figure S4**). At five weeks of age, most acoustic variables did not vary between contexts (**Figure 3b** and **Supplementary Figure S5**). Only USVs emitted during stop isolated events or urination (see **Supplementary data - Urination sequences**) differed from those emitted in the other contexts by their shorter duration, and lower frequency dynamic, diffStartEndFrequency, frequencyTV, linearity index, number of modulations, number of jumps, and harshIndex. USVs emitted during close contact (approach, oral-oral, oral-genital, and side-side) did not differ from each other but displayed significantly higher frequency characteristics than USVs emitted during stop isolation, break contact, and approach reared mouse, as well as during follow, Train2, and longChase behaviors.

**Figure 3.**
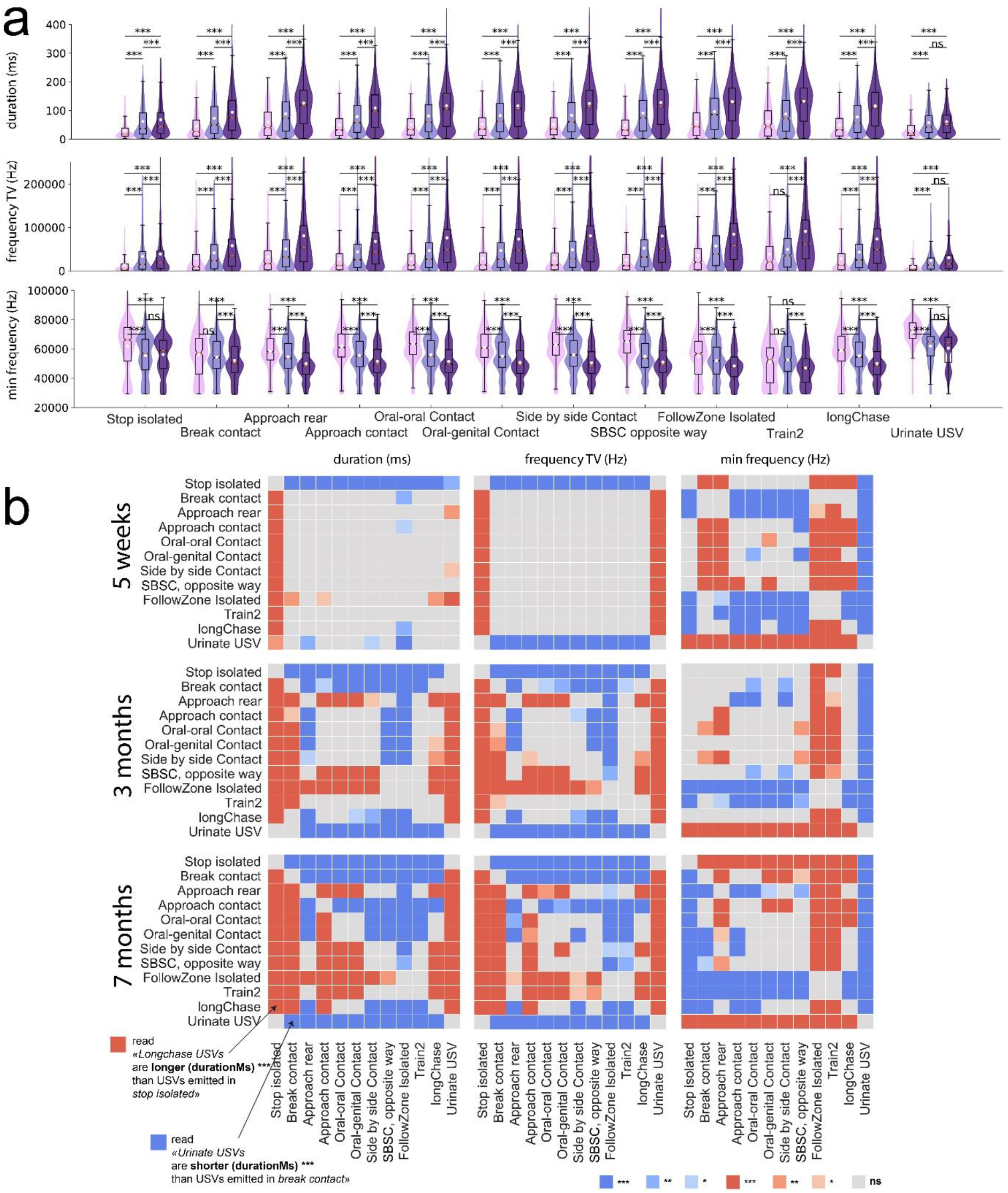
Variations in the acoustic features of USVs emitted in different behavioral contexts in female WT mice. (a) Variations in the acoustic structure of USVs with increasing age of WT female mice. Three acoustic traits are depicted: duration, frequencyTV, and min frequency. Comparisons between age classes were performed using Mann-Whitney U-tests, with P-values adjusted by a Bonferroni correction in each context. SBSC = side-by-side contact. (b) Variations of the acoustic structure of USVs depending on their context of emission. Three acoustic traits are depicted as examples. The color intensity represents the significance levels of the tests and blue and red represent the direction of the difference: red: the acoustic trait of USVs emitted in the y-label event is higher than the same acoustic traits in USVs emitted in the x-label event; blue: the acoustic trait of USVs emitted in the y-label event is lower than the same acoustic traits in USVs emitted in the x-label event. For example, USVs emitted in stop isolated events were significantly shorter than USVs emitted in all other contexts, except urinating sequences, which were accompanied by significantly longer USVs. P-values adjusted for multiple testing by a Bonferroni correction: combination of 2 events tested * number of acoustic traits * age: ((12*(12-1)) / 2) * 16 * 3 = 3,168.

The link between acoustic features of USVs and behaviors became more specific with increasing age (**Figure 3b** and **Supplementary Figure S6**). At three months of age, USVs emitted during urination and stop isolated events were similar (**Figure 3b** and **Supplementary Figure S6**). They were short, with a lower diffStartEnd, frequency dynamic, frequencyTV, number of jumps, number of modulations, and harshIndex. The main differences between stop isolated events and urination were that the USVs emitted during urination displayed a lower diffStartEnd, a higher end, max, min and mean frequency, and a higher mean power than USVs emitted during stop isolated events. Interestingly, USVs emitted during stop isolated and break contact events were similar, with a shorter duration, reduced frequencyTV, and reduced number of modulations than USVs emitted in other contexts. USVs emitted at three months of age during Train2 and follow events did not differ from each other (**Figure 3b** and **Supplementary Figure S6**), despite the fact that these two behaviors differed by the presence (Train2) or absence (follow) of contact in the pursuit. They were comparable in their low end, min, and mean frequencies, as well as high harshIndex relative to USVs emitted in most other behavioral contexts. USVs emitted during side-side contact opposite events differed from USVs emitted during Train2 and follow events by a higher end and min frequency, and lower frequency dynamic, frequencyTV, linearity index, and number of jumps, as well as by a higher mean and start frequency than USVs emitted during following behaviors. When mice were close to each other, the USVs were rarely specific to the different types of contacts (**Figure 3b** and **Supplementary Figure S6**). Indeed, variations in duration, frequency characteristics, frequency modulations, slope, and harshIndex between USVs emitted in approach contact, oral-oral, oral-genital, and side-side contacts were not significant. USVs emitted during longChase events resembled those emitted during break contact events but were less clearly defined. This may be due to the fact that longChase involved a mix of events, such as approach, contact, and break contact.

At seven months of age, the acoustic structure of the USVs became even more specific to each context and the context-specificity observed at three months of age became more pronounced (**Figure 3b** and **Supplementary Figure S7**). USVs emitted in follow and Train2 behaviors remained acoustically indistinguishable from each other but displayed significantly larger frequency dynamic, frequencyTV, linearity, and harshIndex, as well as lower end, min, and mean frequencies than USVs emitted in other contexts.

### Distinctive USVs occur during specific behaviors

We investigated whether distinctive USVs appear more frequently than expected during specific behavioral events. We first observed single USVs that started at a very high frequency and covered a large frequency range (see **Supplementary material – Specific USV types, Isolated vocalizations with high peak frequency**). These USVs were emitted when a mouse was jumping against the wall (descending phase) or jumping down from the Plexiglas house. This occurred for all ages and both sexes. This may be due to the mechanical lung compression when mice hit the ground after a jump. We also identified USVs that were emitted in bursts during urination sequences (see **Supplementary material – Specific USV types, Urination sequences**). The acoustic structure of these USVs was highly characteristic (see **Figure 3**).

USVs characterized by harsh elements (i.e., without pure peak frequency in some parts; **Figure 4a**, left panel) were present in all age groups for the two sexes. At five weeks of age, harsh USVs represented approximately 20% of all USVs for all behavioral events (**Figure 4a**, right panel). At 3 and 7 months of age, the proportion of harsh/pure USVs tended to be balanced (50% each) for most behaviors, except during urination and stop/move isolated evens, for which harsh USVs (30%) were less highly represented than pure USVs (70%; **Figure 4a**, right panel). From 3 to 7 months of age, the proportion of harsh USVs increased during intense social events, such as Train2, follow, getaway, and approach rear (**Figure 4a**, right panel).

**Figure 4.**
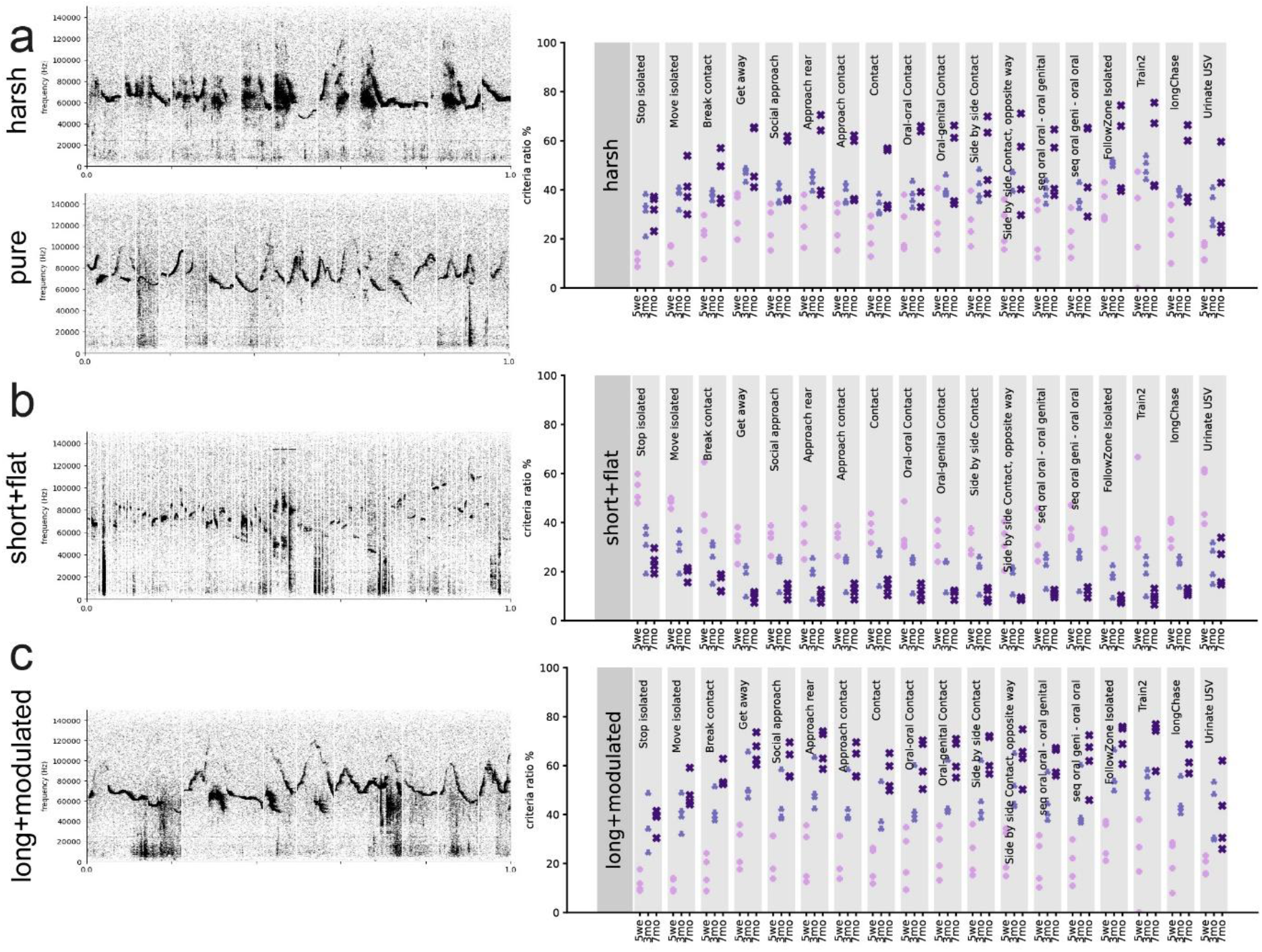
Context-dependent occurrences of specific USV types. (a) Compilation of spectrograms of harsh USVs (left, upper panel) and pure USVs, without harsh components (left, lower panel), from a pair of WT mice aged three months and the proportion of harsh USVs in the different behavioral contexts observed for the three age classes (right). (b) Compilation of spectrograms of short and flat (i.e. reduced frequency modulations) USVs from a pair of WT mice aged three months (left) and the proportion of such short and flat USVs in the different behavioral contexts observed for the three age classes (right). (c) Compilation of spectrograms of long and modulated USVs from a pair of WT mice aged three months (left) and the proportion of such long modulated USVs in the different behavioral contexts observed for the three age classes (right). (FFT-length: 1024 points, 16-bit format, 300 kHz sampling frequency, 75% overlap).

USVs with a short duration and reduced frequency range (short-flat) were more frequently emitted during stop and move isolated events than social contexts for all age classes (**Figure 4b**). The proportion of these short-flat USVs was higher for immature (40%) than older mice, which emitted short-flat USVs in less than 20% of vocalizations at seven months of age. Overall, the vocal repertoire of fully-grown mice is richer than that of immature mice in similar behavioral contexts.

Finally, we detected long-modulated USVs emitted in low proportions (< 35%) in all behavioral contexts at five weeks of age (**Figure 4c**), consistent with the simpler USV repertoire of the immature mice suggested above. At three months of age, the proportion of such long-modulated USVs was close to 50% in all contexts, except urination and isolated move and stop events, for which it was lower. This was even more evident at seven months of age. Remarkably, these long-modulated USVs were emitted more than any other USV type in social contexts, whereas they were more rarely emitted during isolated behaviors and urination.

Overall, the use of complex USVs increased with increasing age, more specifically during socially intense behaviors. Simple and short USVs were more highly associated with nonsocial behaviors, whereas complex USVs, such as harsh and modulated ones, were emitted during socially intense behaviors.

### Higher speed and longer duration of behaviors during which USVs are emitted may reflect high excitement

We next hypothesized that behaviors accompanied by USVs were more intense (i.e., longer and with higher mean speed) than behaviors not accompanied by USVs. As a proxy for excitement, we considered the speed of the animals. We thus quantified the mean speed of the animals during behavior accompanied or not by USVs, as well as their duration.

Our hypothesis was not verified for longChase and Train2 events in any age class at the individual level. Indeed, within each individual, although the speed of Train2 was the highest of all events, the duration of the event and the speed of the mice did not vary much, regardless of whether USVs were emitted or not (**Figure 5a**). In contrast, follow (**Figure 5b**), approach contact (**Figure 5c**), contact, getaway, and oral-genital contact (data not shown) showed a significantly longer duration and higher speed when accompanied by USVs, in agreement with our hypothesis at the individual level. The presence of USVs was accompanied by higher speed but not necessarily longer duration for break contact events (**Figure 5d**). Overall, the relationship between USV emission and behavioral display (speed and duration of the event) suggests higher excitement in simple (e.g., follow, approach contact, break contact) but not complex behavioral macro-events (e.g., longChase) or very intense social investigation, such as Train2.

**Figure 5.**
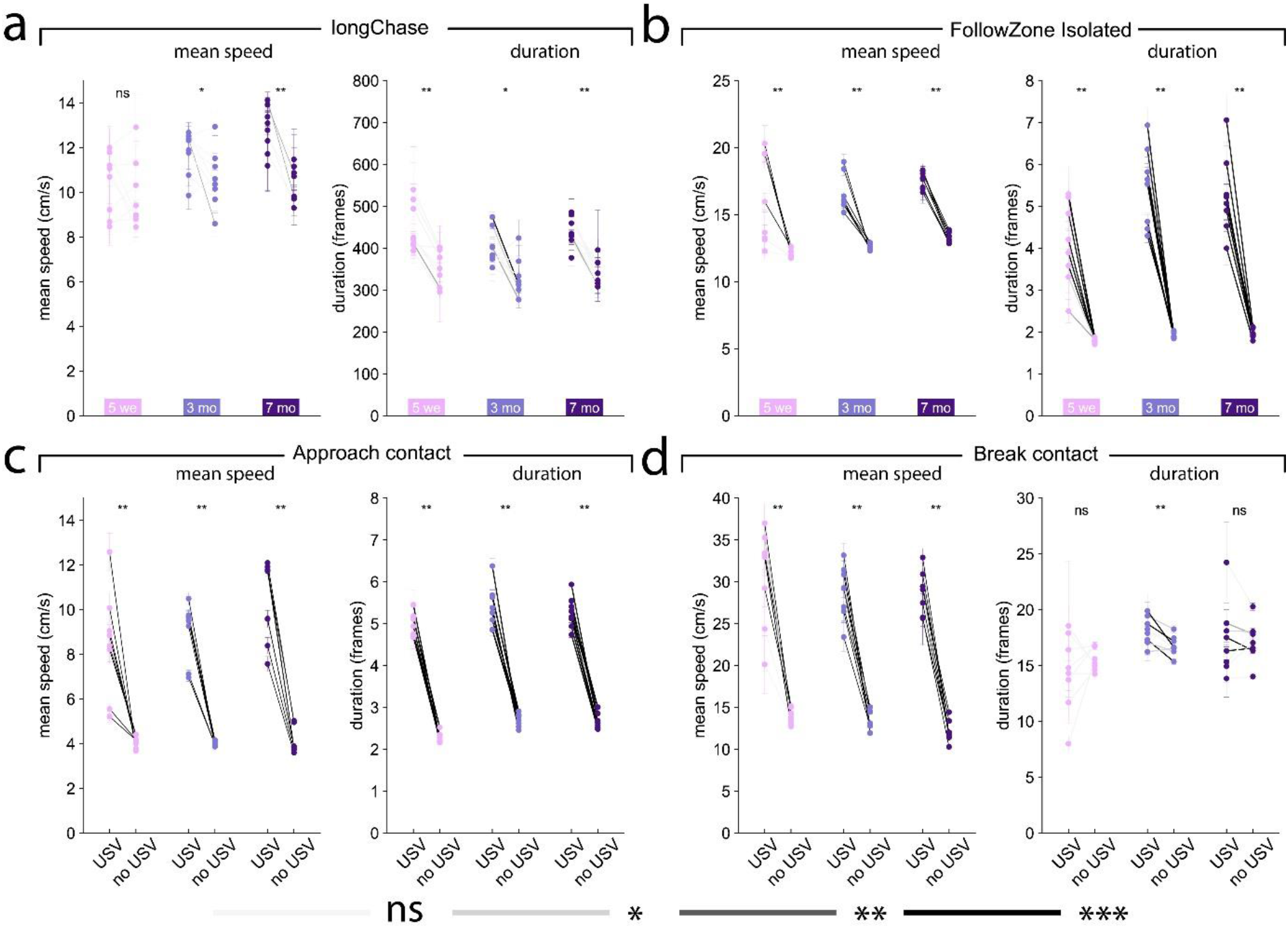
Variations in mean speed and duration of each behavioral event according to the presence or absence of USVs. Variations of the mean speed of the animal performing the behavior (left panel) and of the event duration (right panel) for (a) longChase, (b) follow, (c) approach contact, and (d) break contact. Paired comparisons between behaviors with and without USVs were conducted at the group level using the paired Wilcoxon test for each age class, separately. Significant differences at the individual level using Mann-Whitney U-tests after Bonferroni correction are depicted by the color of the segments linking the mean value, with and without USVs. light grey: not significant, medium grey: p < 0.05, dark grey: p < 0.01, black: p < 0.001.

### From USVs to USV bursts

In both males and females, USVs were gathered in bursts. According to the distribution of intervals between USVs, we defined that USVs belonged to the same bursts if the intervals between USVs were shorter than one second (**Supplementary methods – Burst definition**). For males, no significant age effect was detected for the number of USVs per USV burst. In females, there was only a trend for an increase in the number of USVs per USV burst with increasing age. Indeed, females aged five weeks tended to emit USV bursts with fewer USVs than three-month-old females (paired Wilcoxon test: W = 0, p = 0.125), but the number of USVs per USV burst did not differ between 3 and 7 months of age (**Supplementary Table I**). In addition, the major difference in the use of vocal signals observed between males and females in terms of USVs was also observed in terms of USV bursts for all ages. Females tended to emit USV bursts containing more USVs than males (Mann-Whitney U-test: 5 weeks: U = 1.0, p = 0.061; 3 months: U = 0, p = 0.030; 7 months: U = 3, p = 0.194).

### Context-dependent variations in burst characteristics

We tested whether the USV bursts display characteristics that vary depending on the behavioral context in which they are emitted. As for USVs, the acoustic features and context-specificity of USV bursts increased with age (**Supplementary Figure S8**). Overall, USV bursts emitted during behavioral events of isolated mice (stop, urination) were short (both in duration and the number of USVs per burst) and included short USVs of stable duration relative to USV bursts emitted in other contexts. In contrast, USV bursts emitted during intense social investigation (Train2) were longer (both in duration and the number of USVs per burst) and included long USVs of highly variable duration relative to USV bursts emitted in other contexts. Although most of these traits were already visible at five weeks of age, context-specificity during social behavioral events (follow, nose-nose contacts, oral-genital contacts, side-side contacts) was only significantly detected from three months of age on.

### Intonation within USV bursts are conserved with increasing age

As USVs were emitted sequentially in USV bursts, we investigated whether the USVs within bursts were organized. We thus separated short USV bursts with eight or less USVs from long USV bursts that included nine or more USVs (**Supplementary Figure S9**). Fitting of a simple linear trend for each acoustic trait of each USV burst showed a decreasing trend of harsh index, frequencyTV, and linearity and an increasing trend of frequency characteristics in almost all contexts in long bursts at 3 and 7 months of age (**Supplementary Figure S10a**). In contrast, there was a decreasing linear fit of the number of modulations, the number of jumps, and harsh index in almost all behavioral contexts for short bursts (**Supplementary Figure S10b**). We next examined the intonation in the first and second half of long bursts (**Methods - Intonation within USV bursts**). Overall, variations in the ∩-shape were the most highly represented, whereas USV bursts with stable features in one or both halves were the least observed (**Figure 6a**). Variations in the \\− and //−shape were equally shared. For example, at three months of age, between 35 and 50% of the USV bursts were characterized by variations in the ∩-shape in duration, mean power, frequencyTV, frequency dynamic, mean frequencyTV, harshIndex, number of modulations, and linearity index. Variations in the U-shape were mostly observed for end and mean frequencies. Such organization in the intonation was similar for all age classes. A stable harshIndex or number of jumps between the two halves of the USV bursts were observed in 15 to 30% of the USV bursts for all age classes and these traits were the only ones to remain stable in a substantial number of USV bursts. Quantification of the most highly represented styles with one acoustic trait at three months of age (**Figure 6b**, middle panel) showed the frequencies of occurrence of the different styles to be similar at seven months of age (**Figure 6b**, right panel) but slightly less organized at five weeks of age (**Figure 6b**, left panel).

**Figure 6.**
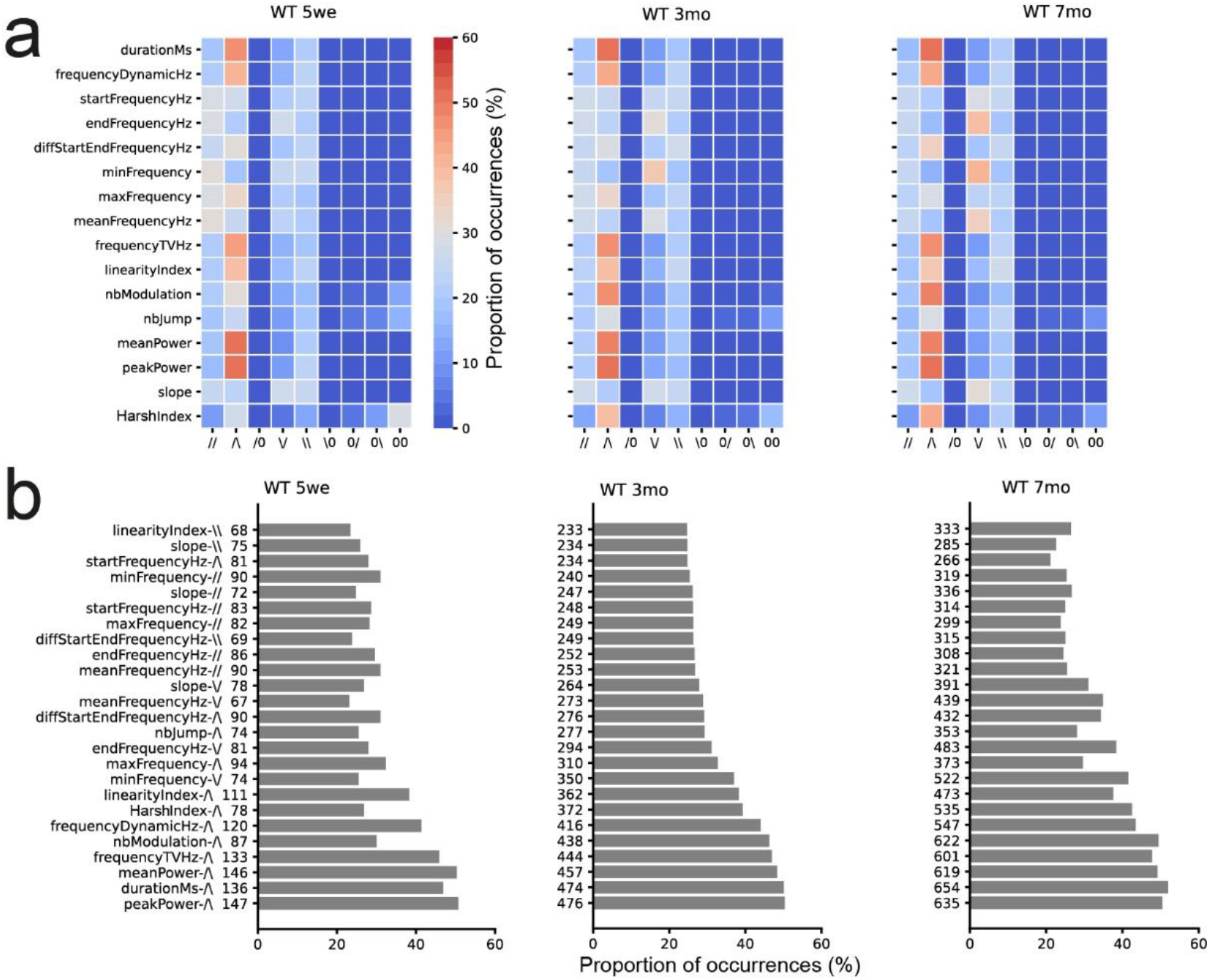
Intonation within USV bursts according to age. (a) Proportion of occurrences of the different possibilities of intonation within long USV bursts in WT mice aged five weeks, three months, and seven months of age. The heatmap reads as the duration varies in ∩-shape in 45% of USV bursts at five weeks of age. (b) Frequency of occurrence of the 20 most frequent USV burst styles with one acoustic trait of female WT mice, according to age.

We hypothesized that certain combinations of intonations are specific for certain types of events and, more specifically, that the styles of bursts are different between intense social events, such as Train2 and longChase, and non-social events, such as stop isolated, move isolated, and break contact or getaway. We tested this hypothesis in three-month-old WT females. We computed the distribution of the 88 most frequent styles (i.e., intonation combinations) over the various types of events. We used combinations of one, two, or three acoustic traits with the different types of intonations. We did not observe differences between intense social events and non-social events (data not shown).

#### Spontaneous vocal behavior in *Shank3* mutant females

### Lower activity and atypical social interactions in Shank3 mutant females

We examined the spontaneous social interaction profiles of *Shank3*^*−/−*^ mutant females in early adulthood (3 months, as in WT mice). *Shank3*^*−/−*^ mice tended to spend less time in approach reared mouse, but more time in stop in contact and oral-oral contact than age-matched WT females (**Supplementary Figure S11**). As previously observed, *Shank3*^*−/−*^ mice also tended to be less active (reduced distance travelled) than WT females. The mean durations of oral-oral contact, side-side contact, and side-side contact in opposite direction were longer for *Shank3^−/−^* mice than WT mice. In contrast, the mean durations of break contact, move in contact, and USV bursts were shorter for *Shank3^−/−^* mice than WT mice (**Supplementary Figure S12**).

### Shank3^−/−^ female mice emit fewer USVs than WT mice after the initial exploration phase

The number of USVs recorded for *Shank3*^−/−^ females over the three days was not statistically different from that of USVs recorded for WT females (Mann-Whitney U-test: U = 8.0, p = 0.228; **Supplementary Figure S13a**). However, *Shank3*^−/−^ mice emitted significantly fewer USVs than WT mice during the second night (U = 2.0, p = 0.021). The rates of USVs (**Supplementary Figure S13b**), contexts of emission (**Supplementary Figure S13c-d**), and temporal organization of behavioral events associated with USVs (**Supplementary Figure S13e**) were similar between *Shank3*^−/−^ and WT mice.

### Shank3^−/−^ female mice emit USV bursts with fewer but longer and more modulated vocalizations than WT mice in most contexts

We compared the acoustic features of individual USVs (**Figure 7a**) and USV bursts (**Figure 7b**). USVs emitted by *Shank3*^−/−^ mice were significantly longer, with a higher diffStartEnd and higher number of modulations, than those emitted by WT mice in all contexts, except side-side opposite and stop isolated behaviors (**Figure 7a;** see also **Supplementary Figure S14**). *Shank3*^−/−^ mice emitted USVs with a lower end frequency and lower power characteristics than WT mice, in most contexts. These differences in the power characteristics were not related to differences in the proximity to the microphone (**Supplementary Figure S15a-b**). The slightly smaller body volume (estimated through body surface on the tracking mask; **Supplementary Figure S15c**) of *Shank3*^−/−^ mice relative to that of WT mice could partially explain the reduced power of *Shank3*^−/−^ mouse USVs.

**Figure 7.**
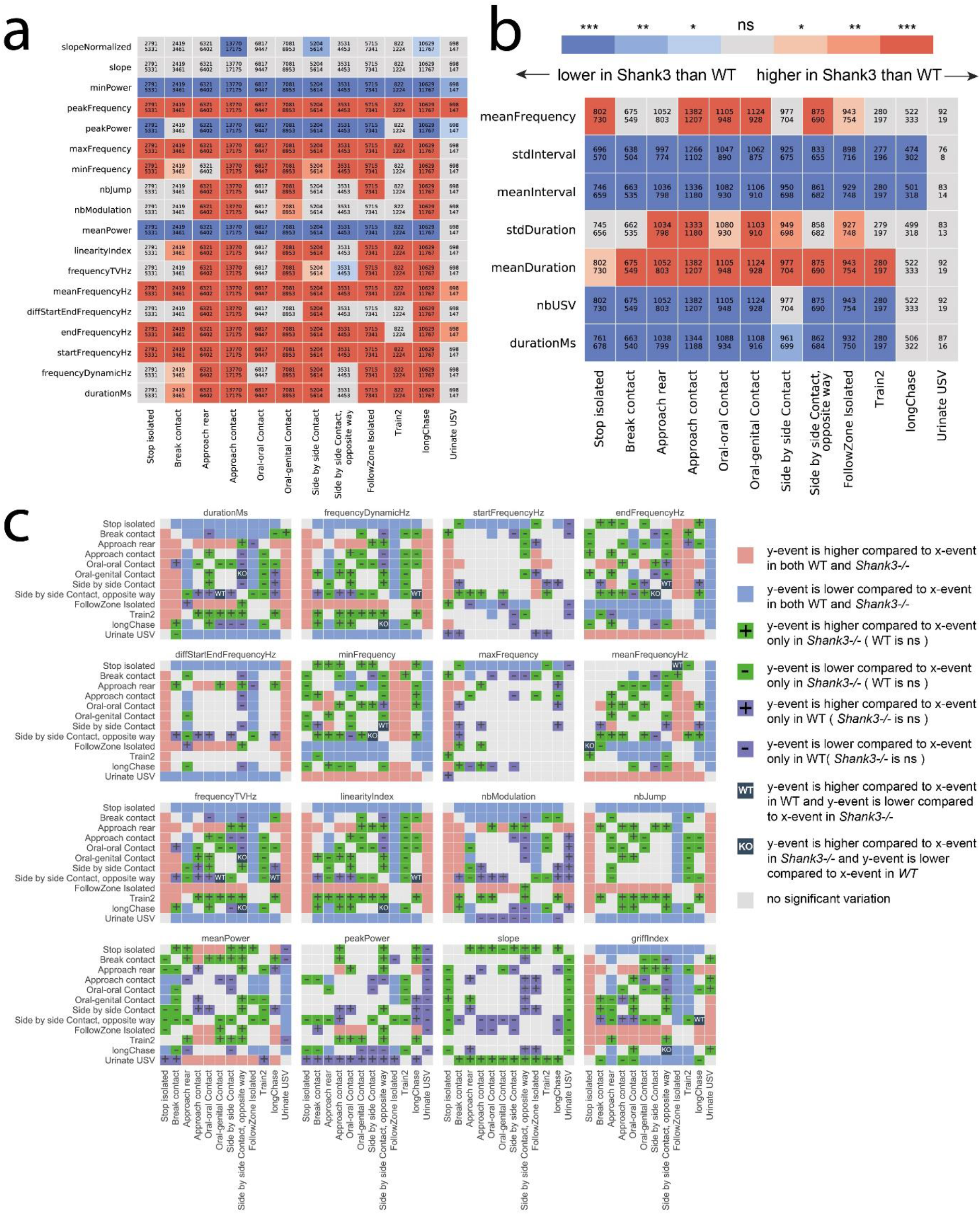
(a) Comparison of USV acoustic variables between *Shank3*^−/−^and B6 mice in the various contexts. Red: the value is significantly higher in *Shank3*^−/−^ than in B6 mice; blue: the value is significantly lower in *Shank3*^−/−^ than in B6 mice (value = nb of stars, Bonferroni corrected for contextEvent * vocTraits). (b) Comparison of burst characteristics between WT and *Shank3*^−/−^ mice according to the behavioral context for burst duration, the number of USVs per USV burst, the mean duration of USVs within a burst, the standard deviation of the duration of USVs, the mean interval between USVs within a USV burst, the standard deviation of intervals between USVs within USV bursts, and the mean peak frequency over all USVs within the burst. The color intensity depicts the significance of the Mann-Whitney U-tests and the colors represent the direction of the difference: red: higher in *Shank3*^−/−^than WT; blue: lower in *Shank3*^−/−^ than in WT. (c) Context-specific acoustic variations common to *Shank3*^−/−^ and WT mice, as well as context-specific acoustic variations different between *Shank3*^−/−^ and WT mice. The red color depicts that USVs emitted in the y-axis behavior display a higher trait value than USVs emitted in the x-axis behavior, and this variation in conserved between *Shank3*^−/−^ and WT mice; the blue color depicts that USVs emitted in the y-axis behavior display a lower trait value than USVs emitted in the x-axis behavior, and this variation in conserved between *Shank3*^−/−^ and WT mice; the green color depicts context-specific variations present only in *Shank3*^−/−^ mice (+: USVs emitted in the y-axis behavior display a higher trait value than USVs emitted in the x-axis behavior; -: USVs emitted in the y-axis behavior display a higher trait value than USVs emitted in the x-axis behavior); the violet color depicts context-specific variations present only in WT mice (+: USVs emitted in the y-axis behavior display a higher trait value than USVs emitted in the x-axis behavior; -: USVs emitted in the y-axis behavior display a higher trait value than USVs emitted in the x-axis behavior). Black squares represent opposite variations between *Shank3*^−/−^ and WT mice.

*Shank3*^−/−^ mice emitted USV bursts that contained fewer (*Shank3*^−/−^: 10.8 ± 11.5 USVs per USV burst; WT: 15.0 ± 17.4 USVs per USV burst; Mann-Whitney U-test: U = 2355,001, p-value = 1.61 × 10^−12^) but longer USVs separated by shorter intervals than WT mice (**Figure 7b**; see also **Supplementary Figure S16**). These differences occurred across all behavioral situations, except urination and longChase events, for which no differences were observed between *Shank3*^−/−^ and WT mice.

Context-related variations in the USVs emitted during stop isolated, follow, and urination were similar between *Shank3^−/−^* and WT mice for most acoustic variables (**Figure 7c**). Context-related variations were more present in *Shank3^−/−^* than in WT mice for the frequency dynamic, end, min, and mean frequency, linearity, number of jumps, mean power, and harshIndex, whereas context-related variations were more present in WT than *Shank3^−/−^* mice for the number of modulations and peak power (**Figure 7c**). Interestingly, opposite variations between *Shank3^−/−^* and WT mice were mostly observed for USVs emitted during side-side contacts, longChase, and follow behaviors, which are amongst the most intense social interactions (**Figure 7c**).

### The frequency and location of USV emission during urination is altered in Shank3^−/−^ female mice

Concerning USVs emitted during specific behavioral contexts, the sounds emitted by mice jumping down the walls and the harsh USVs, short-flat USVs, and long-modulated USVs were preserved in *Shank3*^−/−^ mice (**Supplementary Figure S17**). In contrast, *Shank3*^−/−^ mice emitted fewer urine-specific USV bursts than WT mice (Mann-Whitney U-test: W = 88.5, p = 0.001) and, when recorded, they took place not only in corners, as for WT mice, but also in the middle of the cage in some cases (**Supplementary data - Specific USV types, Urination sequences**). Nevertheless, their acoustic characteristics were not significantly different from those of WT mice (**Figure 7a**). This suggests that the activation but not performance of the urine marking behavior is perturbed in *Shank3^−/−^* mice.

### Intonation and speed variations in Shank3^−/−^ female mice

*Shank3*^−/−^ mice displayed fewer intonations over entire USV bursts, compensated by a simple increase in power (**Supplementary Figure S18a**). Interestingly, long USV bursts emitted by *Shank3*^−/−^ mice during Train2 and longChase did not display the typical variations in slope or mean or end frequency observed in WT mice. However, most USV bursts were characterized by ∩-shape variations in duration, frequencyTV, mean power, frequency dynamic, number of modulations, linearity index, and harshIndex, as in WT mice **(Supplementary Figure S18b-c**). The variations in speed and duration, along with the presence/absence of USVs, were similar to those observed in WT mice (**Supplementary figure S19**).

## Discussion

### Tackling the complexity of mouse ultrasonic vocalizations

In the vast majority of previous studies investigating mouse social communication, USVs were triggered by social deprivation (*e.g.*, two weeks of isolation) and recorded in the first minutes or hours of interaction (Neunuebel et al., 2015; Sangiamo et al., 2020; Scattoni et al., 2011; Warren et al., 2018b, 2020). The USV types were either classified in a pre-determined repertoire (Scattoni et al., 2011) or simplified for modeling to reduce the complexity of the signals (e.g., ignoring harmonic components or frequency jumps or normalizing over duration; (Neunuebel et al., 2015; Sangiamo et al., 2020; Warren et al., 2018b, 2020)). In our study, we provide a complementary approach by determining a large set of acoustic variables to avoid masking the complexity of the signals and leave the door open for any user to design their own classification.

We chose not to build an exhaustive repertoire, as the link between meaning and classification can be a pitfall due to the normalization of time and frequency, which provides the same meaning to short, long, or differently dynamic USVs. We chose to examine specific call types and relate them directly to the behavioral context. We observed that USVs emitted during intense social interactions were longer, harsher, more modulated, and showed more frequency jumps than USVs emitted during non-social behaviors or at the end of social contacts (**Figures 3–4**). This is reminiscent of USVs emitted by wild house mice during social contact, which are longer than USVs emitted when standing alone (near a food spot, in the nest) (Hoier et al., 2016b).

### Hypotheses about the functions of USVs

As previously observed in captive wild house mice (von Merten et al., 2014), we confirm over a long period that males in same-sex pairs emit few spontaneous USVs. In contrast, socially-deprived males have been shown to emit a non-negligible number of USVs (e.g., (Chabout et al., 2012; Ferhat et al., 2015b; Hammerschmidt et al., 2012)). In these previous studies, the vocal repertoire did not differ significantly between males and females in this context (Hammerschmidt et al., 2012) and both sexes emitted USVs mostly during ano-genital sniffing and approach behavior (Ferhat et al., 2015b, 2016b). In the present study, males emitted USVs in short bursts with context-specific rates a hundred times lower than that of females (**Figure 2b**), probably because females were more active (longer distance travelled during the night) and spent more time in social interactions than males (**Supplementary Figure S20**; see also (de Chaumont et al., 2019)). Overall, because males are slightly less social and active than females, they may encounter fewer situations that trigger USVs. Nevertheless, this incommensurate decrease in call rate relative to females may be related to the natural social structure of mice, with only subordinate males regrouped in same-sex subgroups (Palanza et al., 2005). Other communication modalities, such as body posture and tactile contacts, may be sufficient for males to regulate their social interactions.

We observed that females emitted USVs mostly during intense social interactions, such as Train2, follow, and longChase. The speed of the mice was also significantly higher during behaviors associated with USVs than those without (**Figure 5**). It is thus possible that USVs reflect a high level of excitement, at least in C57BL/6J, the mouse strain tested here. The emission of a large number of spontaneous USVs suggests that female mice are sufficiently aroused in their “home-cage-like” life. In short-term experiments, socially-housed mice never reach this level of excitement during the first minutes of interaction and therefore emit few USVs (e.g., (Ey et al., 2018; Hammerschmidt et al., 2012)). The emission of a large quantity of USVs can only be reached by social deprivation over such short periods. Social behaviors, such as longChase, follow, or approach contact, were significantly longer when accompanied by USVs than when occurring without (**Figure 5**). This result parallels the reported increased duration of chasing when USVs were emitted by females chased by males (Neunuebel et al., 2015). It is thus possible that some USVs are emitted only when a certain level of excitement is reached, for example after a specific time spent in contact. Whether certain USVs serve to maintain social contacts is currently unknown and would need to be tested using playback experiments synchronized with behavioral monitoring.

### Subtle social communication abnormalities in Shank3 mutant mice

In our setting, *Shank3^−/−^* mice were less active and spent less time in intense social interaction, whereas individual contact events tended to be longer than those of WT mice (**Supplementary Figure S12**). These activity and social specificities are consistent with the results of previous studies using the same mutant mouse strain (de Chaumont et al., 2019; Vicidomini et al., 2016). Other genetic models of *Shank3* mutant mice also display reduced social interest (*Shank3*-KO ex4-9: (Bozdagi et al., 2010; Wang et al., 2011; Yang et al., 2012); *Shank3*-KO in ex21: (Duffney et al., 2015); *Shank3* ex4-9: (Jaramillo et al., 2016); *Shank3*-cKI: (Mei et al., 2016); *Shank3* exon 4-22 complete KO: (Wang et al., 2016); *Shank3B*: (Balaan et al., 2019); *Shank3* exon 13-16: (Dhamne et al., 2017; Fourie et al., 2018; Peca et al., 2011); *Shank3*B^+/−^: (Orefice et al., 2019; Pagani et al., 2019)). Nevertheless, other models failed to detect any atypical social interest in *Shank3* mutant mice (*Shank3*-KO ex4-9: (Drapeau et al., 2014); *Shank3-*KO ex9: (Lee et al., 2015); *Shank3*-KO ex 21: (Kouser et al., 2013); *Shank3*-KO and HZ in ex21: (Speed et al., 2015); *Shank3*-KO ex11-21 rat model: (Song et al., 2019); conditional *Shank3* exon 4-22 knockout in forebrain, striatum, and striatal D1 and D2 cells: (Bey et al., 2018); *Shank3*^+/Q321R^ and *Shank3*^Q321R/Q321R^: (Yoo et al., 2019)). Such variability may be the consequence of differences in the type of mutation, the behavioral protocols, or housing conditions.

There are few studies that have investigated ultrasonic communication in *Shank3* mutant mice. Pup isolation calls were not affected in *Shank3*-KO ex4-9 mice (Jaramillo et al., 2016; Yang et al., 2012), the *Shank3*-KO ex11-21 rat model (Song et al., 2019), or *Shank3b*-KO mice (Balaan et al., 2019), whereas those of complete *Shank3-*KO ex4-22 mice showed a lower rate, as well as shorter duration, lower frequency, and lower amplitude (Wang et al., 2016). In the context of a male briefly interacting with an estrus female, the rate of USV emission was not affected in *Shank3-*KO ex21 (Kouser et al., 2013) or *Shank3-*KO ex4-9 mice (Yang et al., 2012). Nevertheless, it was lower in *Shank3B*-KO mutant males (Dhamne et al., 2017; Pagani et al., 2019) and higher in *Shank3-*KO ex4-9 males (Jaramillo et al., 2016; Wang et al., 2011) than in WT male mice. In *Shank3*^Q321R/Q321R^ mutant mice, the number of USVs in male-female interactions was not significantly different from that of WT mice, but *Shank3*^+/Q321R^ mutants showed a higher mean duration of USVs than WT mice (Yoo et al., 2019). Male complete *Shank3-*KO ex4-22 mice also showed a lower rate of USVs than WT males when encountering an estrus female, but they were of shorter duration, reduced amplitude, and normal peak frequency (Wang et al., 2016). One strength of our study is that it was designed to directly relate communication deficits with behavioral deficits. For example, the lower call rate of Shank3^−/−^ mice on the second night may be related to fewer longChase events, a behavior that triggers a high rate of USV emission in WT mice.

Concerning USV bursts, *Shank3^−/−^* mice emitted fewer USVs per burst, but their USVs were longer and weaker than those of WT mice, with increasing power throughout the burst, a trait rarely observed in WT mice. Variations in USV duration were already shown to be present in the *Shank3* mutant carrying the Q321R point mutation (Yoo et al., 2019), while the reduction of amplitude/power was also highlighted in the *Shank3* KO ex4-22 mice (Wang et al., 2016). The markedly reduced frequency of vocalized urination sequences observed in our study may parallel the unstable dominance hierarchy in complete *Shank3* KO triads relative to WT triads (Wang et al., 2016). Overall, in our hands, Shank3^−/−^ mice were able to emit USVs with slight acoustic differences and used them in similar contexts as WT mice, but perturbed hierarchical relationships may modify the use and structure of the USVs and the intonation of USV bursts.

### Perspectives

Here, we showed sex- and age-related differences in the spontaneous communication of mice. Major quantitative and qualitative differences emerged between males and females from three months of age on. Females emitted more USVs than males, specifically during intense social investigations. With increasing age, mice emitted longer and more complex USVs, with specific differences between USV acoustic structure and behavioral contexts. Female *Shank3*^*−/−*^ mice emitted USV bursts with fewer but longer and more modulated USVs than age-matched WT females. Our system offers the possibility to characterize spontaneous mouse communication and paves the way for new studies investigating the complex interplay between genetic background, social experience, and hierarchy in the richness of social communication.

## Supporting information

Supplementary data - usv types

## Acknowledgments

The authors thank Raimund Specht from Avisoft Bioacoustics from upgrading the Avisoft Recorder software, Nathalie Lemière and Sabrina Coqueran for genotyping mice, Laetitia Breton, François Rimlinger, Didier Montéan and Christophe Joubert for animal facility support, Benoît Forget and Nicolas Torquet for critical reading of the manuscript. This work was partially funded by the Institut Pasteur, the Bettencourt-Schueller Foundation, the Cognacq– Jay Foundation, the Conny–Maeva Foundation, the ERANET–NEURON SYNPATHY program, the Agence Nationale de la Recherche through grant number ANR-10-LABX-62-IBEID, France-BioImaging infrastructure through grant number ANR-10-INBS-04 and the INCEPTION program through grant number ANR-16-CONV-0005, the Centre National de la Recherche Scientifique, the University Paris Diderot, the BioPsy Labex, and the Foundation for Medical Research (Equipe DEQ20130326488). The funders had no role in study design, data collection and analysis, decision to publish or preparation of the manuscript.

## Author contributions

E.E. designed the study, performed and analyzed experiments. F.d.C. created the segmentation and analysis methods and performed the data mining. E.E., F.d.C. and T.B. conceived the project and wrote the manuscript.

## Methods

### Animals

We tested 8 male and 8 female C57BL/6J mice (hereafter WT mice; Charles River Laboratories, Ecully, France). Mice arrived at 3 weeks of age and were directly housed in pairs in the experimental facility. *ProSAP2/Shank3* mutant mice (Schmeisser et al., 2012) were generated on site from heterozygous parents on a C57BL/6J background (>10 backcrosses). Twelve *Shank3*^−/−^ females arrived at 1-1.5 month of age to the experimental facility and were directly housed in pairs upon arrival. In the experimental facility, mice were housed in classical laboratory cages with food and water ad libitum and 11h/13h dark/light rhythm (lights off at 08:00 p.m.).

We inserted RFID chips (12 × 2.12 mm; Biolog-Id, Bernay, France) subcutaneously behind the left ear and pushed it down to the flank at 4 weeks of age in WT mice and between 5 and 7 weeks in *Shank3*^−/−^ mice. After this operation conducted under gas (isoflurane) anesthesia and local analgesia (<0.05 ml lidocaïne at 20 mg / ml), mice were left at least one week to recover.

We did not control for sexual status in our experiments. Nevertheless, as the experiments lasted 3 days, we cover a large proportion of the sexual cycle (see also (von Merten et al., 2014)). In addition, we wanted to avoid manipulating animals throughout the recording session and following estrus cycle would have necessitated daily handling.

### Behavioral and USV recordings

For the recordings, each pair was placed in the LMT setup (de Chaumont et al., 2019), a Plexiglas cage of 50 × 50 cm with transparent walls furnished with fresh bedding, food dispersed on the ground in the center, a water bottle at the down right side of the cage, a house (width: 100 mm, depth: 75 mm, height: 40 mm) in red Plexiglas in the down left corner and nesting material (6 dental cottons) spread on the bedding in the center of the cage. Recordings were launched between 03:00 and 04:00 p.m. (20-23°C, 80-100 lux when lights were on). Once the recordings were started, we did not disturb the mice anymore for 3 days (11h/13h hours dark/light rhythm, with lights off at 08:00 p.m.). Recordings were stopped after 71 hours and setups were cleaned with soap water, dried and re-furnished before launching another recording session. In WT mice, the first recordings occurred at 5 weeks of age. We next recorded in identical conditions the same mice at 3 and 7 months of age. Between the recording sessions, the pairs of mice were left undisturbed, with only a weekly change of the bedding. The six pairs of *Shank3*^−/−^ mice were recorded only once, at 2.5-3 months of age. Launching recordings consisted in monitoring both behavior and USV emission. Behavioral monitoring occurred through the LMT system (plugins 465, 524, 600 or 705; (de Chaumont et al., 2019)), in which mice were automatically identified and tracked throughout the recording session. All spontaneous ultrasonic vocalization sequences were recorded using the Avisoft UltraSoundGate Recorder system (Avisoft Bioacoustics, Berlin, Germany; 300 kHz sampling rate, 16-bit format) using the trigger function (trigger: level of this channel; pre-trigger: 1 s; hold time: 1 s; duration > 0.005 s; trigger event: 2% energy in 25-125 kHz with entropy < 50%; **Supplementary Figure S21**). Both systems were synchronized within LMT (see hereafter).

### Behavioral events

The spontaneous behavior of the mice was automatically labelled using the Live Mouse Tracker system. Both social and non-social behaviors were used, as in (de Chaumont et al., 2019) (recapitulated in **Table I**).

### Ethical statement

The protocol has been validated by the ethical committee of the Institut Pasteur (CETEA n°89) and has been conducted under the approval of the Ministère de l’Enseignement Supérieur, de la Recherche et de l’Innovation under the reference APAFIS#7706-2016112317252460 v2.

### Motivation to create a new recording and analysis pipeline

To detect and analyze USVs within background noise while monitoring the behaviors, automation was needed to handle large data sets, reduce processing time and avoid variability in human-generated errors. Existing systems (listed on the MouseTube website; (Torquet et al., 2016)) did not completely fulfill our needs. Indeed, some systems provide a classification of USV types without automated USV detection (e.g., VoiCE, (Burkett et al., 2015)). Other systems provide both automatic detection of USVs and extraction of acoustic features (A-MUD, (Zala et al., 2017); USVSEG, (Tachibana et al., 2020); Ax, (Seagraves et al., 2016)). Some even classify USVs into call types, either in a pre-determined repertoire (Mouse Song Analyzer v1.3, (Arriaga et al., 2012; Chabout et al., 2015)) or in an open repertoire determined by the data themselves (MUSE, (Neunuebel et al., 2015); MUPET, (Van Segbroeck et al., 2017); DeepSqueak, (Coffey et al., 2019)). Nevertheless, most of these systems do not handle background noise, and only MUSE provides the synchronization with behavioral monitoring, along with a heavy triangulation system that could not be easily replicated and adapted to our Live Mouse Tracker behavioral monitoring system.

### USV Segmentation Method

#### Vocalization waveform to spectrum

We process wav files recorded at a sampling rate of 300 kHz with a resolution of 16 bits per sample. We first apply an FFT (overlap of 0.75 and FFTSize = 1024 points) to the original audio signal to get its spectrum. We note here *spectrum[t][f]* each magnitude data of the spectrum.

The signal recorded presents three main problems: 1/ The continuity of the USVs can be interrupted if the signal gets too low. 2/ The animals produce a lot of noise by interacting with their environment. 3/ The animal facility environment itself produces interfering ultrasonic noise. We illustrate in the **Supplementary Figure S22** these three noise types. To overcome these problems which might prevent correct classification, we need to filter the signal of the spectrum.

#### Method parameters

We aimed to create a method with a minimum of parameters. Nevertheless, we still have one: is the vocalization emitted by an adult or a pup? Indeed, the major difference is the recording protocol. Pups are recorded at a close range, with low environmental noise. Juvenile and adult mice are recorded with the microphone placed at a distance to cover the surface of the test cage. Also, as they move freely, noise generated by their interactions with the environment is important. Therefore, as the signal to noise ratio is better in pup recordings than in adult conditions, we lower our detection threshold for pups. In addition, as the range of emission of pups rises up to 140 kHz, we increase the maximum frequency allowed for detection.

#### Filtering spectrum data

For each step, we provide pseudo code. For an easier reading, we removed the boundary check conditions that exists in the real code.

For each time point of the spectrum, we center the data by removing the mean of all magnitude for all frequencies at the current time point.

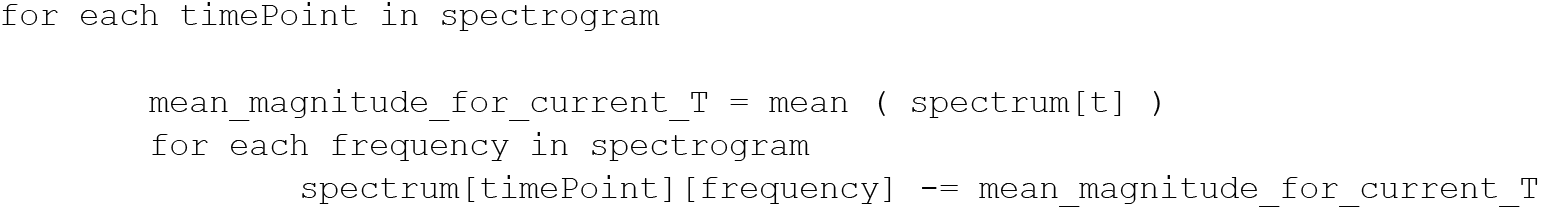

We then use a filter that creates continuity in signal. It basically fills the gap in the signal if a direction is found in the spectrum’s curves. To perform this filtering, we first create a filter bank for a number of angles. The following code pre-computes the filters for the different orientations:

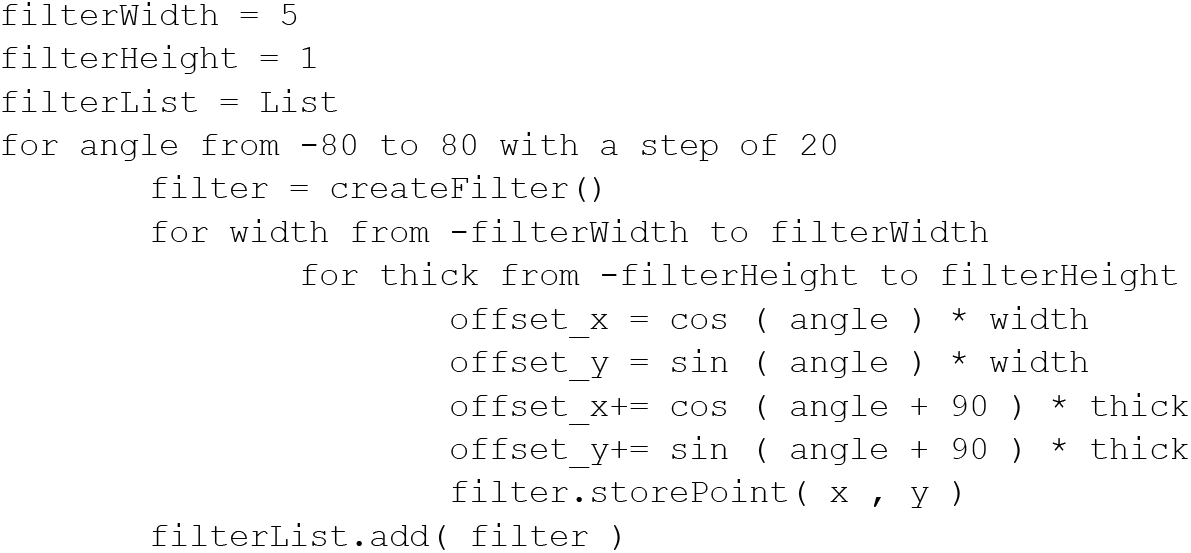

Then, we apply those pre-computed filters on each point of the spectrogram, and we keep the maximum response. This code is parallelized for each time point for maximum performance.

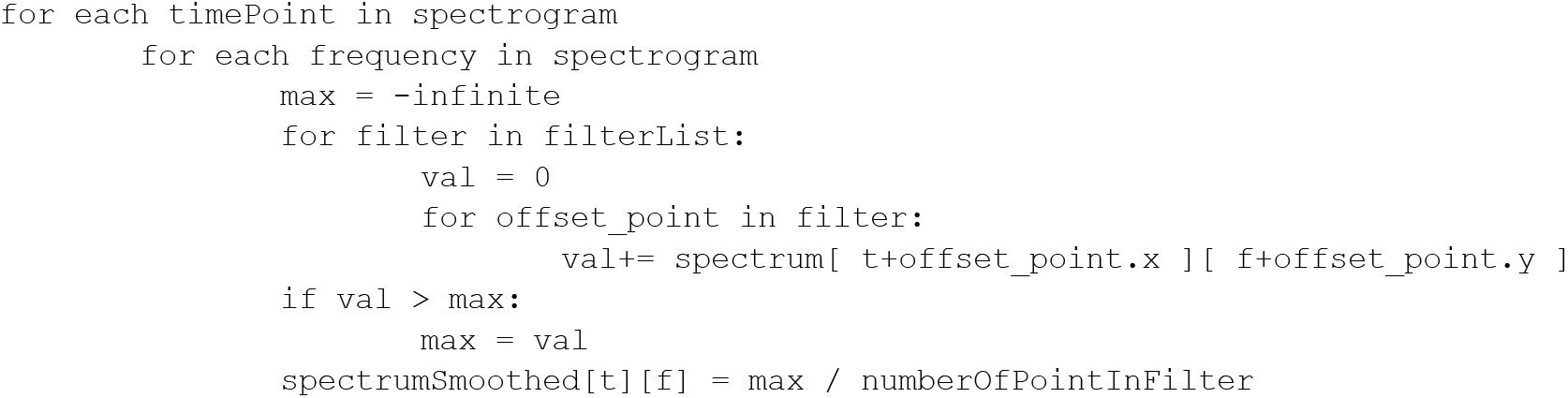

We then filter the vertical signal to remove noise (seen as strong vertical scratches in the spectrum). The following pseudo code removes for each frequency the local mean frequency of the spectrum, using a sliding window of +/− 1.5 kHz.

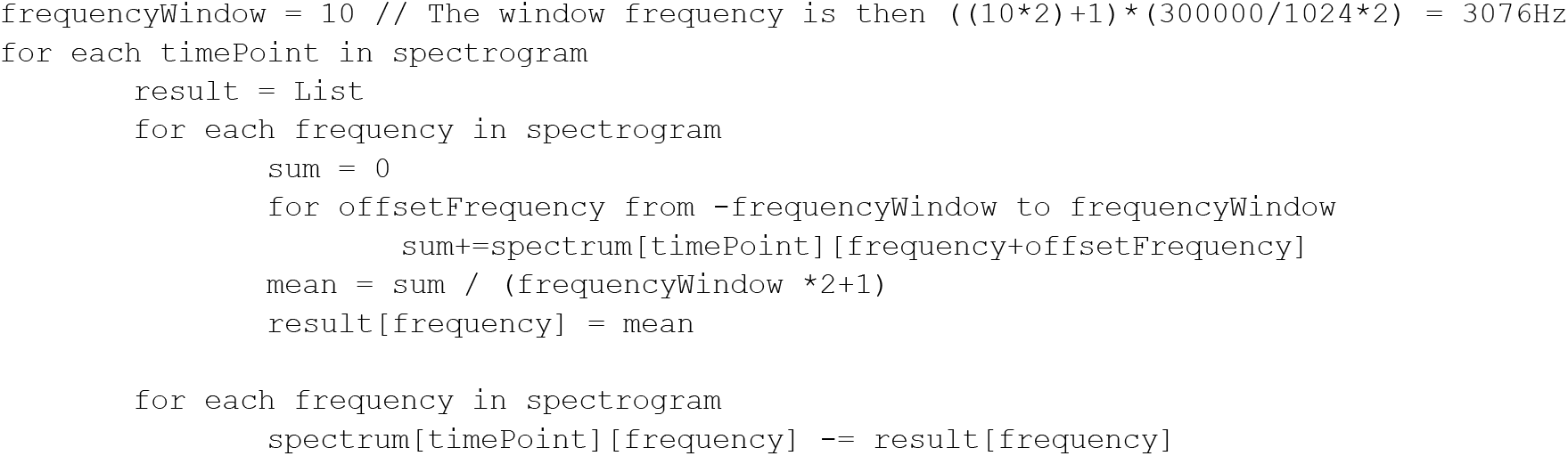

#### Constant and blinking frequency canceler

In the experiments, we observed noise due to light, fan, power sources and air-conditioning (**Supplementary Figure S22**). They appear and disappear randomly during experiment, at unpredictable frequencies, and can switch in frequency. We process the spectrum to find the frequencies of those noise to store them in a “frequency cancelation list”.

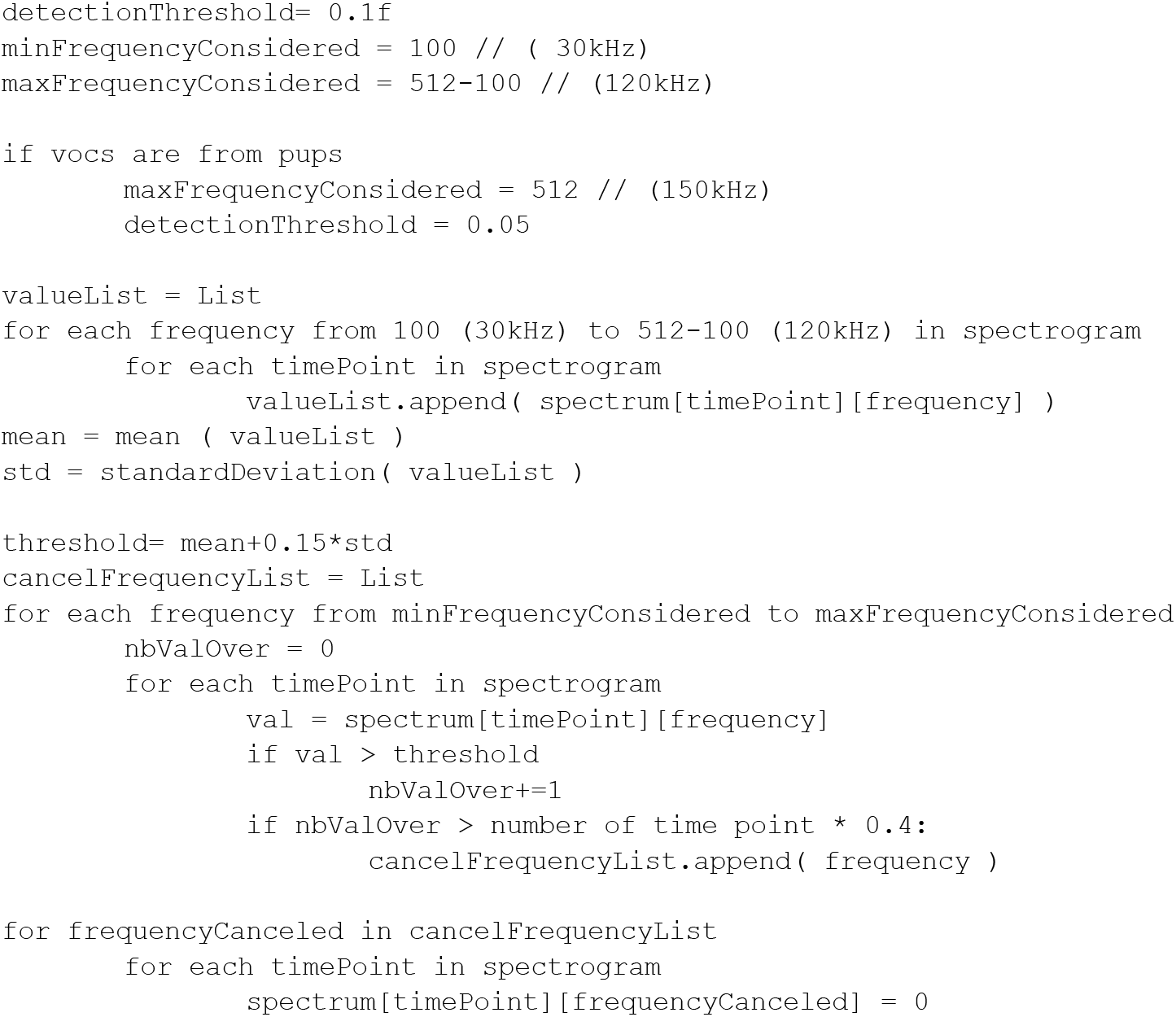

#### Voc segmentation

On the filtered signal, we now consider all values over 0 in spectrum.

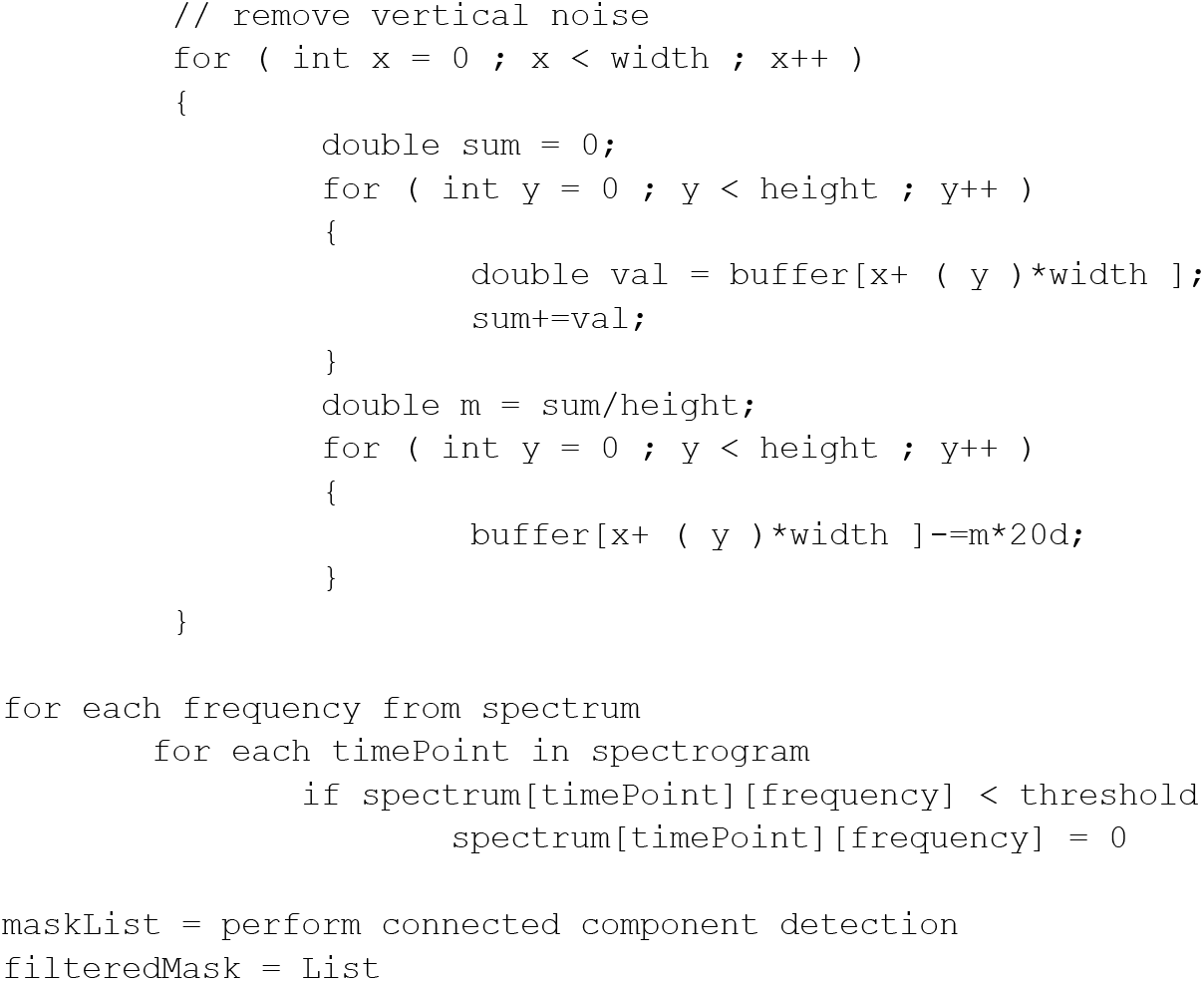

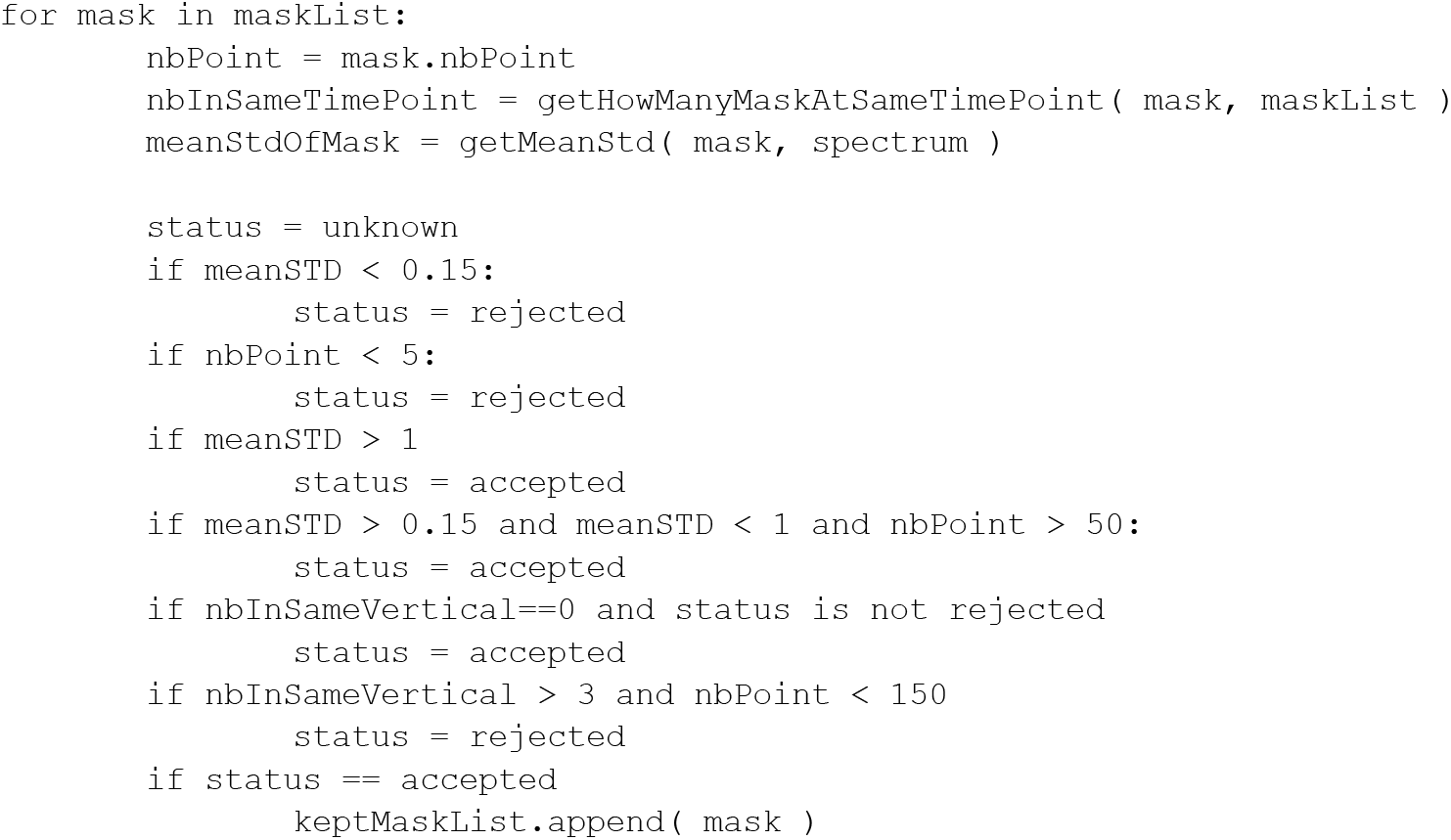

Then masks are merged together if they are sharing a time point. They are then temporally merged again if the silence between the signals is below 40 ms.

#### Spectrum signal extraction

The final extraction of the signal is the maximum magnitude per time point that belongs to the mask of the time point (if available).

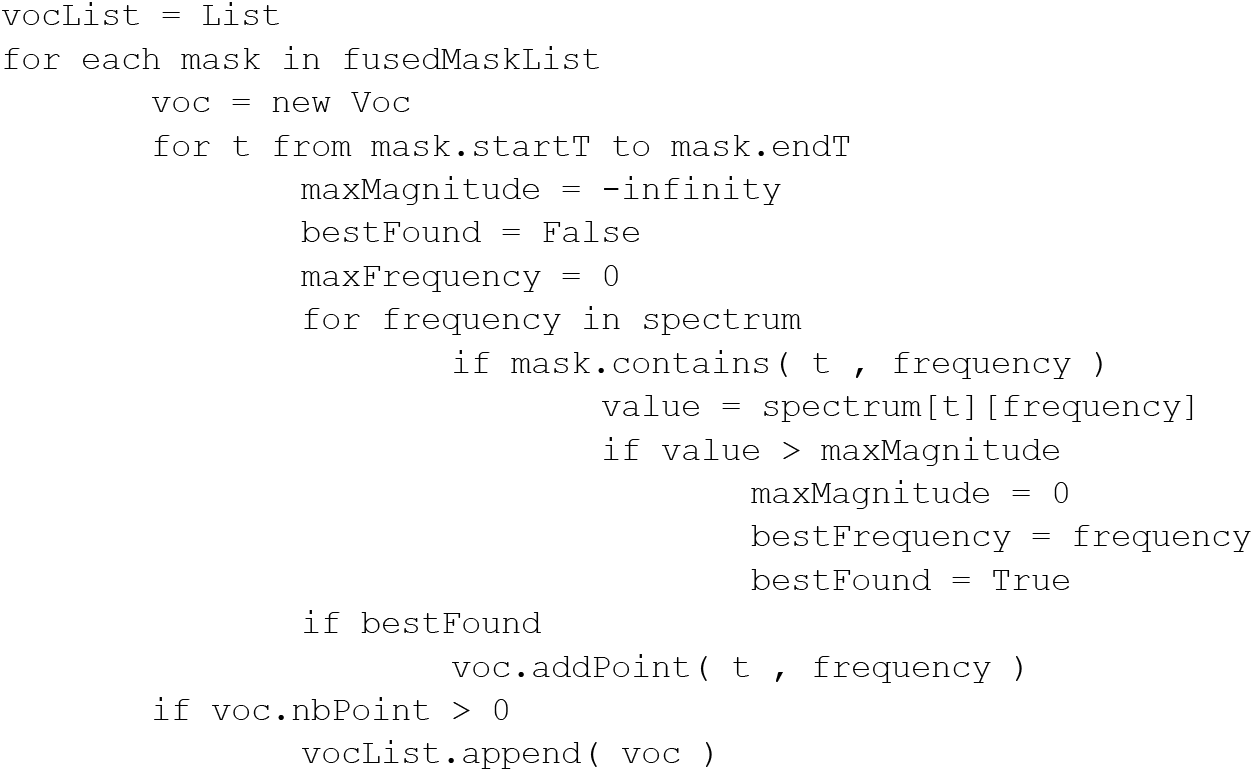

#### Validation

To perform the validation, we consider the beginning and the end of each USV. We run our segmentation algorithm on a set of manually annotated USVs (10 files for each experiment; **Supplementary Figure S23**). If the USV boundaries match at +/− 40 ms, we consider that the USV has been correctly detected (**Table II**).

**Table II:**
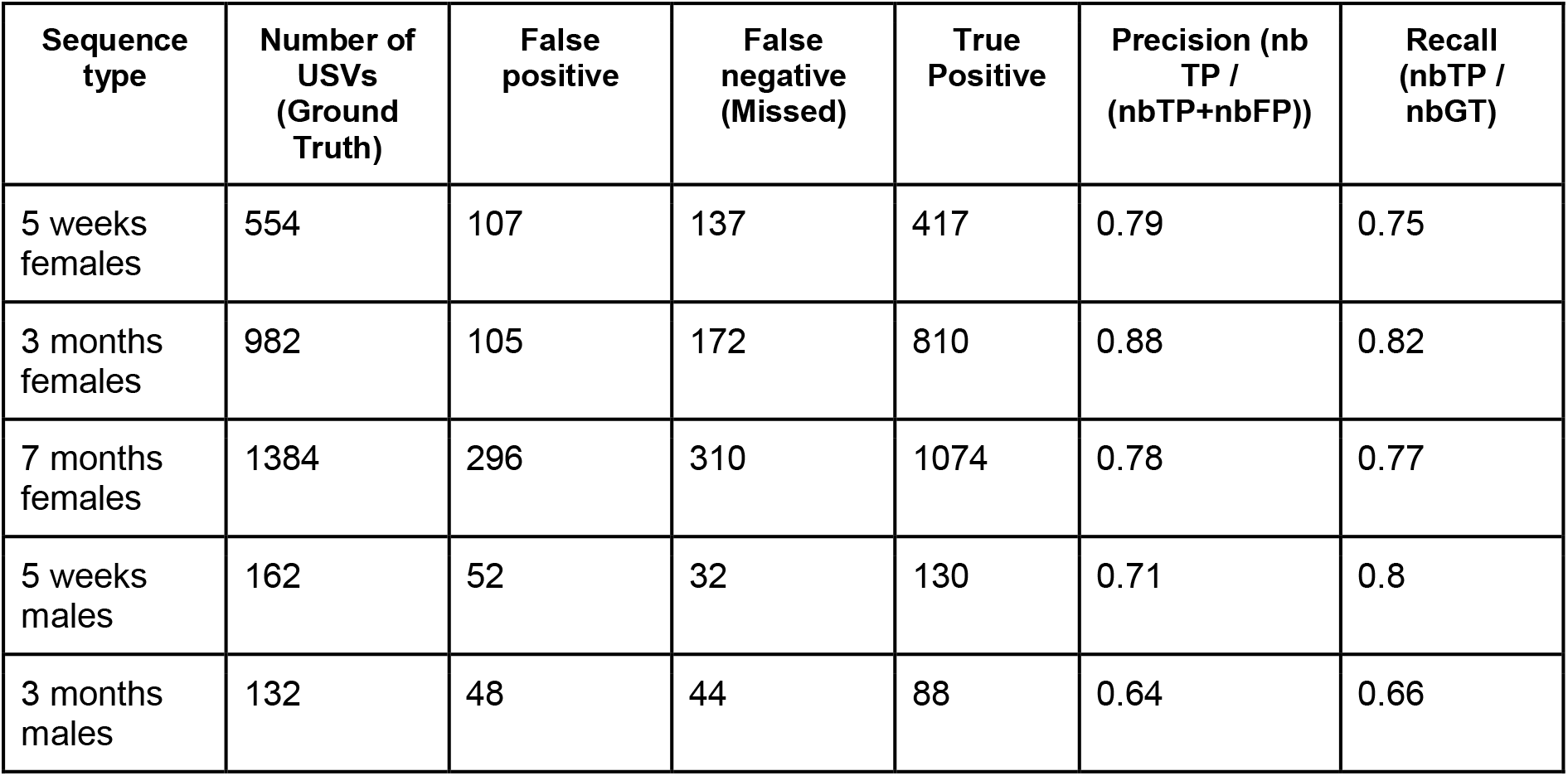
Validation scores by comparison with manual annotation of USVs.

Nevertheless, this metric is based on the start and end time of the sequences, which raises two concerns:

- After the processing, we checked again the ground truth. We found that most discrepancies between the ground truth and the automatic segmentation emerge from the merging of two parts of USV connected by a low-power signal in the manual annotation, while the two parts were considered separately by the automatic segmentation. In that case, the penalty is very high, as this leads to two false positives and one false negative.
- The second concern is that our metric does not check if we segment correctly the peak frequency itself. This would have required the development of a dedicated annotation tool. Nevertheless, we believe that such method should be introduced in our further developments.

#### Acoustic feature produced

The **Table III** and **Table IV** provide the description for the acoustic traits computed for each USV and each USV burst.

**Table III:**
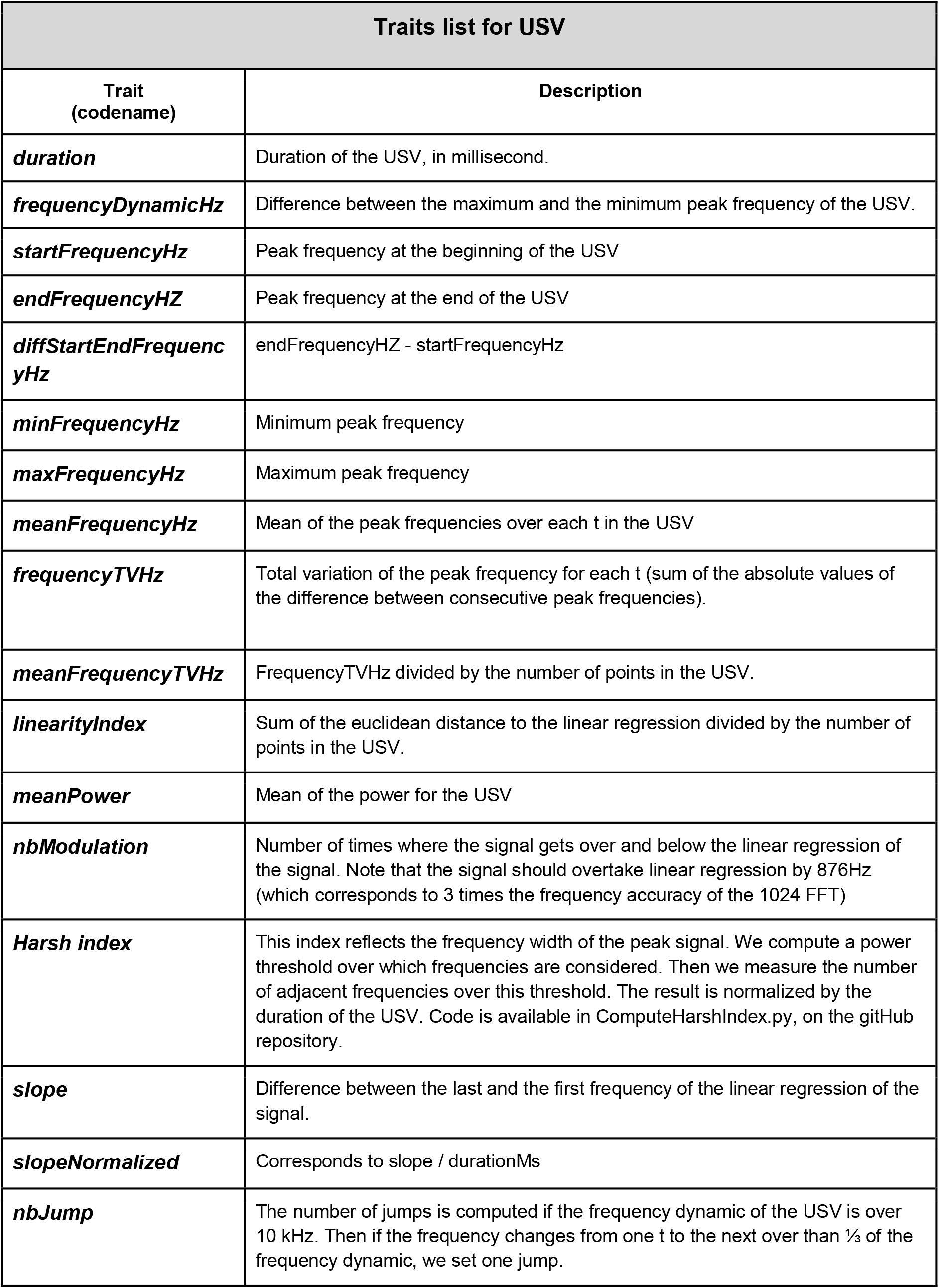
Description of the acoustic variables measured for each USV with the system.

| Traits list for USV |  |
| --- | --- |
| Trait<br>(codename) | Description |
| ***duration*** | Duration of the USV, in millisecond. |
| ***frequencyDynamicHz*** | Difference between the maximum and the minimum peak frequency of the USV. |
| ***startFrequencyHz*** | Peak frequency at the beginning of the USV |
| ***endFrequencyHZ*** | Peak frequency at the end of the USV |
| ***diffStartEndFrequencyHz*** | $\text{endFrequencyHZ} - \text{startFrequencyHz}$ |
| ***minFrequencyHz*** | Minimum peak frequency |
| ***maxFrequencyHz*** | Maximum peak frequency |
| ***meanFrequencyHz*** | Mean of the peak frequencies over each t in the USV |
| ***frequencyTVHz*** | Total variation of the peak frequency for each t (sum of the absolute values of the difference between consecutive peak frequencies). |
| ***meanFrequencyTVHz*** | FrequencyTVHz divided by the number of points in the USV. |
| ***linearityIndex*** | Sum of the euclidean distance to the linear regression divided by the number of points in the USV. |
| ***meanPower*** | Mean of the power for the USV |
| ***nbModulation*** | Number of times where the signal gets over and below the linear regression of the signal. Note that the signal should overtake linear regression by 876Hz (which corresponds to 3 times the frequency accuracy of the 1024 FFT) |
| ***Harsh index*** | This index reflects the frequency width of the peak signal. We compute a power threshold over which frequencies are considered. Then we measure the number of adjacent frequencies over this threshold. The result is normalized by the duration of the USV. Code is available in ComputeHarshIndex.py, on the gitHub repository. |
| ***slope*** | Difference between the last and the first frequency of the linear regression of the signal. |
| ***slopeNormalized*** | Corresponds to $\text{slope} / \text{durationMs}$ |
| ***nbJump*** | The number of jumps is computed if the frequency dynamic of the USV is over 10 kHz. Then if the frequency changes from one t to the next over than $\frac{1}{3}$ of the frequency dynamic, we set one jump. |

**Table IV:** Description of the characteristics for each USV burst with the system.

| Traits list for USV Burst |  |
| --- | --- |
| Trait | Description |
| *nbUSV* | Number of USVs contained in the burst |
| *durationMs* | Duration of the burst, in millisecond. |
| *meanDuration* | Mean duration of USVs contained in the burst |
| *stdDuration* | Standard deviation of the durations of USVs contained in the burst |
| *meanInterval* | Mean interval (silence duration) between the USVs |
| *meanFrequency* | Mean of mean peak frequency of USVs contained in the burst |
| *mean power* | Mean power of USVs contained in the burst |

#### Filtering out wave files containing only noise

During an experiment, we do not record continuously the audio information. We use Avisoft-RECORDER automatic triggering capability which monitors the audio and starts the recording when a predetermined power threshold is reached. Therefore, the dataset of an experiment is composed of thousands of files that may contain USVs or just noise due to the activity within the cage. Therefore, data of an experiment can be preprocessed to filter out wave files containing only noise, which represent 50% of the files generated in our experiment.

An expert sorted files containing USVs from files containing noise to train a random forest classifier. We use the following features as machine learning features: for each file, we extract the mean power of the whole file, the number of presumed USVs detected, the duration of the USVs over the overall length of the file, the average duration of USV and its standard deviation, the mean peak frequency and the standard deviation of the peak frequency.

We then train the random forest classifier. We provided a pre-trained classifier but one can re-train the classifier with its own data. One just needs to start the python script and point two different folders (noise and USVs) to train the system. For our experiments, we trained the system with 451 files containing USVs and 247 with only noise. The accuracy of the training is 97%, using 10 folds.

Then, the system can be used in predictive mode to sort dataset containing both USVs and noise. The script copies the file in a noise and USV folder, so that the user can easily control the sorting accuracy.

#### Avisoft burst record and synchronization with Live Mouse Tracker

The system is designed to work for an unlimited duration. As USVs are infrequent events, we do not record the sound continuously. We instead use the automatic record trigger functionality of Avisoft-RECORDER (**Supplementary Figure S21**). The automatic trigger of Avisoft-RECORDER monitors the sound level and start recording a sound if the current sound level within a given frequency range is over a given threshold. The sound is recorded as long as the sound level is over the threshold. This function takes a hold time parameter: if Avisoft-RECORDER detects another signal during the hold period, the record is not interrupted. The hold period also adds a record period around the first and the last signal over threshold. In our experiment, we use a hold time of one second.

To synchronize USV recording with the tracking, we use the “Trigger control” of Avisoft-RECORDER. This function allows to launch an external program at each start and end of records. We use the free software PacketSender (https://packetsender.com/) to perform communication between Avisoft-RECORDER and Live Mouse Tracker (LMT). Through PacketSender, we send an UDP string packet containing the file number currently recorded by Avisoft-RECORDER. This information is recorded by LMT within the database as an “USV event”. The goal of the synchronization is to match the USV record with the current data frame number recorded by LMT.

### USV Toolbox, an open-source, free and online USV analysis pipeline

The currently available methods to detect and analyze mouse USVs need specific installations and software. To facilitate the testing of our own algorithm, we provide a website to test the method or to process data online: https://usv.pasteur.cloud. The user simply drags and drops his/her wave file, waits a few seconds (depending on the length of the sample file) and finally evaluates the quality of the USV segmentation and the data extracted from the sound file. The goal of this website is to provide immediate access to the method without installing any software.

The first panel of the website is dedicated to evaluate USV detection. The first spectrogram represents the original data and the second one provides the annotated data. The player under this spectrogram allows to listen to the sound file slowed down by twenty times. The other panels display:

- the length of the wave file given as input.
- the number of USVs detected within the wave file.
- a timeline displaying the USVs detected over the whole file and their temporal organization in USV bursts, in which the intervals between USV are shorter than one second.
- the frequency characteristics of each USV within the sound file (in kHz). Each vertical black bar displays the min/max peak frequency of the USV (and therefore also the frequency range) while the black dot displays the mean peak frequency and the red dot displays the peak frequency with the maximum amplitude in each USV.
- the duration of each USV (in ms).
- the power (i.e., amplitude) of each USV, depicted in arbitrary unit.
- the proportion of USVs with frequency modulations.
- the proportion of USVs containing one or more frequency jump(s).
- a table gathering all acoustic variables extracted on each USV of the sound file.

The user can download all these results for his/her own sound file. These results are deleted after one hour. Data downloaded from this web page can be directly used with the scripts that we provide with the present study. To perform the analysis on thousands of files, we also provide the desktop version of the analysis program, working in batch mode (link available on https://usv.pasteur.cloud after publication).

### USV analysis toolbox

As for Live Mouse Tracker, we provide an API in Python for the biologists to process USVs. This package allows one to re-create all data representations used in this study with its own data. This API is available on gitHub (will be released after publication process).

For Live Mouse Tracker, we provided a full API in Python to process event classification, and to process queries.

#### Analyses and statistical tests

#### Behavioral profiles of mice

We built a behavioral profile for each mouse considering the following behavioral event types: stop isolated, move isolated, huddling, rear isolated, wallJump, water Stop, break contact, getaway, social approach, approach contact, approach rear, stop in contact, move in contact, rear in contact, contact, oral-oral contact, oral-genital contact, side by side contact, side by side contact / opposite way, follow, train2, longChase, USV burst. We also examined the total distance travelled. For each behavioral event type, we compared the total time spent in this behavioral event type, the number of events, and the mean duration of these events.

Each of these variables were tested for the effect of age within each sex separately. We used paired Wilcoxon tests to compare age classes within each sex. The effect of sex was tested within each age class separately using Mann-Whitney U-tests. Variables for *Shank3*^−/−^ mice were compared to variables for WT mice aged of 3 months using Mann-Whitney U-tests. Data were corrected for multiple testing whenever necessary.

#### Comparison between age classes, sexes or genotypes of USVs and USV bursts

To examine the effect of age class on USVs and on USV bursts, we used non parametric paired Wilcoxon tests to compare the amount of USVs, of USV bursts and of USVs per USV burst between age classes within each sex.

To detect a sex effect on the number of USVs per USV burst, we used non-parametric unpaired Mann-Whitney U-tests to compare the number of USVs per USV burst between sexes within each age class (5 weeks, 3 months and 7 months). We compared the amount of USVs and USV bursts as well as the number of USVs per USV bursts between *Shank3*^−/−^ and WT mice using non parametric Mann-Whitney U-tests for each night separately.

Acoustic features of USV bursts were compared between *Shank3*^−/−^ and WT mice using non parametric Mann-Whitney U-tests in each behavioral context separately corrected for the number of variables tested (burst duration, number of USVs, mean duration of USVs, standard deviation of the duration of USVs, mean duration of intervals between USVs, standard deviation of the duration of intervals between USVs, mean peak frequency of USVs), as well as the number of behavioral contexts examined (approach contact, approach rear, break contact, follow, oral-genital contact, oral-oral contact, side-by-side contact, side-by-side contact opposite way, stop isolated, train2, urination, longChase).

#### Temporal organization between USVs and behaviors

We tested whether specific behavioral event types occurred significantly more before or after USVs. For this purpose, we focused on the following behavioral events: stop isolated, move isolated, break contact, getaway, social approach, approach rear, approach contact, contact, oral-oral contact, oral-genital contact, side-by-side contact, side-by-side contact opposite way, sequence oral-oral / oral-genital, sequence oral-genital / oral-oral, follow, train2, and longChase.

For each event timeline, we created two counters (pre and after) representing the number of times the USV is after or before an event. For each USV, we compute the pre and after interval, which are respectively [start of USV - 1 second, start of USV] and [end of USV, end of USV + 1 second]. We seek for the presence of an event in those two intervals by counting the number of frames of the event overlapping those intervals. We increment pre or after depending on this result (the interval with the maximum number of overlapping frame with the events wins, if this is a draw, no increment occurs). Once the two counters representing the presence of events before and after the USVs are obtained, we use the central limit theorem to test if the events are significantly before or after the USVs. We therefore computed p = after / (pre+after). We perform the correction alpha = (sqrt(p * (1-p)) / sqrt(n)) * coef with coef equals to 3.29, 3.92, 5.15 for 90%, 95%, 99% confidence respectively. Then corrected p = p - alpha. if p > 0.5 then the events were significantly more frequent after the given event type. We performed the opposite test to check if events were significantly more frequent before an event type.

#### Context-specific acoustic features

We tested whether acoustic features of USVs (duration, frequency characteristics, frequency range, modulations, harshness, slope) varied according to the contexts in which USVs were emitted. For that purpose, after computing the acoustic features of all USVs, we compared acoustic features of USVs between the different behavioral contexts (stop isolated, break contact, approach rear, approach contact, oral-oral contact, oral-genital contact, side-by-side contact, side-by-side contact / opposite way, follow, train2, longChase, urinate) using Mann-Whitney U-tests with a Bonferroni correction (combination of 2 within 12 behavioral events × 16 acoustic features). Acoustic variations between contexts in *Shank3*^−/−^ mice were tested similarly and compared to the ones presented in WT mice using Mann-Whitney U-tests (Figure M1c-d). Heatmaps represented significance levels after Bonferroni correction as well as the direction of variations.

#### Relationship between the occurrence of USV bursts and the speed of mice as well as the duration of social events

We focused on the following sample of behavioral events: break contact, getaway, approach contact, contact, oral-genital contact, follow, longChase and Train2. First, we tested whether animals displaying these behaviors accompanied by USVs had a higher speed than when they displayed the same behaviors without USVs. For that purpose, for each behavioral event of each type, we computed the mean speed of the mouse displaying this behavior over the whole event. We performed a paired Wilcoxon test (alternative = “greater”) to compare the mean speed of the animal with and without USVs at the group level. In addition, to better understand inter-individual variations, we compared, within each individual and for each behavioral event type, the speeds during events overlapping with USVs to the speeds during events not overlapping with USVs using Mann-Whitney U-tests (unpaired, one-sided).

Second, we tested whether one type of behavioral event was longer when emitted concomitantly with USVs than when it was not concomitant with USVs. For that purpose, within each individual and for each behavioral event type, we gathered the durations of events overlapping with USVs as well as the durations of events not overlapping with USVs. We compared these two lists of durations using Mann-Whitney U-tests (unpaired, one-sided) within each individual. We also performed a paired Wilcoxon test (alternative = “greater”) to compare at the group level the mean duration of the event with USVs and the mean duration of the same event without USVs. To control for the fact that longer behavioral events were more likely to include USVs, we used a random process in which USVs would occur at a random time. In this case, the longer USVs would have more chances to overlap with an USV. We simulated 1000 different random distributions of the USVs, and we kept the same proportions of USVs overlapping with behavioral events as in the original data. We performed this test for each behavioral event in each experiment separately (**Supplementary Tables II-V**). In each simulation, we performed the same statistical test (Mann-Whitney U-test) as in the original data, and we kept the p-value. The ratio of the number of p-values below the original one provided the probability to obtain the same effect randomly. In our study, this probability was less than 3%, suggesting that the effects of the presence of USVs on the duration of behavioral events could be found randomly with only a weak probability, i.e., longer events were not more likely to occur with USVs than shorter ones.

##### simulation code: Compute_Duration_Events_With_USV_Simulation.py

We conducted these tests for both animals of each WT female pair at 5 weeks, 3 months and 7 months of age as well as for both animals of *Shank3*−/− pairs aged of 3 months for all behavioral events above cited. Overall, we adjusted our statistical analyses for multiple testing accordingly (Bonferroni correction).

#### Intonation within USV bursts

For each burst that contains a minimum of 8 USVs, we split in two halves the first USVs and the last USVs. Then for each half and for each acoustic feature, we computed if the acoustic feature was increasing or decreasing. When the acoustic feature increased and then decreased, the variations were in ∩-shape. If the feature was decreasing first and then increasing, the variations were in U-shape. If the feature was increasing or decreasing, in both halves, it was qualified as //-shape or \\-shape, respectively. The feature could also be stable (marked as 0) over one or two half(ves) of the burst. **Figure 6a** displays the proportion of occurrences of the different possibilities of intonation within long USV bursts in WT mice aged of 5 weeks, 3 months and 7 months of age. **Figure 6b** displays the frequency of occurrence of the 20 most frequent USV burst styles with one acoustic trait according to the age of female WT mice.

