## Supplementary data - usv types for "Spontaneous social communication in laboratory mice - placing ultrasonic vocalizations in their behavioral context"

### Specific USV types

#### Isolated vocalizations with high peak frequency

We recorded isolated USVs displaying a high peak frequency and covering a large frequency range (**Supplementary Figure Isolated USV-1**). These USVs occurred when a mouse was jumping down the wall or down the plexiglas house.

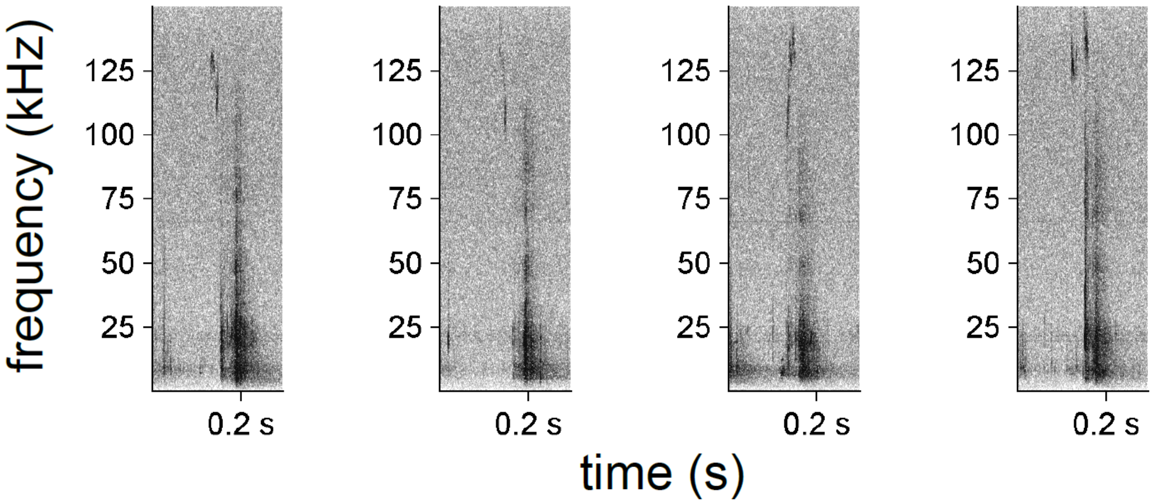


**Supplementary Figure Isolated USV-1.** Examples of spectrograms of USVs emitted during jumping against the wall or down from the house of WT female pairs aged three months (FFT-length: 1024 points, 16-bit format, 300 kHz sampling frequency, 75% overlap).

#### Urination sequences

In the present study, we observed a specific behaviour co-occurring with USV emission. During these episodes of interest, one mouse was acting in an unspecified way (e.g., in the nest, foraging, resting, moving). In contrast, the other mouse was acting specifically. It was first sniffing the bedding in a corner, then turning around to place its ano-genital region in the corner, and lifting the tail. At that stage, this mouse emitted a USV burst and stayed immobile with the tail lifted. This attitude is typical for a mouse urinating in the corner, with a large quantity of urine deposited. Indeed, females deposit small quantities of urine while moving but they stop to deposit large quantities (Coquelin, 1992). After that, most of the time, the urinating mouse turned around and sniffed the place where the ano-genital region was located (turnaround). These turnarounds occurred frequently, especially when urination sequences were numerous (see proportions in **Supplementary Tables Urination-I and II**). These urination events rarely triggered any approach behaviour in the other mouse (**Supplementary Tables Urination-I and II**). Several of these sequences are available in **Supplementary videos – Urination sequences**.

This behaviour was displayed by both sexes at any age (**Supplementary Tables I and II**). Nevertheless, it was more frequent in WT females than in WT males in all age classes (Wilcoxon rank sum test: 5 weeks: W=14, p=0.031; 3 months: W=3, p=0.001; 7 months:

W=5.5, p=0.002). This behaviour was much less frequent in WT than in *Shank3*$^{-/-}$ mice (Wilcoxon rank sum test: W=88.5, p=0.001; **Supplementary Table II**).

**Supplementary Table Urination-I**: Occurrences of urination sequences in pairs of WT males.

| category | sex | pair | ind | total number of urinate USV in pair | total number of urinate USV | proportion of all urinate USV | number of turnarounds | proportion of urinate USV with turnaround | number of occurrences of approach by the other mouse | proportion of urinate USV with approach |
|---|---|---|---|---|---|---|---|---|---|---|
| WT 5 we | male | M40_M41 | 4612467 | 15 | 0 | 0.00 | - | - | - | - |
| WT 5 we | male | M40_M41 | 4612582 | 15 | 15 | 1.00 | 5 | 0.33 | 6 | 0.40 |
| WT 5 we | male | M42_M43 | 4612572 | 3 | 0 | 0.00 | - | - | - | - |
| WT 5 we | male | M42_M43 | 4612229 | 3 | 3 | 1.00 | 2 | 0.67 | 0 | 0.00 |
| WT 5 we | male | M44_M45 | 4612506 | 7 | 2 | 0.29 | 2 | 1.00 | 0 | 0.00 |
| WT 5 we | male | M44_M45 | 4612326 | 7 | 5 | 0.71 | 4 | 0.80 | 0 | 0.00 |
| WT 5 we | male | M46_M47 | 4612368 | 0 | 0 | - | - | - | - | - |
| WT 5 we | male | M46_M47 | 4612549 | 0 | 0 | - | - | - | - | - |
| WT 3 mo | male | M40_M41 | 4612467 | 11 | 0 | 0.00 | - | - | - | - |
| WT 3 mo | male | M40_M41 | 4612582 | 11 | 11 | 1.00 | 9 | 0.82 | 4 | 0.36 |
| WT 3 mo | male | M42_M43 | 4612572 | 0 | 0 | - | - | - | - | - |
| WT 3 mo | male | M42_M43 | 4612229 | 0 | 0 | - | - | - | - | - |
| WT 3 mo | male | M44_M45 | 4612506 | 0 | 0 | - | - | - | - | - |
| WT 3 mo | male | M44_M45 | 4612326 | 0 | 0 | - | - | - | - | - |
| WT 3 mo | male | M46_M47 | 4612368 | 1 | 1 | 1.00 | 0 | 0.00 | 1 | 1.00 |
| WT 3 mo | male | M46_M47 | 4612549 | 1 | 0 | 0.00 | - | - | - | - |
| WT 7 mo | male | M40_M41 | 4612467 | 7 | 0 | 0.00 | - | - | - | - |
| WT 7 mo | male | M40_M41 | 4612582 | 7 | 7 | 1.00 | 4 | 0.57 | 1 | 0.14 |
| WT 7 mo | male | M42_M43 | 4612572 | 0 | 0 | - | - | - | - | - |
| WT 7 mo | male | M42_M43 | 4612229 | 0 | 0 | - | - | - | - | - |
| WT 7 mo | male | M44_M45 | 4612506 | 0 | 0 | - | - | - | - | - |
| WT 7 mo | male | M44_M45 | 4612326 | 0 | 0 | - | - | - | - | - |
| WT 7 mo | male | M46_M47 | 4612368 | 0 | 0 | - | - | - | - | - |
| WT 7 mo | male | M46_M47 | 4612549 | 0 | 0 | - | - | - | - | - |

**Supplementary Table Urination-II**: Occurrences of urination sequences in pairs of females for WT and *Shank3-/-* mice.

| category | sex | pair | ind | total number of urinate USV in pair | total number of urinate USV | proportion of all urinate USV | number of turnarounds | proportion of urinate USV with turnaround | number of occurrences of approach by the other mouse | proportion of urinate USV with approach |
|---|---|---|---|---|---|---|---|---|---|---|
| WT 5 we | female | F13_F14 | 4612190 | 40 | 21 | 0.53 | 8 | 0.38 | 0 | 0.00 |
| WT 5 we | female | F13_F14 | 4612361 | 40 | 19 | 0.48 | 13 | 0.68 | 0 | 0.00 |
| WT 5 we | female | F15_F16 | 4612578 | 18 | 18 | 1.00 | 6 | 0.33 | 2 | 0.11 |
| WT 5 we | female | F15_F16 | 4612626 | 18 | 0 | 0.00 | - | - | - | NA |
| WT 5 we | female | F17_F18 | 4612400 | 9 | 3 | 0.33 | 2 | 0.67 | 0 | 0.00 |
| WT 5 we | female | F17_F18 | 4612605 | 9 | 6 | 0.67 | 2 | 0.33 | 1 | 0.17 |
| WT 5 we | female | F19_F20 | 4612241 | 16 | 15 | 0.94 | 6 | 0.40 | 1 | 0.07 |
| WT 5 we | female | F19_F20 | 4612303 | 16 | 1 | 0.06 | 1 | 1.00 | 0 | 0.00 |
| WT 3 mo | female | F13_F14 | 4612190 | 27 | 15 | 0.56 | 10 | 0.67 | 2 | 0.13 |
| WT 3 mo | female | F13_F14 | 4612361 | 27 | 12 | 0.44 | 8 | 0.67 | 4 | 0.33 |
| WT 3 mo | female | F15_F16 | 4612578 | 14 | 13 | 0.93 | 12 | 0.92 | 5 | 0.38 |
| WT 3 mo | female | F15_F16 | 4612626 | 14 | 1 | 0.07 | 0 | 0.00 | 0 | 0.00 |
| WT 3 mo | female | F17_F18 | 4612400 | 21 | 10 | 0.48 | 9 | 0.90 | 4 | 0.40 |
| WT 3 mo | female | F17_F18 | 4612605 | 21 | 11 | 0.52 | 9 | 0.82 | 5 | 0.45 |
| WT 3 mo | female | F19_F20 | 4612241 | 33 | 20 | 0.61 | 19 | 0.95 | 2 | 0.10 |
| WT 3 mo | female | F19_F20 | 4612303 | 33 | 13 | 0.39 | 7 | 0.54 | 3 | 0.23 |
| WT 7 mo | female | F13_F14 | 4612190 | 25 | 22 | 0.88 | 15 | 0.68 | 2 | 0.09 |
| WT 7 mo | female | F13_F14 | 4612361 | 25 | 3 | 0.12 | 1 | 0.33 | 2 | 0.67 |
| WT 7 mo | female | F15_F16 | 4612578 | 9 | 9 | 1.00 | 7 | 0.78 | 2 | 0.22 |
| WT 7 mo | female | F15_F16 | 4612626 | 9 | 0 | 0.00 | - | - | - | - |
| WT 7 mo | female | F17_F18 | 4612400 | 35 | 17 | 0.49 | 16 | 0.94 | 6 | 0.35 |
| WT 7 mo | female | F17_F18 | 4612605 | 35 | 18 | 0.51 | 16 | 0.89 | 5 | 0.28 |
| WT 7 mo | female | F19_F20 | 4612241 | 30 | 14 | 0.47 | 13 | 0.93 | 2 | 0.14 |
| WT 7 mo | female | F19_F20 | 4612303 | 30 | 16 | 0.53 | 12 | 0.75 | 1 | 0.06 |
| Shank3 3mo | female | 4724356_4724113 | 4724356 | 3 | 0 | 0.00 | - | - | - | - |
| Shank3 3mo | female | 4724356_4724113 | 4724113 | 3 | 3 | 1.00 | 3 | 1.00 | 1 | 0.33 |
| Shank3 3mo | female | 4849069_4849197 | 4849069 | 3 | 3 | 1.00 | 0 | 0.00 | 0 | 0.00 |
| Shank3 3mo | female | 4849069_4849197 | 4849197 | 3 | 0 | 0.00 | - | - | - | - |
| Shank3 3mo | female | 4849144_4849294 | 4849294 | 3 | 2 | 0.67 | 0 | 0.00 | 0 | 0.00 |
| Shank3 3mo | female | 4849144_4849294 | 4849144 | 3 | 1 | 0.33 | 1 | 1.00 | 0 | 0.00 |
| Shank3 3mo | female | 4849155_4849457 | 4849155 | 11 | 9 | 0.82 | 9 | 1.00 | 1 | 0.11 |
| Shank3 3mo | female | 4849155_4849457 | 4849457 | 11 | 2 | 0.18 | 2 | 1.00 | 1 | 0.50 |
| Shank3 3mo | female | 4849484_4849378 | 4849378 | 13 | 4 | 0.31 | 1 | 0.25 | 2 | 0.50 |
| Shank3 3mo | female | 4849484_4849378 | 4849484 | 13 | 9 | 0.69 | 7 | 0.78 | 2 | 0.22 |
| Shank3 3mo | female | 4417496_4419810 | 4417496 | 0 | 0 | - | - | - | - | - |
| Shank3 3mo | female | 4417496_4419810 | 4419810 | 0 | 0 | - | - | - | - | - |

These behavioural sequences were accompanied by characteristic USV bursts (see example spectrograms in **Supplementary Figure Urination-1** for WT mice and **Supplementary Figure Urination-2** for *Shank3*$^{-/-}$ mice). These USV bursts were characterised by short duration (in time and in the number of USVs included within the burst), reduced variability in the duration of the USVs composing the burst and higher mean peak frequency compared to USV bursts emitted in all other contexts. USVs within these USV bursts were shorter, with lower frequency dynamic, higher frequency characteristics, reduced frequency modulations, reduced frequency jumps and higher power characteristics compared to USVs emitted in most other contexts (**Figure 3**; see also results in the main text).

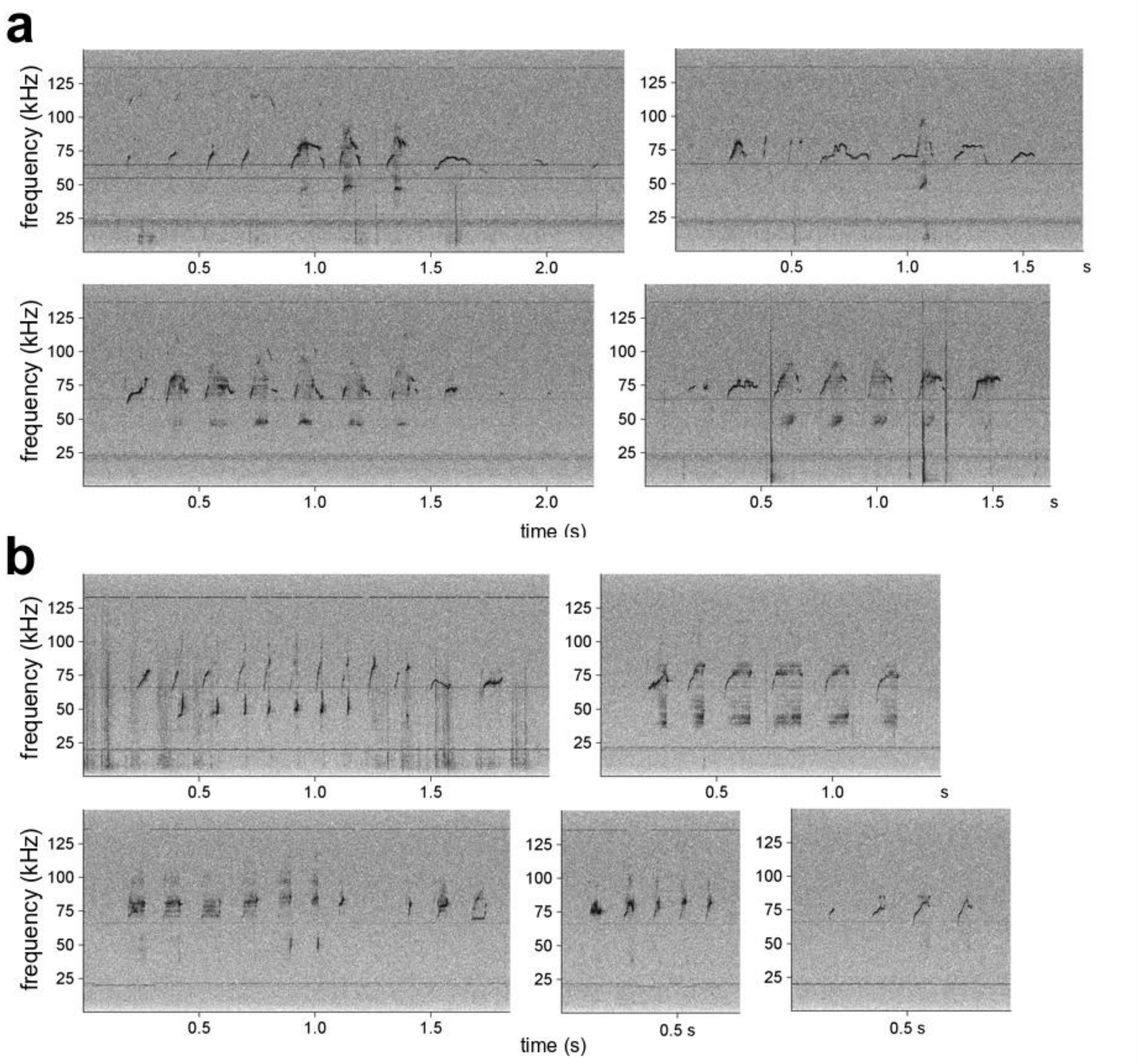


**Supplementary Figure Urination-1**: Spectrograms of USV bursts emitted during urination in WT mice. USV bursts emitted during urination by (a) a WT male aged three months, and (b) a WT female aged three months (FFT-length: 1024 points, 16-bit format, 300 kHz sampling frequency, 75% overlap).

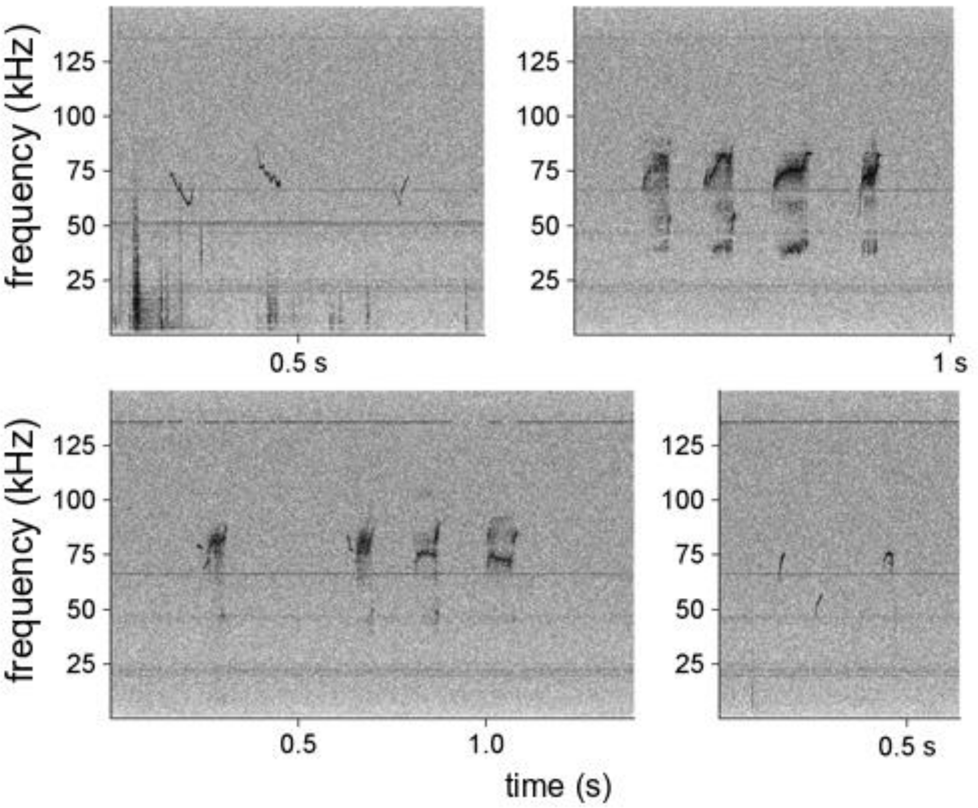


**Supplementary Figure Urination-2**: Spectrograms of USV bursts emitted during urination in *Shank3*$^{-/-}$ female mice aged three months (FFT-length: 1024 points, 16-bit format, 300 kHz sampling frequency, 75% overlap).

In our setting, usually, these "vocalised urination" sequences were recorded when the acting mouse was localised in one of the two corners the furthest from the nest (**Supplementary Figure Urination-3**). This was not always the case in *Shank3*$^{-/-}$ mice (**Supplementary Figure Urination-4**). In some pairs, only one mouse was displaying this behaviour, while in other pairs both mice displayed this behaviour (**Supplementary Tables Urination-I and II**). Interestingly, if this behaviour was unbalanced between the two mice of the pair in the youngest age class, it tended to remain unbalanced in older age classes (**Supplementary Tables Urination-I and II**). According to these observations, we concluded that not all urine deposits were accompanied by USV emission. Up to now, we cannot determine the factors ruling the expression of this behaviour. To our knowledge, there was only one mention of USVs emitted during urination, but this was not investigated further (Hoier et al., 2016). The expression of this behaviour might be related to hierarchy establishment, but this would need further investigations to be cleared.

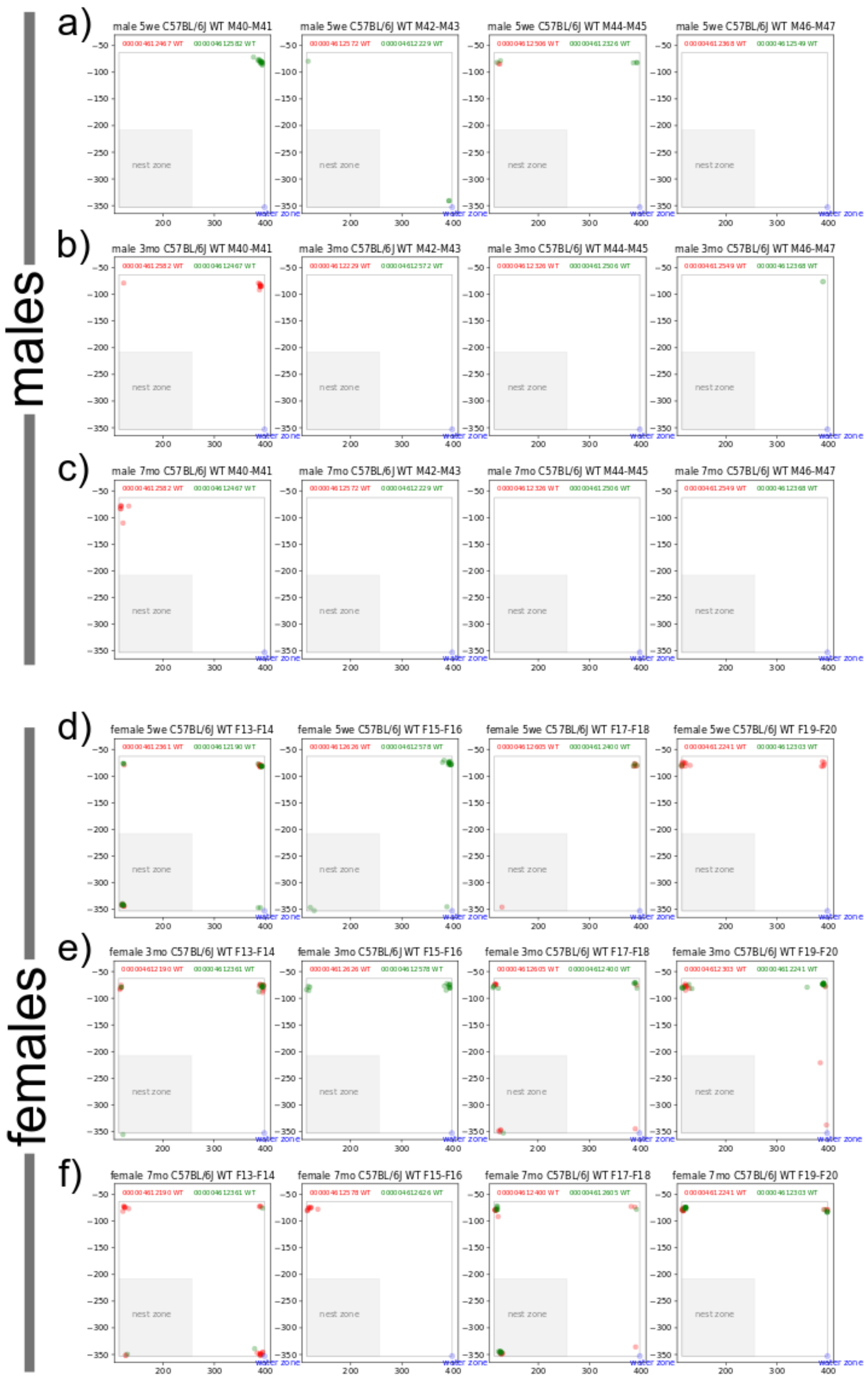


**Supplementary Figure Urination-3**: Urine deposits accompanied by USV bursts in WT mice. Location of the mouse emitting a USV burst during urination in males aged of a) 5 weeks, b) 3 months, and c) 7 months as well as in females aged of d) 5 weeks, e) 3 months and f) 7 months. The nest zone (i.e., the zone around the house) as well as the water zone (i.e., the access to the water bottle) are depicted within the cage. Colours (red and green) depict the individuals.

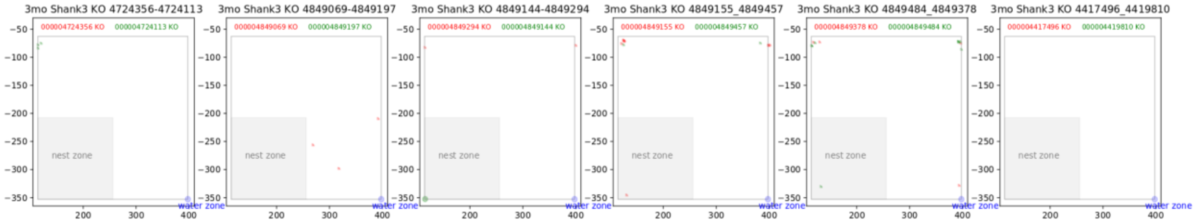


**Supplementary Figure Urination-4**: Urine deposits accompanied by USV bursts in *Shank3*$^{-/-}$ mice. Location of the mouse emitting a USV burst during urination in *Shank3*$^{-/-}$ females aged of 3 months. The nest zone (i.e., the zone around the house) as well as the water zone (i.e., the access to the water bottle) are depicted within the cage. Colours (red and green) depict the individuals.
