## Supplementary material for "Spontaneous social communication in laboratory mice - placing ultrasonic vocalizations in their behavioral context"

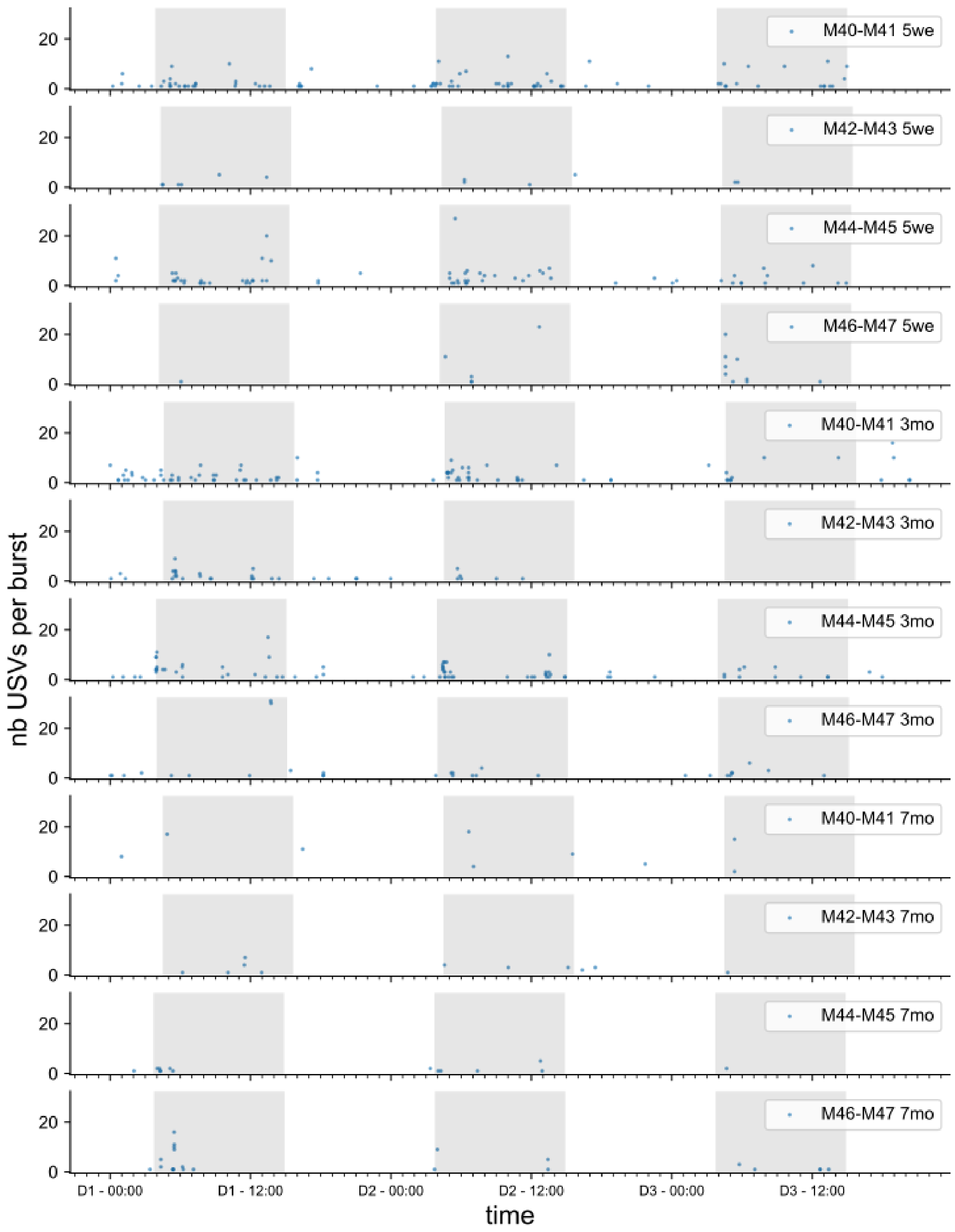


**Supplementary Figure S1.** Timeline of USV burst emission in WT male pairs. We represented the amount of USVs per burst as a function of the time of emission over the three days of recording in male pairs, recorded at 5 weeks, 3 months and 7 months.

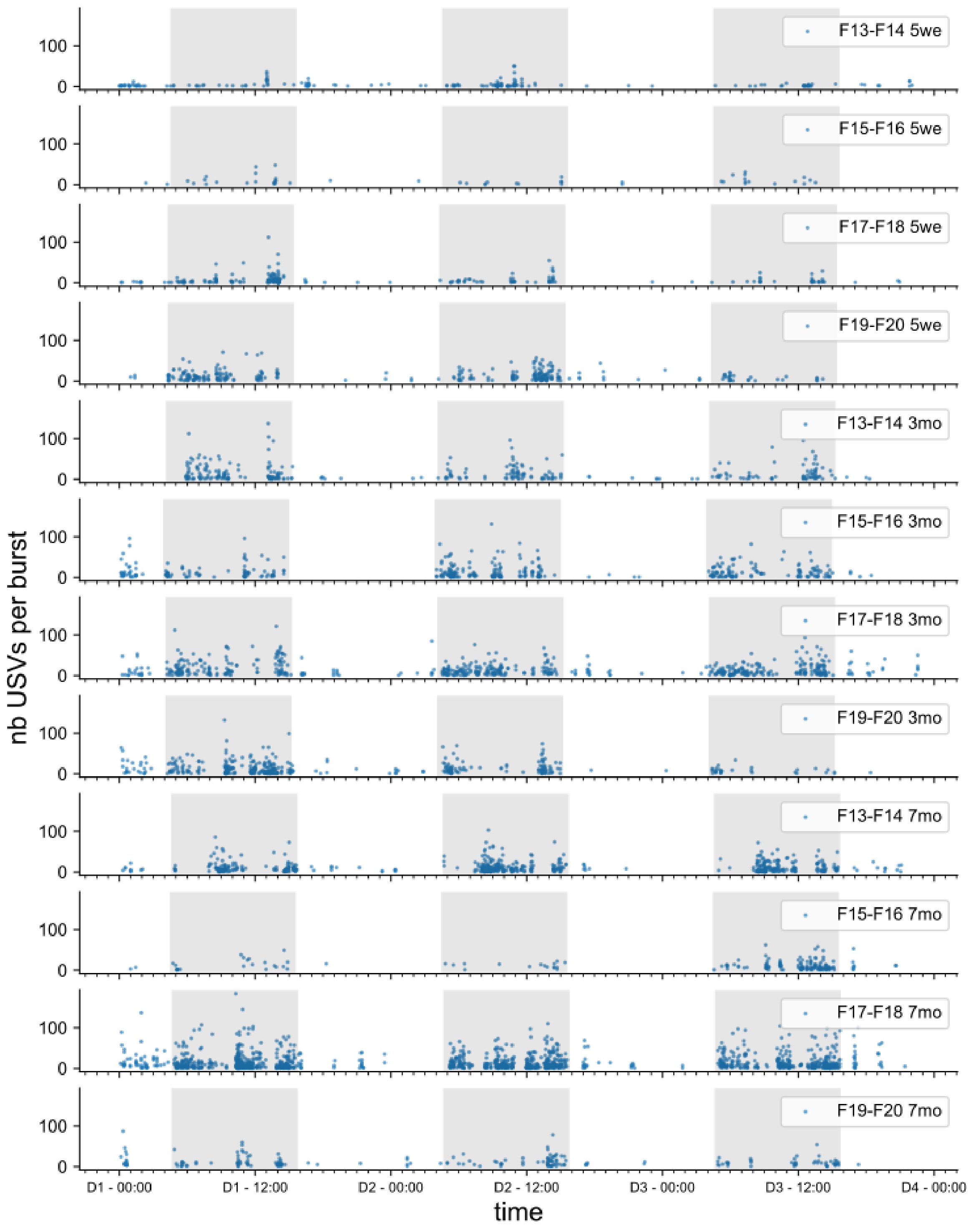


**Supplementary Figure S2.** Timeline of USV burst emission in WT female pairs. We represented the amount of USVs per burst as a function of the time of emission over the three days of recording in WT female pairs, recorded at 5 weeks, 3 months and 7 months.

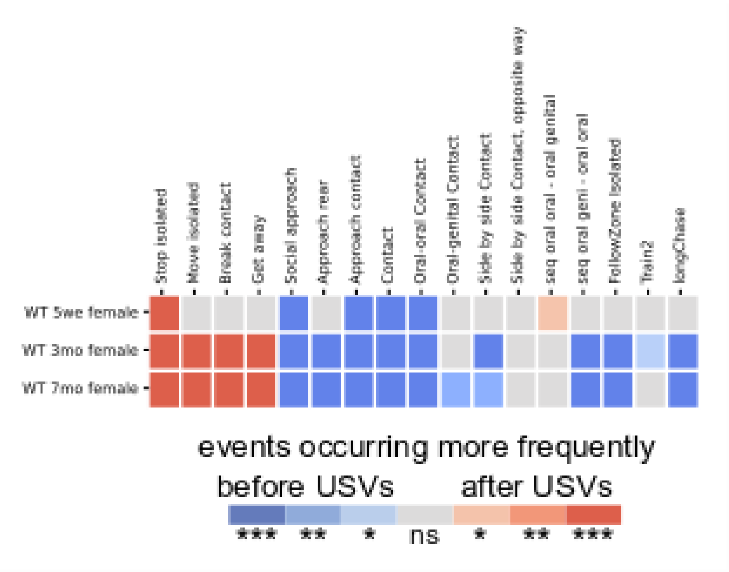


**Supplementary Figure S3.** Temporal organization of behavioral events around USVs in a time window of one second before and one second after USV. We calculated the proportion of the time window covered by the behavioral event before and after USV, for each USV. These proportions were compared using the central limit theorem. Colors of the heatmap depict the significance of the difference as well as the direction (blue colors: the behavior is significantly more expressed before the USV; red colors: the behavior is significantly more expressed after the USV).

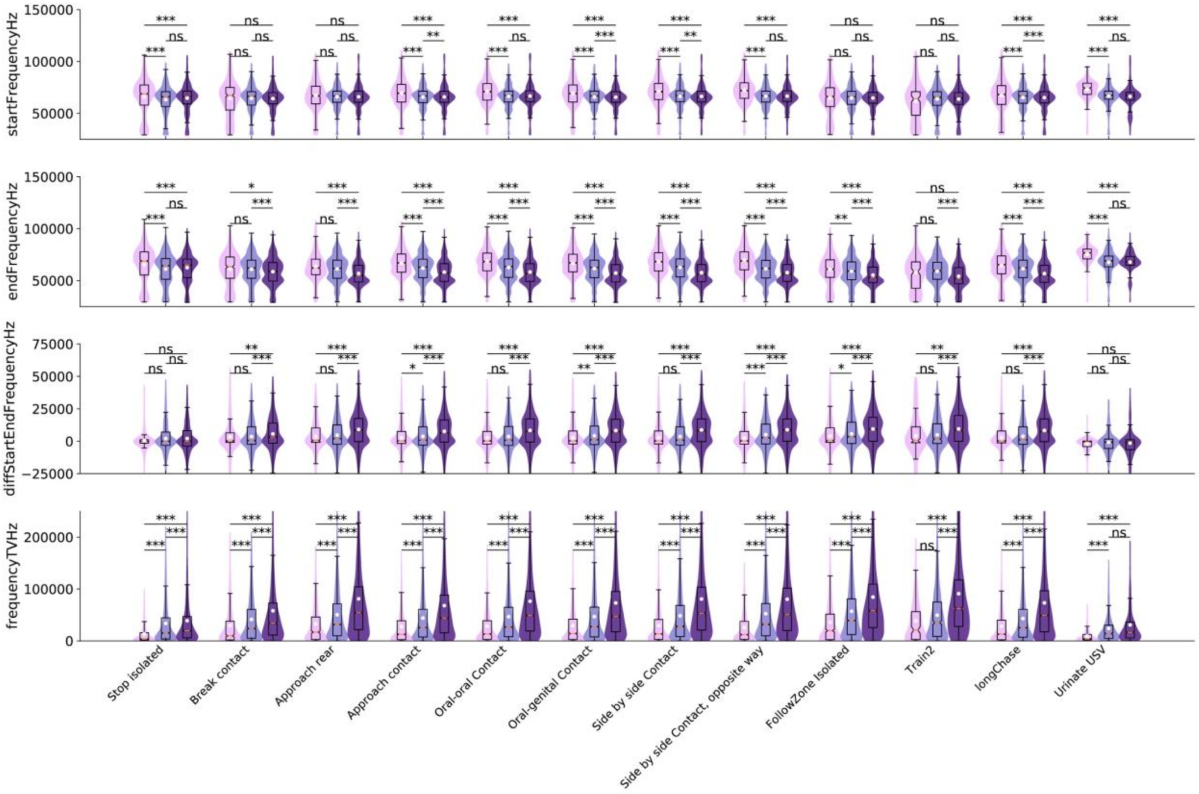
startFrequencyHz
150000
100000
50000
endFrequencyHz
150000
100000
50000
diffStartEndFrequencyHz
75000
50000
25000
0
−25000
frequencyTVHz
200000
100000
0
Stop isolated
Break contact
Approach rear
Approach contact
Oral-oral Contact
Oral-genital Contact
Side by side Contact
Side by side Contact, opposite way
FollowZone Isolated
Train2
longChase
Urinate USV

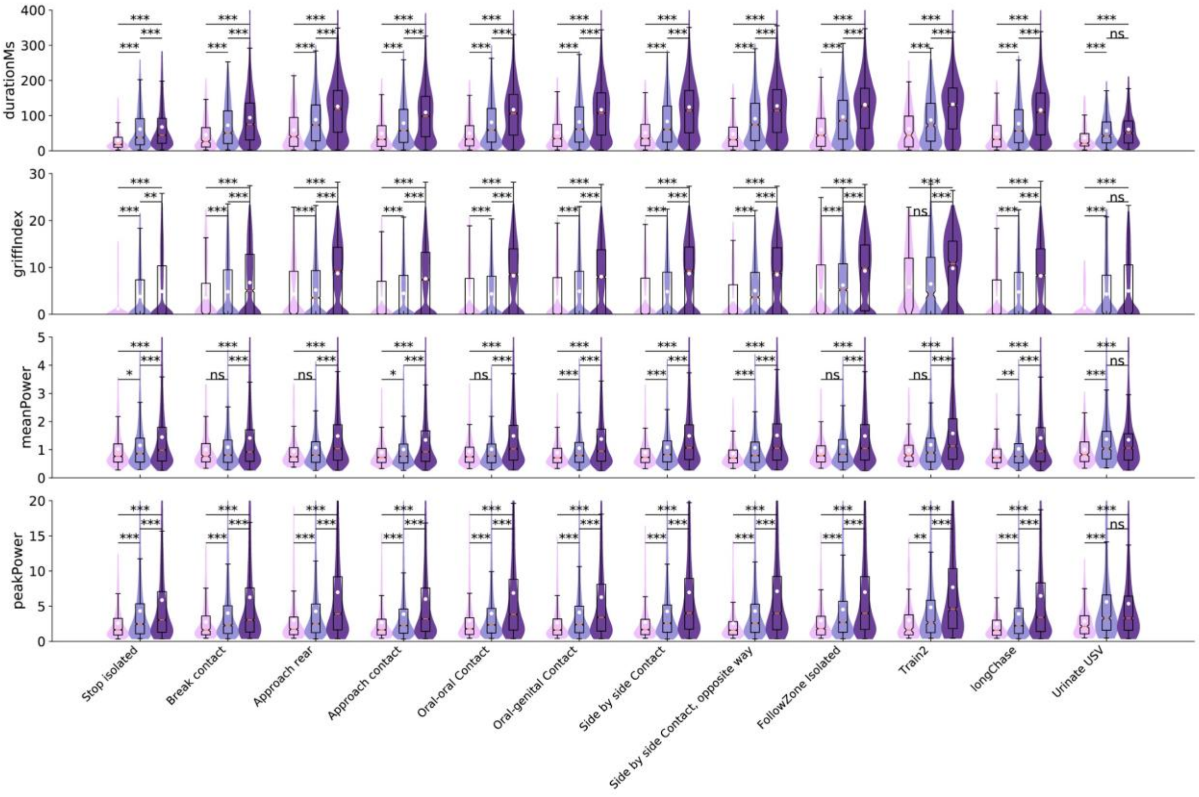
durationMs
400
300
200
100
0
griffIndex
30
20
10
0
meanPower
5
4
3
2
1
0
peakPower
20
15
10
5
0
Stop isolated
Break contact
Approach rear
Approach contact
Oral-oral Contact
Oral-genital Contact
Side by side Contact
Side by side Contact, opposite way
FollowZone Isolated
Train2
longChase
Urinate USV

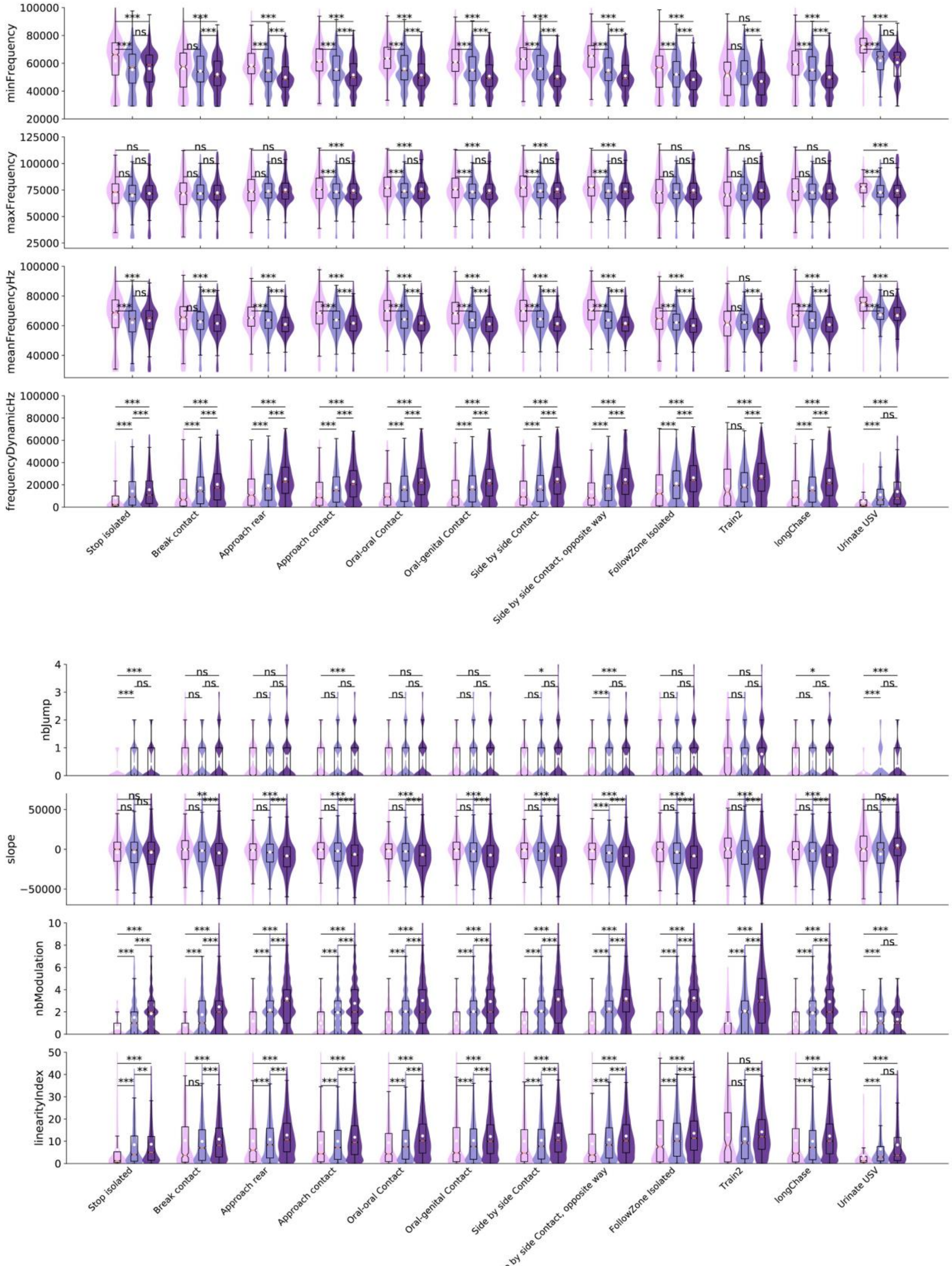


**Supplementary Figure S4.** Variations of the acoustic characteristics in the three age classes for all acoustic variables tested. Age-related variations are tested using Mann-Whitney U-tests and a Bonferroni correction for multiple testing is applied.

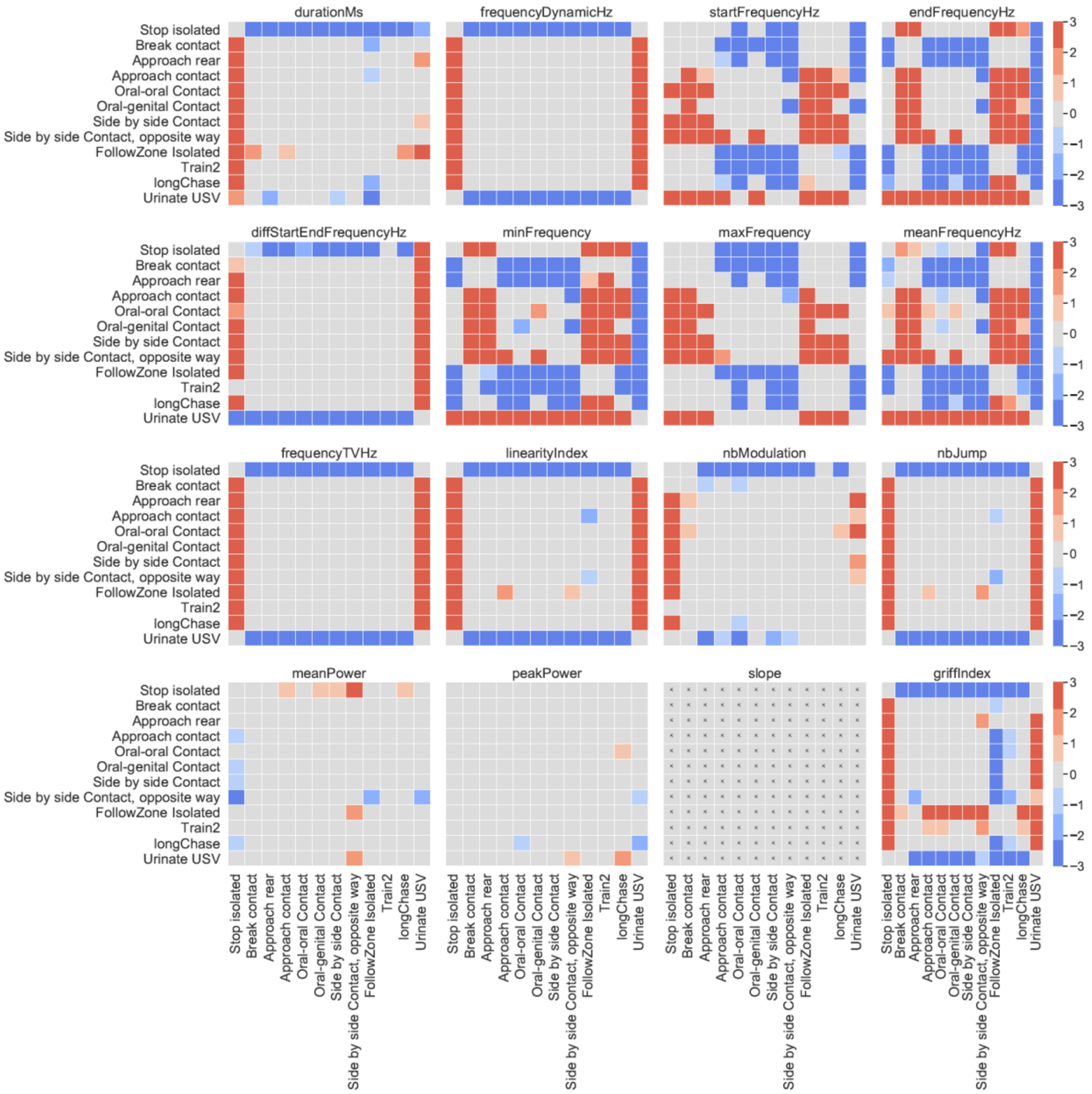


**Supplementary Figure S5.** Variations of the acoustic features of USVs emitted in different behavioral contexts in 5-week-old pairs of female WT mice. All acoustic traits measured for each USV are depicted. Color intensity represented the significance levels of the tests, while blue and red colors represented the direction of change: red: the acoustic trait of USVs emitted in the y-label event is higher than the same acoustic traits in USVs emitted in the x-label event; blue: the acoustic trait of USVs emitted in the y-label event is lower than the same acoustic traits in USVs emitted in the x-label event. For instance, USVs emitted in stop isolated were significantly shorter than USVs emitted in all other contexts, except urinate sequences that was accompanied by significantly longer USVs.

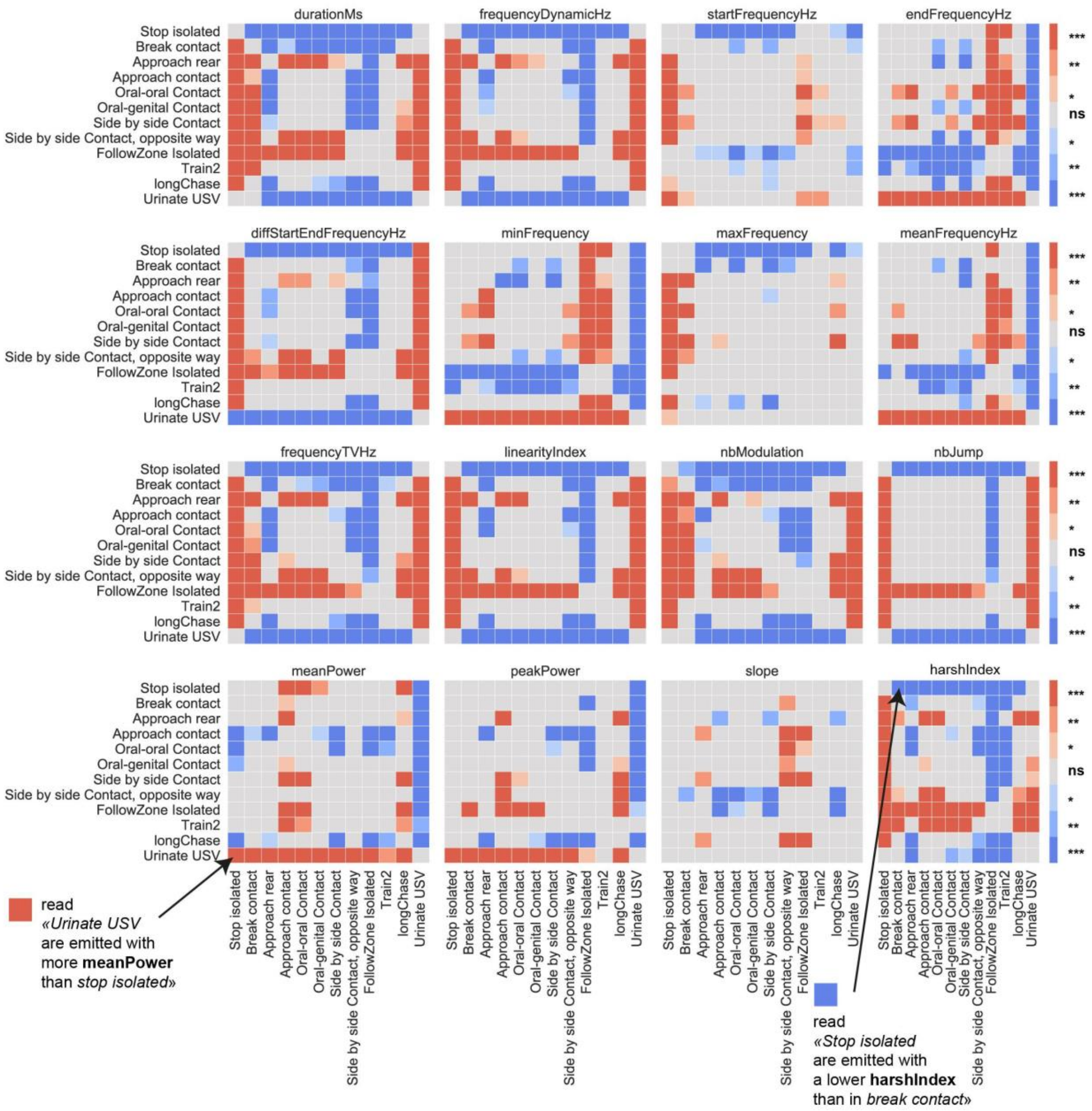


**Supplementary Figure S6.** Variations of the acoustic features of USVs emitted in different behavioral contexts in 3-month-old pairs of female WT mice. All acoustic traits measured for each USV are depicted. Color intensity represented the significance levels of the tests, while blue and red colors represented the direction of change: red: the acoustic trait of USVs emitted in the y-label event is higher than the same acoustic traits in USVs emitted in the x-label event; blue: the acoustic trait of USVs emitted in the y-label event is lower than the same acoustic traits in USVs emitted in the x-label event. For instance, USVs emitted in stop isolated were significantly shorter than USVs emitted in all other contexts, except urinate sequences that was accompanied by significantly longer USVs.

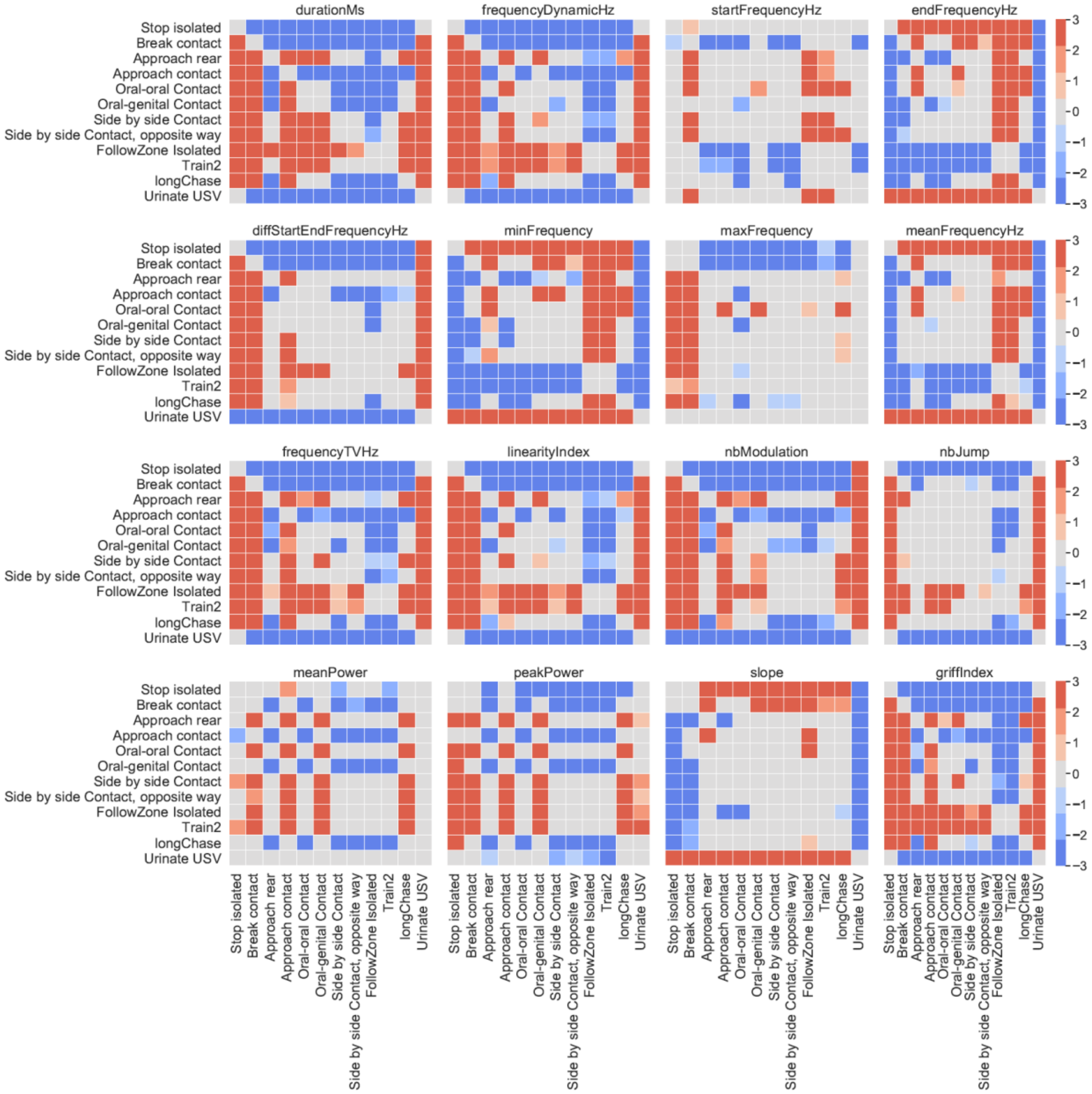


**Supplementary Figure S7.** Variations of the acoustic features of USVs emitted in different behavioral contexts in 7-month-old pairs of female WT mice. All acoustic traits measured for each USV are depicted. Color intensity represented the significance levels of the tests, while blue and red colors represented the direction of change: red: the acoustic trait of USVs emitted in the y-label event is higher than the same acoustic traits in USVs emitted in the x-label event; blue: the acoustic trait of USVs emitted in the y-label event is lower than the same acoustic traits in USVs emitted in the x-label event. For instance, USVs emitted in stop isolated were significantly shorter than USVs emitted in all other contexts, except urinate sequences that was accompanied by significantly longer USVs.

**Supplementary Table I.** Amount of USVs recorded over three days in WT and Shank3 same-sex pairs of mice. nbUsv: total number of USVs recorded. nbBurst: total number of USV burst recorded. mean nbUsv/Burst: mean number of USVs per burst. std nbUsv/Burst: standard deviation of the number of USVs per burst.

| experiment | sex | age | strain | genotype | pair | nbUsv | nbBurst | mean nbUsv/Burst | std nbUsv/Burst |
|---|---|---|---|---|---|---|---|---|---|
| female 5we C57BL/6J WT F13-F1 | female | 5we | C57BL/6J | WT | F13-F14 | 881 | 214 | 4,22 | 5,56 |
| female 5we C57BL/6J WT F15-F1 | female | 5we | C57BL/6J | WT | F15-F16 | 501 | 57 | 10,86 | 9,30 |
| female 5we C57BL/6J WT F17-F1 | female | 5we | C57BL/6J | WT | F17-F18 | 1713 | 309 | 7,48 | 9,78 |
| female 5we C57BL/6J WT F19-F2 | female | 5we | C57BL/6J | WT | F19-F20 | 4072 | 341 | 14,17 | 9,94 |
| female 3mo C57BL/6J WT F13-F1 | female | 3mo | C57BL/6J | WT | F13-F14 | 4719 | 341 | 16,11 | 17,46 |
| female 3mo C57BL/6J WT F15-F1 | female | 3mo | C57BL/6J | WT | F15-F16 | 5938 | 479 | 14,93 | 13,88 |
| female 3mo C57BL/6J WT F17-F1 | female | 3mo | C57BL/6J | WT | F17-F18 | 9921 | 881 | 12,14 | 10,32 |
| female 3mo C57BL/6J WT F19-F2 | female | 3mo | C57BL/6J | WT | F19-F20 | 5779 | 490 | 14,21 | 13,31 |
| female 7mo C57BL/6J WT F13-F1 | female | 7mo | C57BL/6J | WT | F13-F14 | 7196 | 792 | 9,77 | 9,67 |
| female 7mo C57BL/6J WT F15-F1 | female | 7mo | C57BL/6J | WT | F15-F16 | 2161 | 266 | 9,23 | 6,97 |
| female 7mo C57BL/6J WT F17-F1 | female | 7mo | C57BL/6J | WT | F17-F18 | 22027 | 1637 | 16,40 | 16,41 |
| female 7mo C57BL/6J WT F19-F2 | female | 7mo | C57BL/6J | WT | F19-F20 | 2570 | 266 | 10,30 | 8,55 |
| male 5we C57BL/6J WT M40-M41 | male | 5we | C57BL/6J | WT | M40-M41 | 251 | 114 | 2,49 | 2,86 |
| male 5we C57BL/6J WT M42-M43 | male | 5we | C57BL/6J | WT | M42-M43 | 29 | 15 | 2,73 | 1,98 |
| male 5we C57BL/6J WT M44-M45 | male | 5we | C57BL/6J | WT | M44-M45 | 252 | 79 | 3,25 | 3,55 |
| male 5we C57BL/6J WT M46-M47 | male | 5we | C57BL/6J | WT | M46-M47 | 98 | 15 | 7,00 | 7,02 |
| male 3mo C57BL/6J WT M40-M41 | male | 3mo | C57BL/6J | WT | M40-M41 | 260 | 103 | 2,66 | 2,70 |
| male 3mo C57BL/6J WT M42-M43 | male | 3mo | C57BL/6J | WT | M42-M43 | 70 | 48 | 1,54 | 1,83 |
| male 3mo C57BL/6J WT M44-M45 | male | 3mo | C57BL/6J | WT | M44-M45 | 250 | 103 | 2,51 | 2,82 |
| male 3mo C57BL/6J WT M46-M47 | male | 3mo | C57BL/6J | WT | M46-M47 | 108 | 38 | 3,00 | 5,48 |
| male 7mo C57BL/6J WT M40-M41 | male | 7mo | C57BL/6J | WT | M40-M41 | 99 | 10 | 11,80 | 5,06 |
| male 7mo C57BL/6J WT M42-M43 | male | 7mo | C57BL/6J | WT | M42-M43 | 32 | 18 | 1,89 | 1,79 |
| male 7mo C57BL/6J WT M44-M45 | male | 7mo | C57BL/6J | WT | M44-M45 | 25 | 16 | 3,19 | 2,67 |
| male 7mo C57BL/6J WT M46-M47 | male | 7mo | C57BL/6J | WT | M46-M47 | 85 | 31 | 3,26 | 4,12 |
| female 3mo Shank3 KO 4724356- | female | 3mo | Shank3 | KO | 4724356-4724113 | 3315 | 452 | 7,67 | 6,61 |
| female 3mo Shank3 KO 4849069- | female | 3mo | Shank3 | KO | 4849069-4849197 | 2216 | 334 | 7,07 | 5,21 |
| female 3mo Shank3 KO 4849144- | female | 3mo | Shank3 | KO | 4849144-4849294 | 8666 | 844 | 10,74 | 8,51 |
| female 3mo Shank3 KO 4849155_ | female | 3mo | Shank3 | KO | 4849155_4849457 | 2972 | 386 | 8,45 | 7,04 |
| female 3mo Shank3 KO 4849484_ | female | 3mo | Shank3 | KO | 4849484_4849378 | 11032 | 1150 | 9,95 | 10,45 |
| female 3mo Shank3 KO 4417496_ | female | 3mo | Shank3 | KO | 4417496_4419810 | 4961 | 669 | 7,97 | 5,97 |

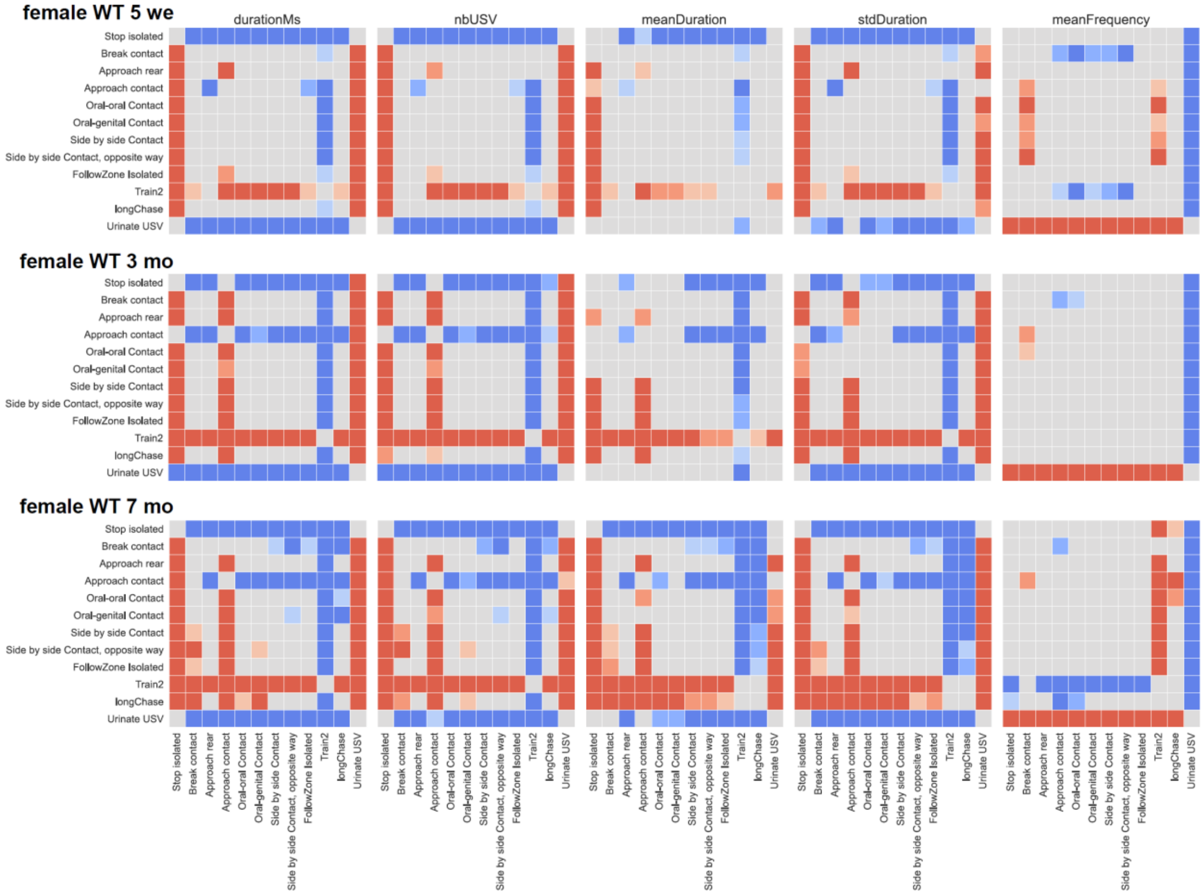


**Supplementary Figure S8.** Variations of the acoustic features of USV bursts emitted in different behavioral contexts in WT female mouse pairs. The total duration of the USV bursts, the number of USVs per USV burst, the mean duration of each USV within a burst, the standard deviation of the duration of the USVs within a burst and the mean frequency of the USVs within a burst were compared between the different contexts in WT female mice aged of 5 weeks (upper panel), 3 months (middle panel) and 7 months (lower panel). Color intensity represented the significance levels of the tests, while blue and red colors represented the direction of change: red: the acoustic trait of USVs emitted in the y-label event is higher than the same acoustic traits in USVs emitted in the x-label event; blue: the acoustic trait of USVs emitted in the y-label event is lower than the same acoustic traits in USVs emitted in the x-label event. For instance, USV bursts emitted at 5 weeks of age during stop isolated were significantly shorter than USV bursts emitted in any other contexts except urination.

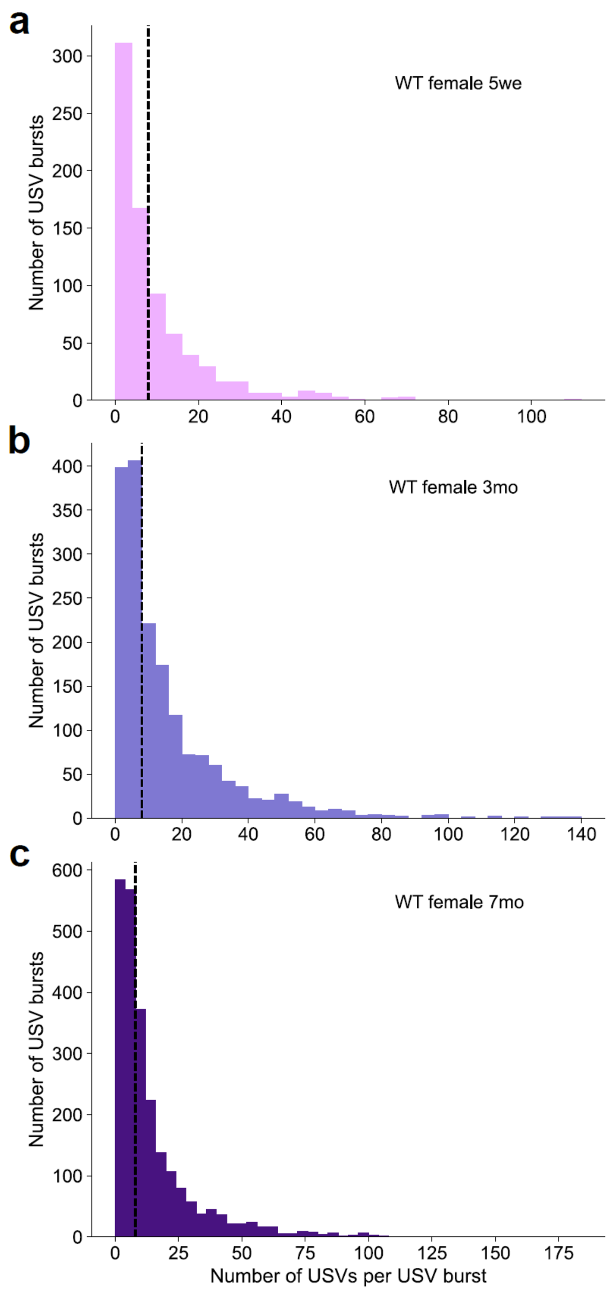


**Supplementary Figure S9.** Distribution of the USV bursts according to the number of USVs encompassed in pairs of WT female mice aged of 5 weeks, 3 months and 7 months. The threshold was set at 8 USVs per burst.

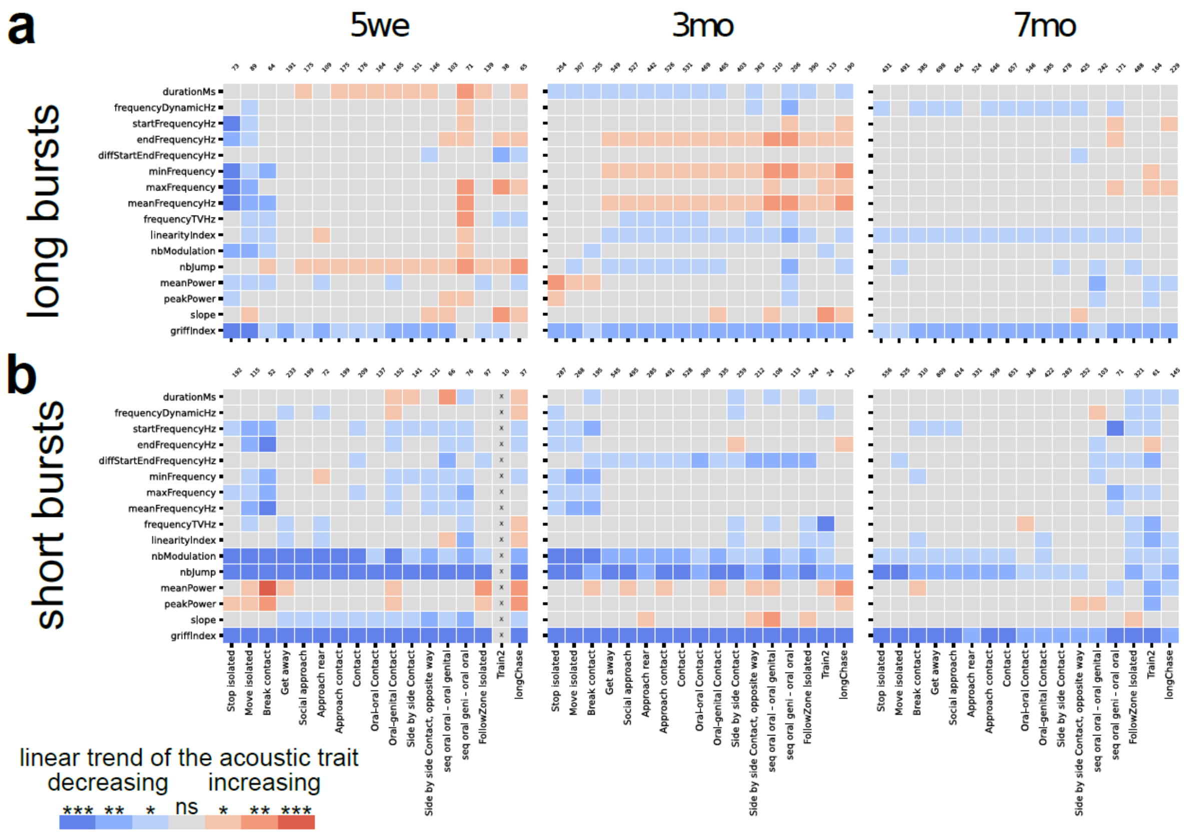


**Supplementary Figure S10.** Linear trend of the intonation within (a) long and (b) short bursts.

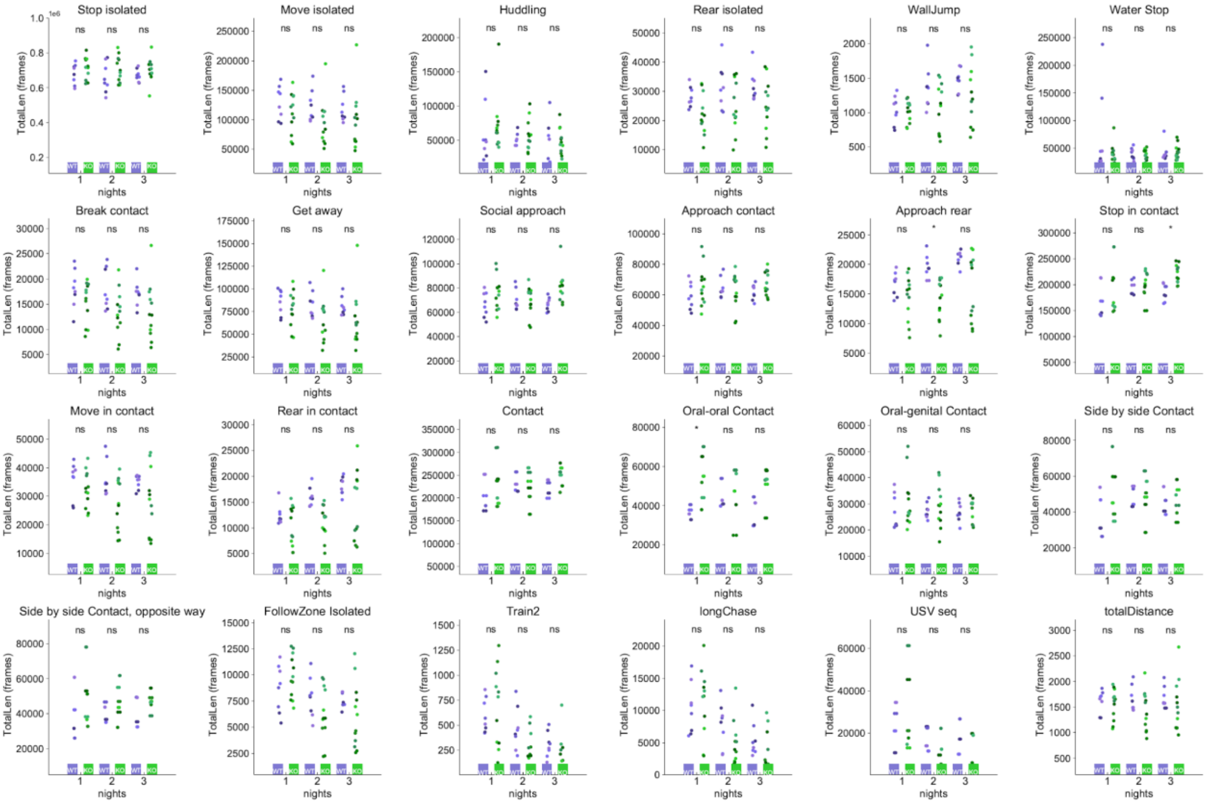


**Supplementary Figure S11.** Behavioral profiles of *Shank3*$^{-/-}$ female mice and WT female mice aged of 3 months over the three nights of recording. We depicted the total time spent in each behavior for each individual. We matched the colors of the points according to the pairs. We used Mann-Whitney U-tests to compare WT versus *Shank3*$^{-/-}$ (KO) mice within each night, and conducted a Bonferroni correction for multiple testing (for each night, 24 tests were conducted).

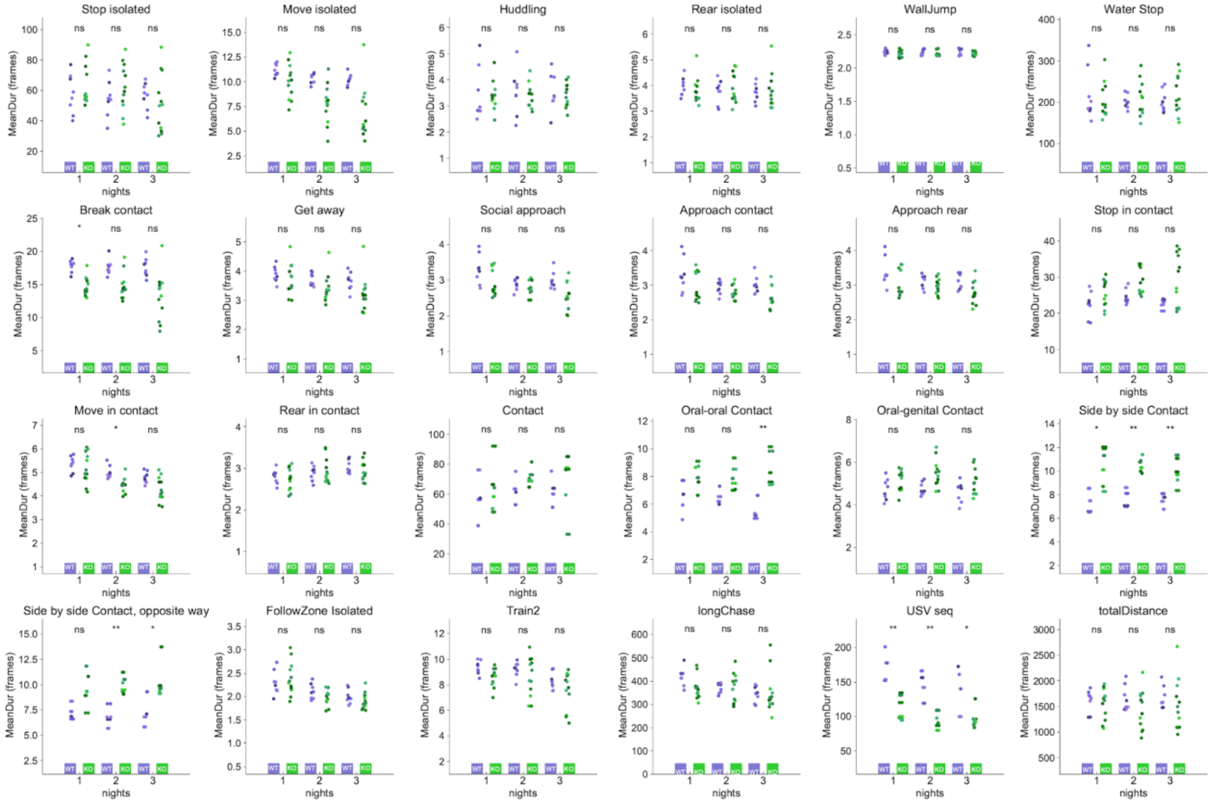


**Supplementary Figure S12.** Behavioral profiles of *Shank3*$^{-/-}$ female mice and WT female mice aged of 3 months over the three nights of recording. We depicted the mean duration of each behavior for each individual. We matched the colors of the points according to the pairs. We used Mann-Whitney U-tests to compare WT versus *Shank3*$^{-/-}$ (KO) mice within each night, and conducted a Bonferroni correction for multiple testing (for each night, 24 tests were conducted).

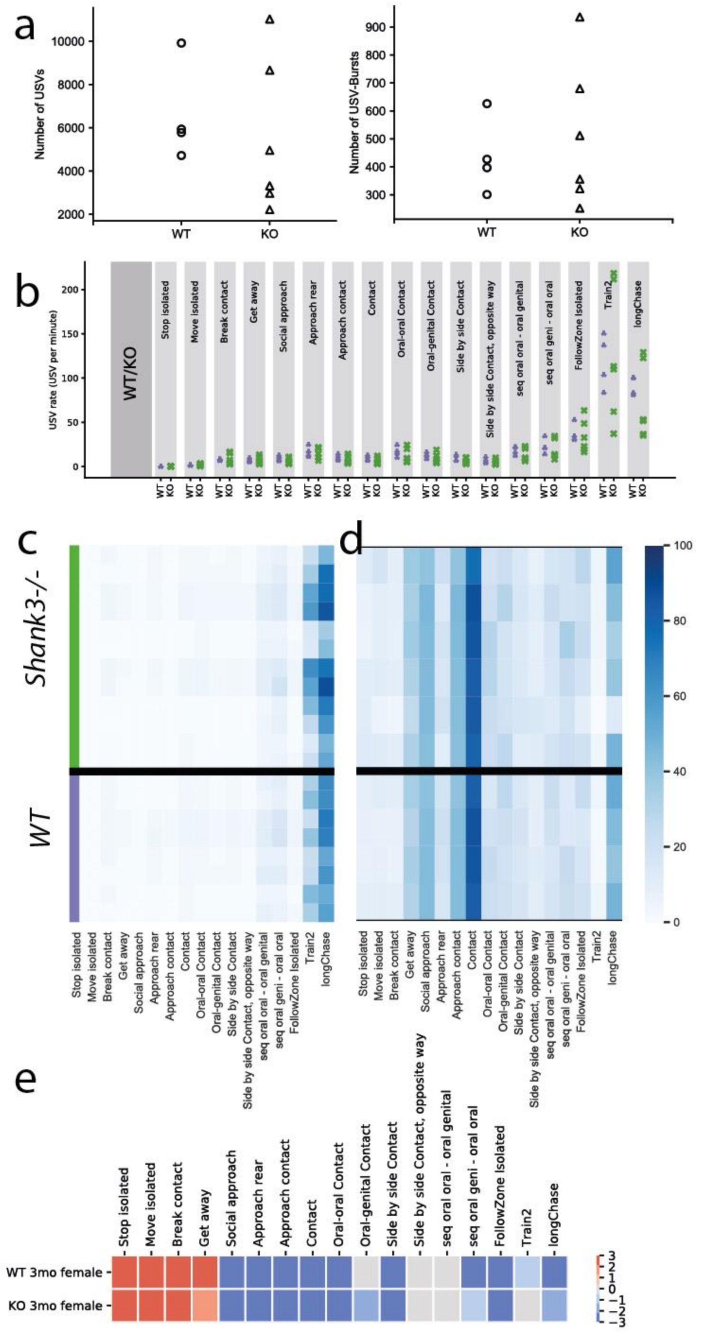


**Supplementary Figure S13.** Contexts of emission of USVs in *Shank3*$^{-/-}$ female mice. (a) Number of USVs (left panel) and number of USV bursts (right panel) emitted by *Shank3*$^{-/-}$ mice and WT mice. (b) Rate of emission of USVs in the different behaviors in WT females and *Shank3*$^{-/-}$ females aged of 3 months. (c) Proportion of the frames of the USVs overlapping with the different behavioral events across age and sex classes. The reference is the USVs. For dark blue, the figure reads like 80% of the USV duration were accompanied by event A. d) Proportion of the frames of the different behavioral events overlapping with USVs across age and sex classes. The reference is the behavioral events. For dark blue, the figure reads like 80% of the Train2 duration were accompanied by USVs.

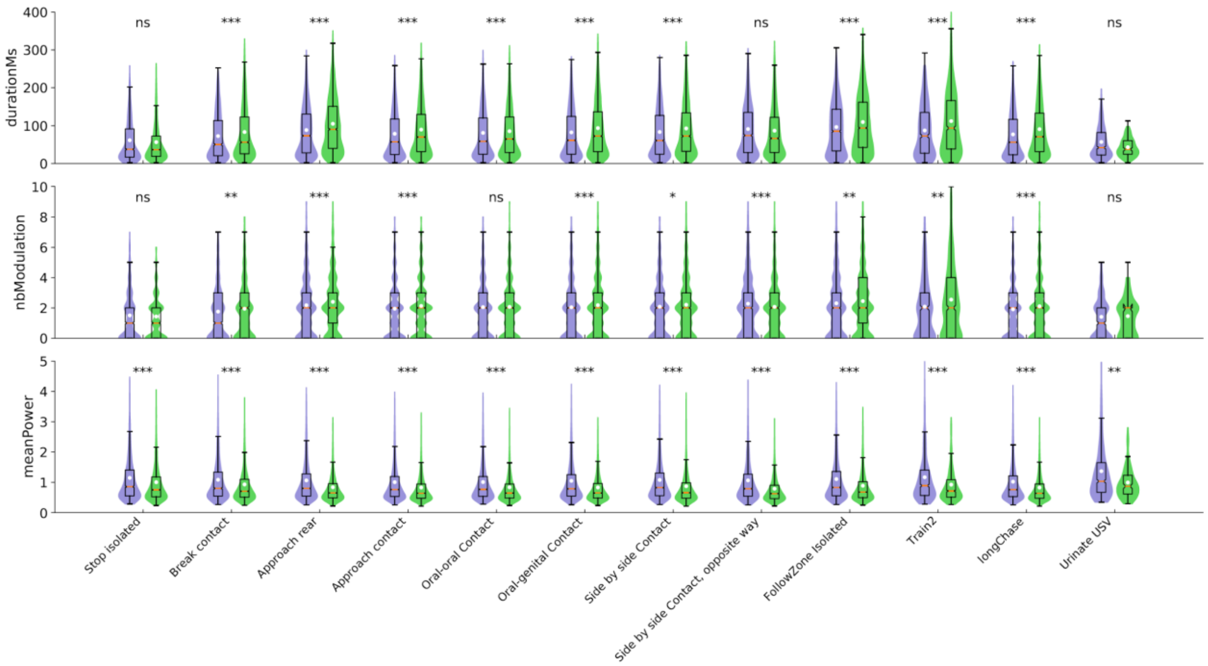


**Supplementary Figure S14.** Genotype-related differences in duration, number of modulations and mean power in USVs emitted in specific behavioral contexts.

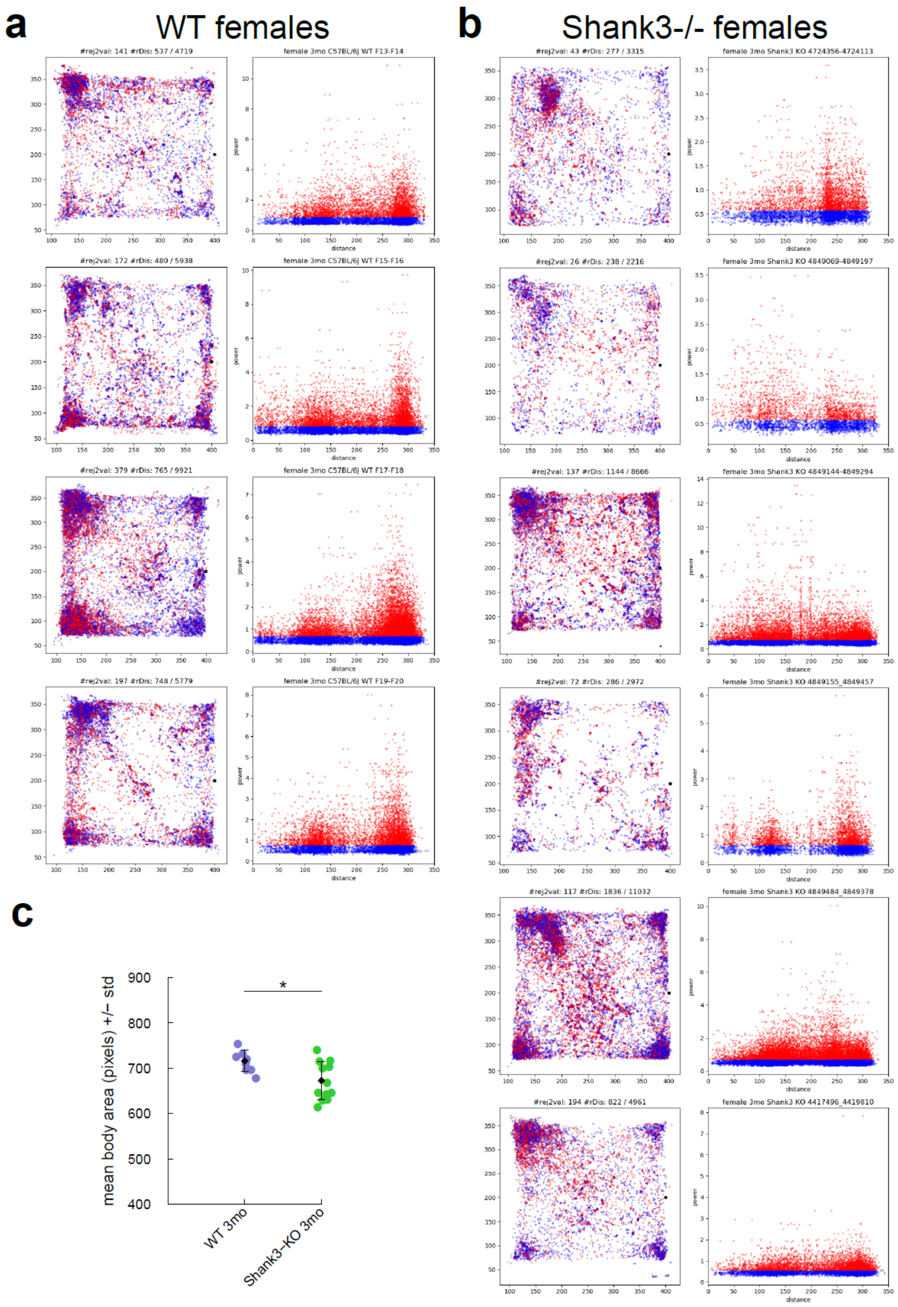


**Supplementary Figure S15.** Smaller body size might explain the lower power in USVs from *Shank3*$^{-/-}$ compared to WT more than the distance to the microphone during USV emission. Location of the pairs of mice and distance to the microphone during USV emission in (a) WT mice and (b) *Shank3*$^{-/-}$ mice. Left graphs: The mean position of the pair of mice over the whole USV duration is represented only when the two mice were detected within 10 cm from one another. Blue points represent USVs that were emitted with a power in the lowest 50$^{th}$ percentile of the power distribution for each experiment separately. Red points represent USVs that were emitted with a power in the highest 50$^{th}$ percentile of the power distribution for each experiment separately. The position of the microphone is depicted by a black dot in the middle of the right hand side of the cage. Right graphs: The power of the USVs recorded did not appeared to be correlated with the distance between the microphone and the position of the pair of the mice (the two mice were within 10 cm from one another). (c) Mean body surface measured on the masking detected by the Live Mouse Tracker system on the first xx hours of tracking in *Shank3*$^{-/-}$ female mice and in age-matched WT female mice. Mann-Whitney U-test was used for statistical comparison.

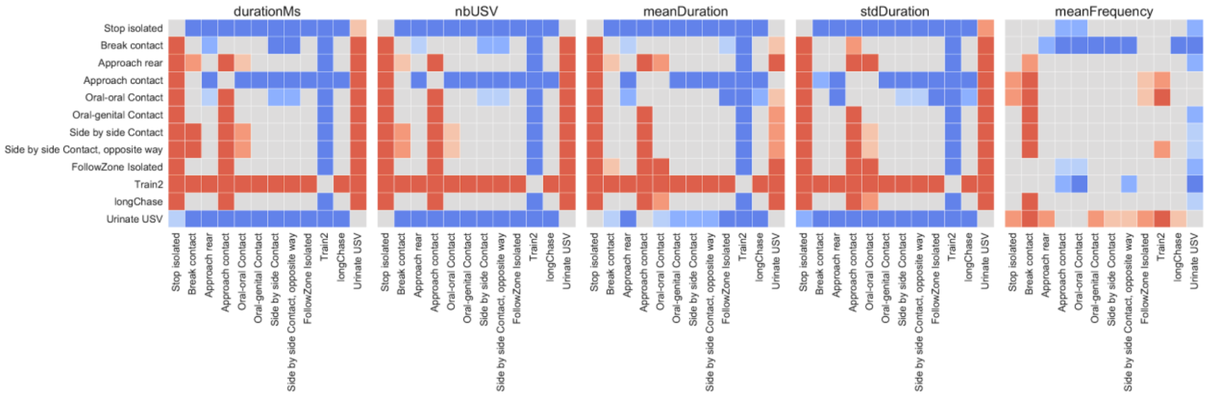


**Supplementary Figure S16.** Variations of the acoustic features of USV bursts emitted in different behavioral contexts in *Shank3*$^{-/-}$ female mouse pairs. The total duration of the USV bursts, the number of USVs per USV burst, the mean duration of each USV within a burst, the standard deviation of the duration of the USVs within a burst and the mean frequency of the USVs within a burst were compared between the different contexts in *Shank3*$^{-/-}$ female mice aged of 3 months. Color intensity represented the significance levels of the tests, while blue and red colors represented the direction of change: red: the acoustic trait of USVs emitted in the y-label event is higher than the same acoustic traits in USVs emitted in the x-label event; blue: the acoustic trait of USVs emitted in the y-label event is lower than the same acoustic traits in USVs emitted in the x-label event. For instance, USV bursts emitted at 5 weeks of age during stop isolated were significantly shorter than USV bursts emitted in any other contexts except urination.

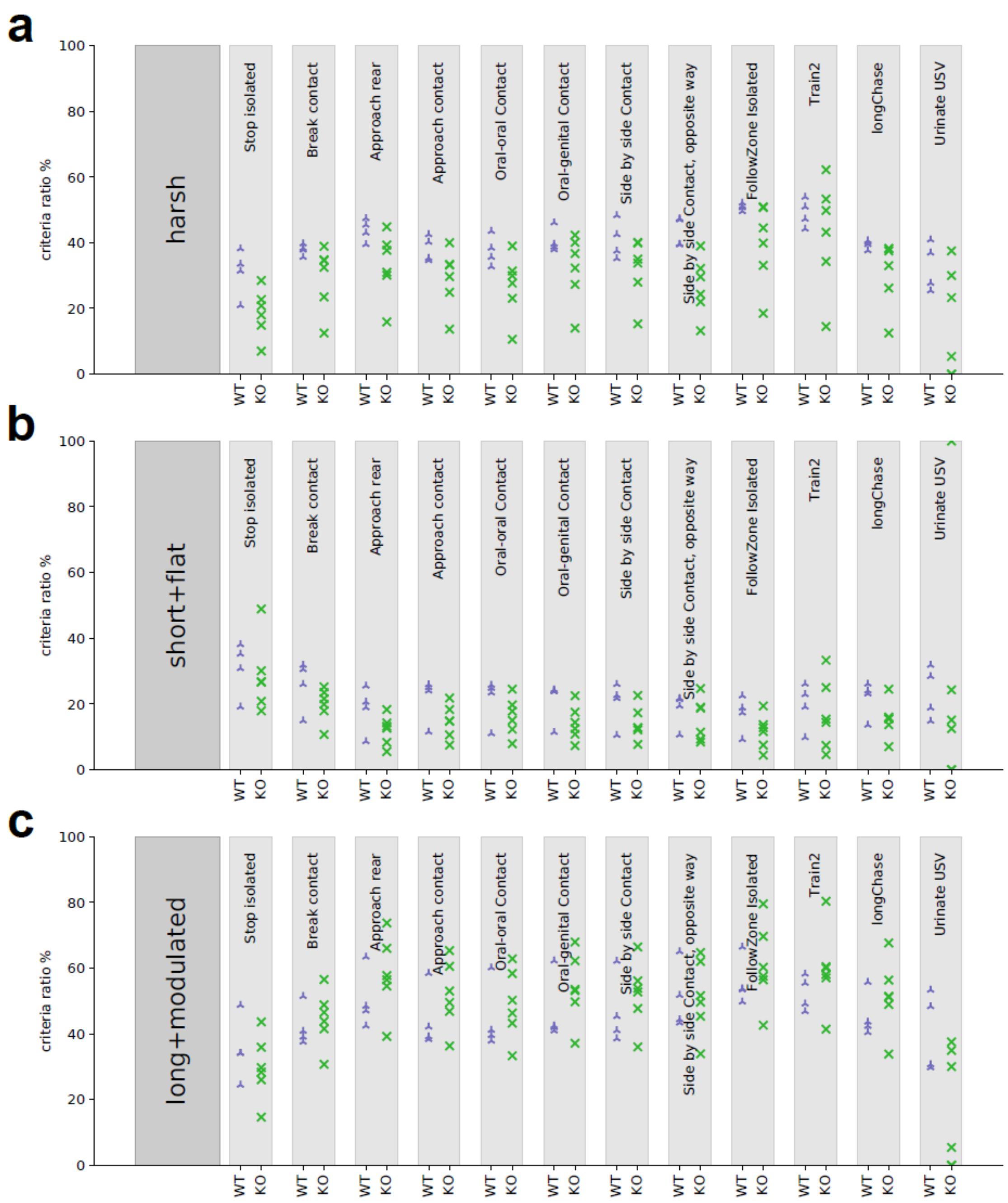


**Supplementary Figure S17.** Proportion of specific USV types across behavioral events in WT and *Shank3*$^{-/-}$ female mice aged of 3 months. (a) Proportion of harsh USVs in the different behavioral contexts observed. (b) Proportion of short and flat USVs in the different behavioral contexts observed. (c) Proportion of long modulated USVs in the different behavioral contexts observed.

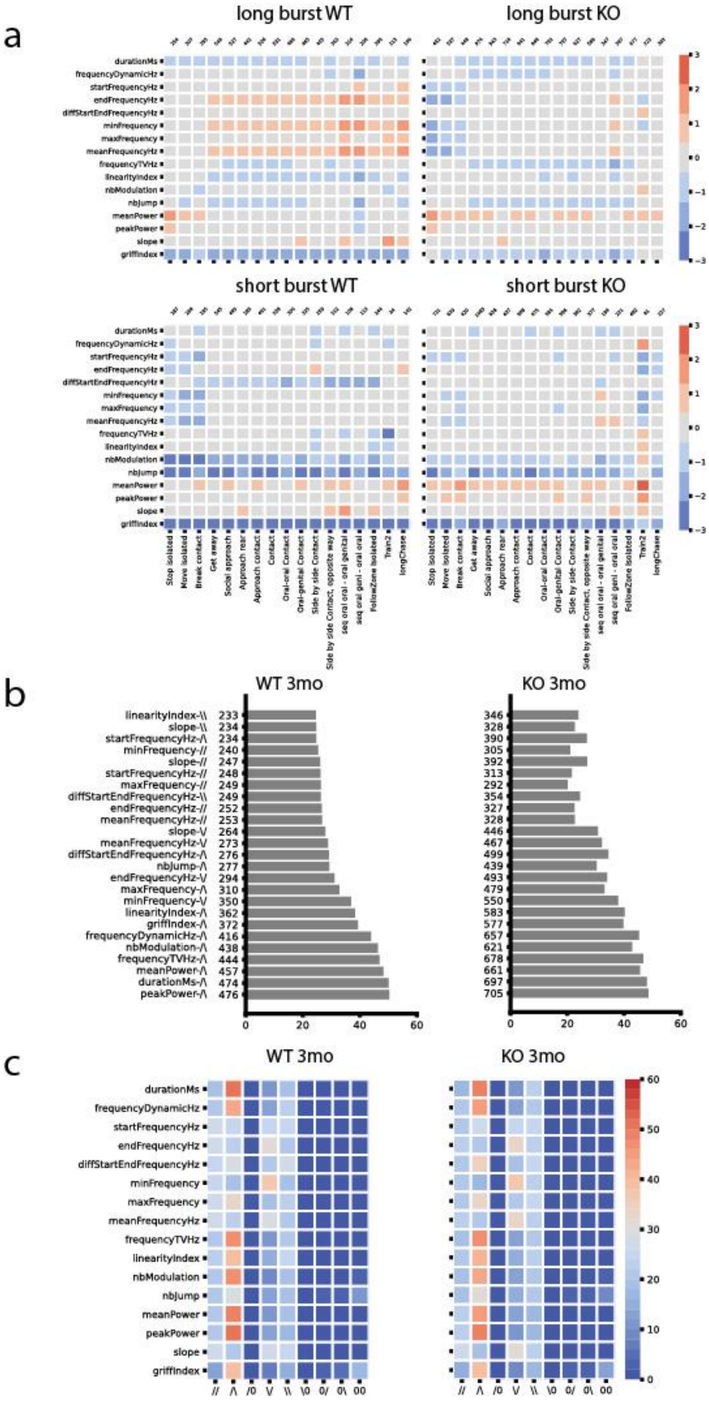

**Supplementary Figure S18.** (a) Direction of variations of the linear fit of each acoustic variable measured over the whole long (upper panels) and short (lower panels) USV bursts in WT mice (left) and *Shank3*$^{-/-}$ mice (right) aged of 3 months. (b) Proportion of occurrences of the 20 first most frequent USV burst types in WT (left) and in *Shank3*$^{-/-}$ (right) mice. (c) Proportion of occurrences of the different possibilities of intonations within long USV bursts in WT (left) and in *Shank3*$^{-/-}$ (right) mice aged of 3 months.

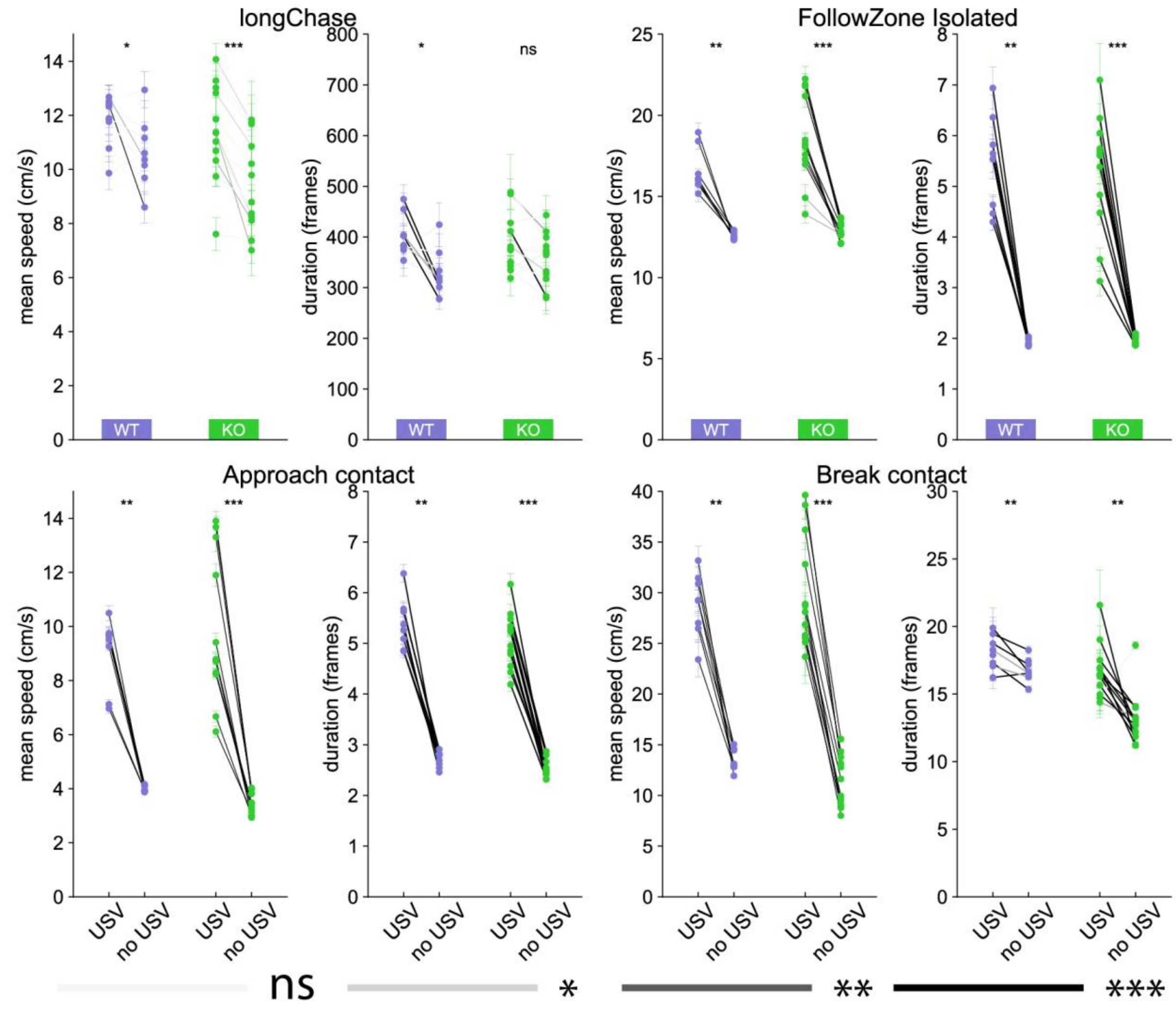


**Supplementary Figure S19.** Variations in mean speed and duration of each behavioral event according to the presence or absence of USVs. Variations of the mean speed of the animal performing the behavior (left panel) and of the event duration (right panel) for (a) longChase, (b) follow, (c) approach contact and (d) break contact. Significant differences using Mann-Whitney U-tests within each individual after Bonferroni correction (24 tests conducted for each behavioral event and variable tested) are depicted by the color of the segments linking the mean value with and without USVs: light grey: not significant, medium grey: adjusted p<0.05, dark grey: adjusted p<0.01, black: adjusted p<0.001.

**a**

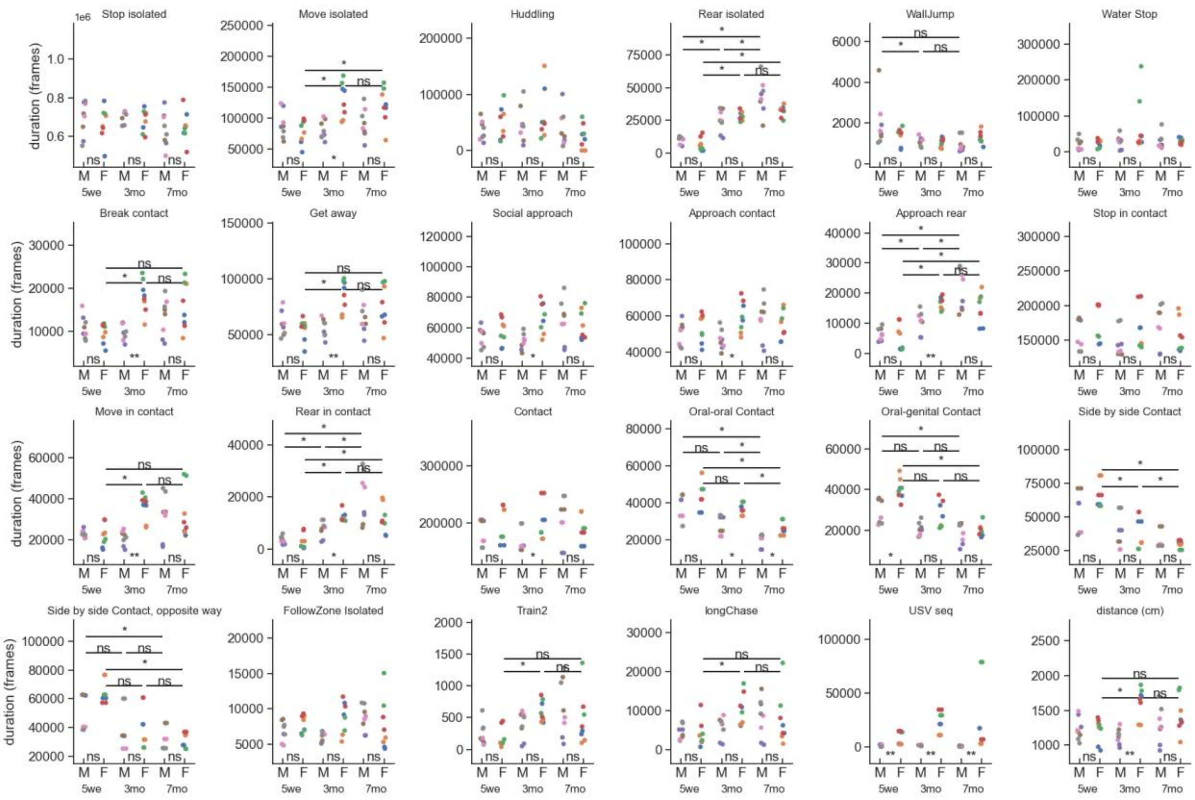
Stop isolated
Move isolated
Huddling
Rear isolated
WallJump
Water Stop
Break contact
Get away
Social approach
Approach contact
Approach rear
Stop in contact
Move in contact
Rear in contact
Contact
Oral-oral Contact
Oral-genital Contact
Side by side Contact
Side by side Contact, opposite way
FollowZone Isolated
Train2
longChase
USV seq
distance (cm)
duration (frames)
1e6
M F M F M F
5we 3mo 7mo

**b**

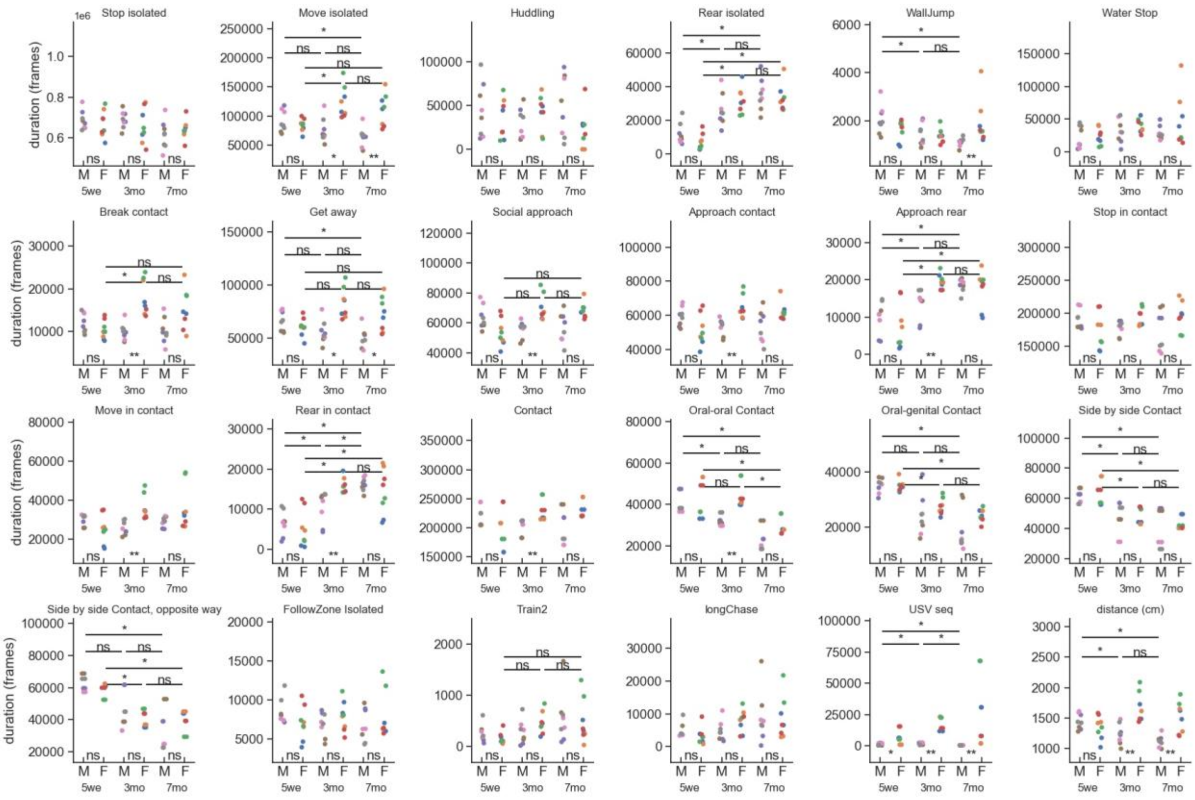
Stop isolated
Move isolated
Huddling
Rear isolated
WallJump
Water Stop
Break contact
Get away
Social approach
Approach contact
Approach rear
Stop in contact
Move in contact
Rear in contact
Contact
Oral-oral Contact
Oral-genital Contact
Side by side Contact
Side by side Contact, opposite way
FollowZone Isolated
Train2
longChase
USV seq
distance (cm)
duration (frames)
1e6
M F M F M F
5we 3mo 7mo

**c**

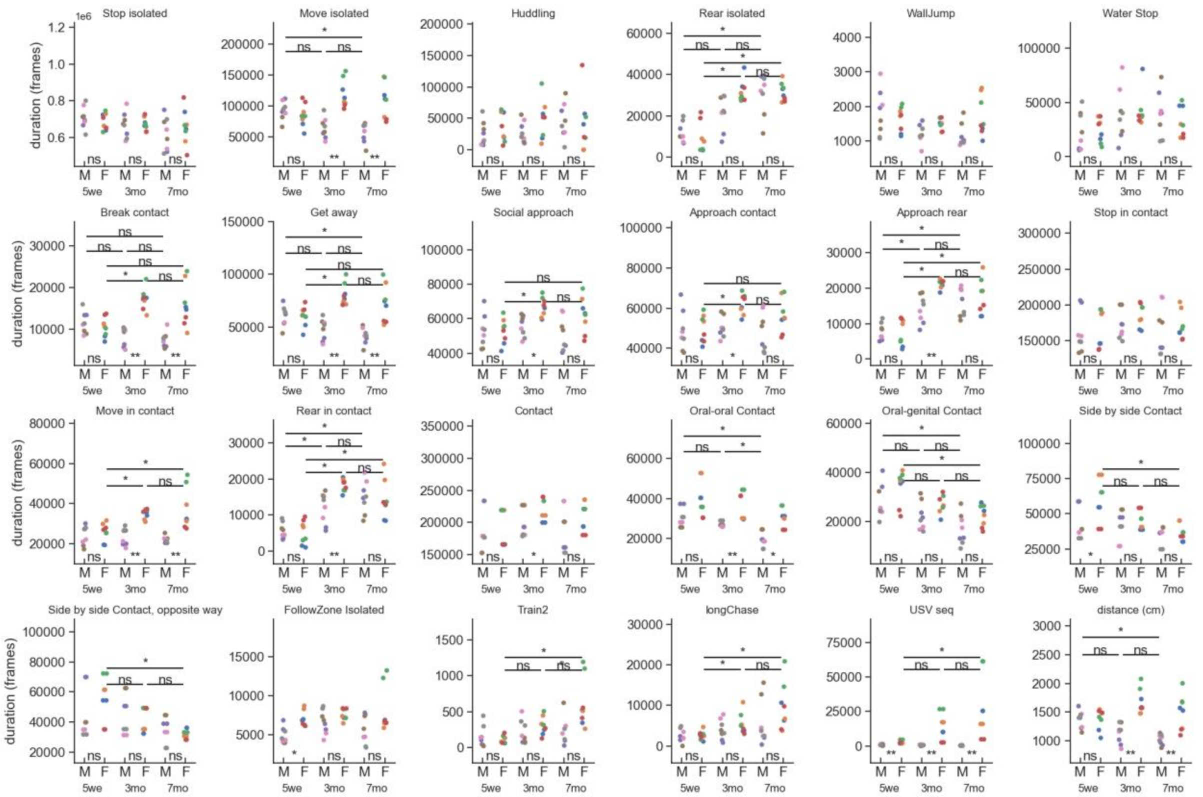


**Supplementary Figure S20.** Behavioral profiles of WT males and females aged of 5 weeks, 3 months and 7 months in the first (a), second (b) and third (c) nights of the experiment. We depicted the total time spent in each behavior for each individual. The colors of the markers match the pairs.

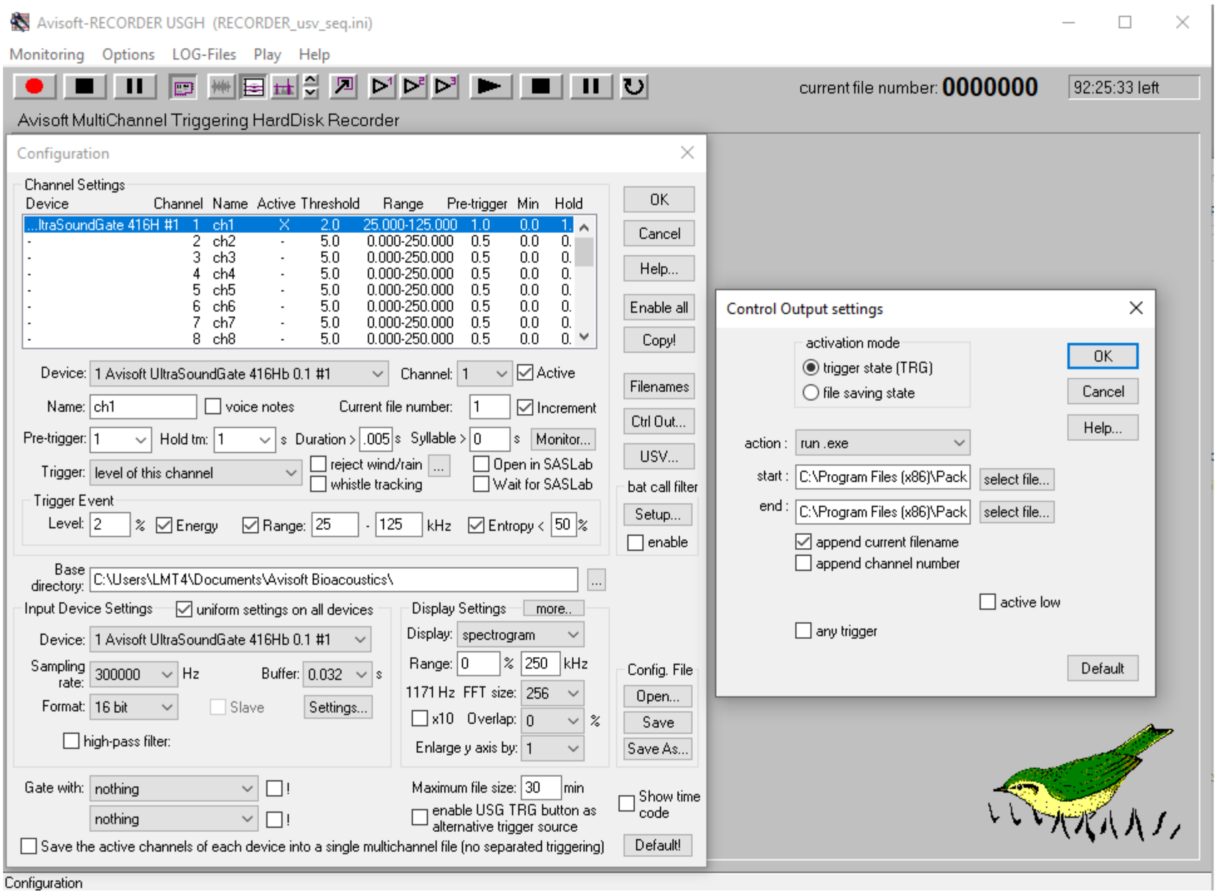


**Supplementary Figure S21.** Avisoft-RECORDER screenshot with the recording parameters.

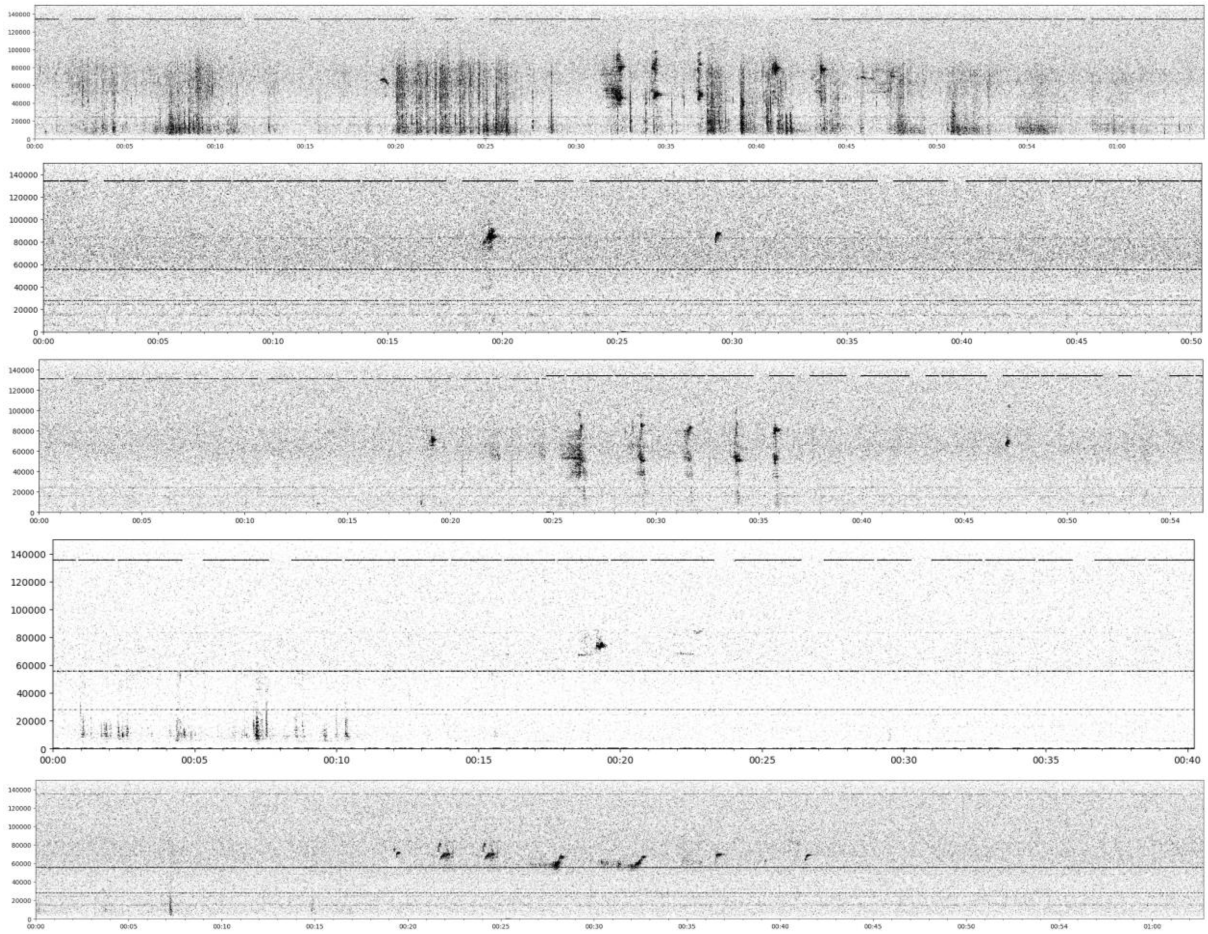


**Supplementary Figure S22.** Noise types. **a**) Example of low power signal **b**) Noise produced by the animals moving in the bedding in the environment. **c**) An extreme case with 5 different noise sources: at 135 kHz, a strong signal at a constant frequency that is appearing and disappearing randomly. At 55 kHz, a blinking signal at 100 Hz. At 85 kHz, 28 kHz, 25 kHz, 18 kHz, other perturbing signal signals. **d**) A typical example where the noisy signal above 120 kHz shifts in frequency. **e**) An example of strong noise emitted in at the same frequency range as the USVs.

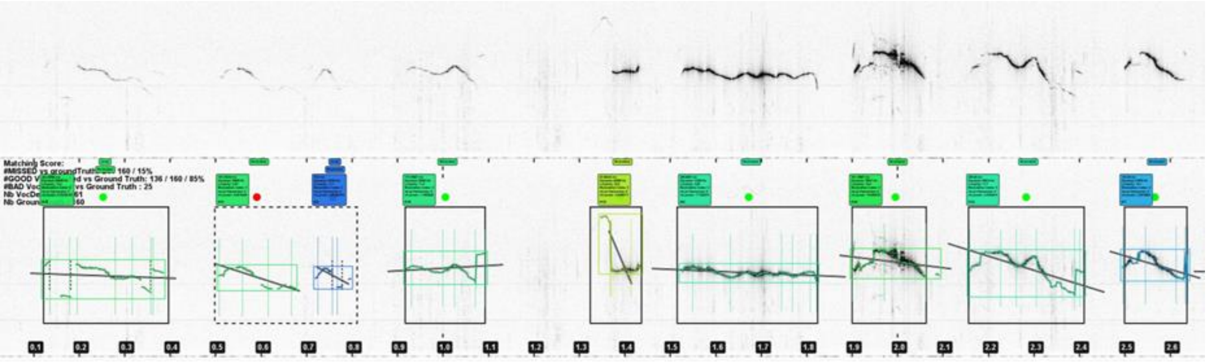


**Supplementary Figure S23.** Validation. Black rectangles show matching validation (in time) and dotted rectangles show mismatching USVs. In the case showed here, the second segmentation is divided into two USVs by the algorithm where the expert had merge it, causing a mismatch.

**Supplementary Table II.** Simulation of the random distribution of the events to check whether long events are more likely to include USVs compared to short events in female WT pairs aged 5 weeks.

| | | | | | | | | | nb of cases for the given sign. level | | | |
|---|---|---|---|---|---|---|---|---|---|---|---|---|
| experiment | event | MW | p original | m_with_usv | sem_with_usv | m_no_usv | sem_no_usv | difference | 0.1 | 0.05 | 0.01 | 0.001 |
| female 5we C57BL/6J WT F13-F14 | Approach contact | 3488125,5 | 1,901E-82 | 8,370 | 0,347 | 3,154 | 0,010 | 2,653 | 179 | 83 | 10 | 0 |
| female 5we C57BL/6J WT F15-F16 | Approach contact | 2451310,5 | 7,719E-55 | 9,056 | 0,546 | 3,078 | 0,010 | 2,942 | 223 | 113 | 30 | 0 |
| female 5we C57BL/6J WT F17-F18 | Approach contact | 7479377,5 | 7,802E-134 | 9,332 | 0,356 | 3,307 | 0,011 | 2,822 | 197 | 96 | 11 | 1 |
| female 5we C57BL/6J WT F19-F20 | Approach contact | 22110559 | 0,000E+00 | 9,586 | 0,220 | 3,491 | 0,012 | 2,746 | 220 | 104 | 15 | 2 |
| female 5we C57BL/6J WT F13-F14 | Break contact | 119691 | 3,133E-01 | 14,579 | 1,973 | 16,144 | 0,236 | 0,903 | 203 | 106 | 35 | 7 |
| female 5we C57BL/6J WT F15-F16 | Break contact | 35104 | 1,324E-01 | 15,176 | 1,916 | 16,079 | 0,231 | 0,944 | 206 | 88 | 18 | 0 |
| female 5we C57BL/6J WT F17-F18 | Break contact | 55579,5 | 2,892E-01 | 15,100 | 3,097 | 15,594 | 0,201 | 0,968 | 200 | 95 | 16 | 3 |
| female 5we C57BL/6J WT F19-F20 | Break contact | 330688,5 | 4,046E-02 | 16,165 | 0,969 | 17,574 | 0,231 | 0,920 | 202 | 97 | 24 | 2 |
| female 5we C57BL/6J WT F13-F14 | Contact | 207119 | 5,760E-32 | 175,593 | 33,572 | 92,872 | 5,853 | 1,891 | 192 | 102 | 19 | 3 |
| female 5we C57BL/6J WT F15-F16 | Contact | 44612,5 | 8,734E-16 | 208,622 | 30,925 | 99,244 | 5,791 | 2,102 | 170 | 77 | 13 | 0 |
| female 5we C57BL/6J WT F17-F18 | Contact | 114517 | 1,987E-34 | 491,361 | 97,965 | 78,623 | 4,336 | 6,250 | 183 | 100 | 18 | 1 |
| female 5we C57BL/6J WT F19-F20 | Contact | 568099,5 | 2,987E-77 | 321,918 | 37,074 | 81,668 | 4,284 | 3,942 | 200 | 92 | 20 | 4 |
| female 5we C57BL/6J WT F13-F14 | FollowZone Isolated | 206088,5 | 1,898E-43 | 6,659 | 0,657 | 1,945 | 0,011 | 3,424 | 182 | 83 | 11 | 2 |
| female 5we C57BL/6J WT F15-F16 | FollowZone Isolated | 94388 | 2,271E-14 | 6,200 | 0,876 | 1,907 | 0,009 | 3,251 | 200 | 113 | 24 | 4 |
| female 5we C57BL/6J WT F17-F18 | FollowZone Isolated | 241778,5 | 8,543E-18 | 4,822 | 0,689 | 1,984 | 0,010 | 2,431 | 210 | 101 | 26 | 1 |
| female 5we C57BL/6J WT F19-F20 | FollowZone Isolated | 1472371,5 | 1,534E-190 | 7,379 | 0,369 | 2,013 | 0,011 | 3,665 | 186 | 96 | 19 | 2 |
| female 5we C57BL/6J WT F13-F14 | Get away | 4837611,5 | 1,041E-47 | 6,652 | 0,588 | 3,525 | 0,015 | 1,887 | 202 | 106 | 24 | 0 |
| female 5we C57BL/6J WT F15-F16 | Get away | 3423256 | 5,535E-27 | 5,221 | 0,428 | 3,282 | 0,013 | 1,591 | 198 | 93 | 13 | 4 |
| female 5we C57BL/6J WT F17-F18 | Get away | 7872846,5 | 2,205E-36 | 5,784 | 0,541 | 3,497 | 0,013 | 1,654 | 188 | 94 | 20 | 6 |
| female 5we C57BL/6J WT F19-F20 | Get away | 31129770 | 2,478E-186 | 6,587 | 0,240 | 3,691 | 0,015 | 1,784 | 180 | 92 | 20 | 3 |
| female 5we C57BL/6J WT F13-F14 | longChase | 96 | 4,188E-03 | 403,417 | 25,207 | 313,000 | 46,404 | 1,289 | 258 | 143 | 35 | 6 |
| female 5we C57BL/6J WT F15-F16 | longChase | 139 | 2,000E-01 | 485,375 | 113,265 | 401,488 | 31,681 | 1,209 | 219 | 101 | 21 | 1 |
| female 5we C57BL/6J WT F17-F18 | longChase | 125 | 1,767E-02 | 521,500 | 46,242 | 391,659 | 34,253 | 1,332 | 257 | 134 | 31 | 5 |
| female 5we C57BL/6J WT F19-F20 | longChase | 918,5 | 1,247E-04 | 414,818 | 16,104 | 322,625 | 18,885 | 1,286 | 261 | 151 | 42 | 8 |
| female 5we C57BL/6J WT F13-F14 | Oral-genital Contact | 1406877 | 1,704E-31 | 13,814 | 1,378 | 7,585 | 0,115 | 1,821 | 196 | 112 | 19 | 1 |
| female 5we C57BL/6J WT F15-F16 | Oral-genital Contact | 1115020 | 1,308E-14 | 10,169 | 1,421 | 6,397 | 0,070 | 1,590 | 215 | 124 | 24 | 3 |
| female 5we C57BL/6J WT F17-F18 | Oral-genital Contact | 2385256,5 | 1,342E-39 | 12,138 | 1,072 | 6,647 | 0,074 | 1,826 | 215 | 93 | 22 | 3 |
| female 5we C57BL/6J WT F19-F20 | Oral-genital Contact | 6735609 | 4,702E-89 | 9,057 | 0,479 | 6,005 | 0,095 | 1,508 | 206 | 101 | 10 | 1 |
| female 5we C57BL/6J WT F13-F14 | Train2 | 774 | 6,028E-02 | 9,318 | 1,069 | 7,326 | 0,303 | 1,272 | 244 | 128 | 33 | 1 |
| female 5we C57BL/6J WT F15-F16 | Train2 | 133 | 2,560E-01 | 7,500 | 0,901 | 7,524 | 0,413 | 0,997 | 213 | 87 | 15 | 1 |
| female 5we C57BL/6J WT F17-F18 | Train2 | 205,5 | 9,903E-02 | 5,800 | 0,716 | 7,287 | 0,384 | 0,796 | 223 | 104 | 23 | 0 |
| female 5we C57BL/6J WT F19-F20 | Train2 | 4519,5 | 6,320E-03 | 10,000 | 0,718 | 7,865 | 0,262 | 1,271 | 244 | 140 | 30 | 8 |

**Supplementary Table III.** Simulation of the random distribution of the events to check whether long events are more likely to include USVs compared to short events in female WT pairs aged 3 months.

| | | | | | | | | | nb of cases for the given sign. level | | | |
|---|---|---|---|---|---|---|---|---|---|---|---|---|
| experiment | event | MW | p original | m_with_usv | sem_with_usv | m_no_usv | sem_no_usv | difference | 0.1 | 0.05 | 0.01 | 0.001 |
| female 3mo C57BL/6J WT F13-F14 | Approach contact | 30576122 | 0,000E+00 | 9,762 | 0,204 | 4,086 | 0,015 | 2,389 | 215 | 111 | 26 | 5 |
| female 3mo C57BL/6J WT F15-F16 | Approach contact | 34251384 | 0,000E+00 | 9,906 | 0,176 | 4,059 | 0,016 | 2,441 | 185 | 88 | 17 | 0 |
| female 3mo C57BL/6J WT F17-F18 | Approach contact | 81250488 | 0,000E+00 | 9,627 | 0,124 | 4,187 | 0,014 | 2,299 | 234 | 129 | 29 | 5 |
| female 3mo C57BL/6J WT F19-F20 | Approach contact | 34522233 | 0,000E+00 | 10,731 | 0,194 | 3,634 | 0,012 | 2,953 | 205 | 85 | 14 | 2 |
| female 3mo C57BL/6J WT F13-F14 | Break contact | 762830 | 6,114E-05 | 18,217 | 1,022 | 17,174 | 0,189 | 1,061 | 230 | 118 | 26 | 2 |
| female 3mo C57BL/6J WT F15-F16 | Break contact | 560652,5 | 4,887E-04 | 19,237 | 1,052 | 18,615 | 0,235 | 1,033 | 216 | 106 | 19 | 2 |
| female 3mo C57BL/6J WT F17-F18 | Break contact | 1708672,5 | 1,047E-04 | 17,441 | 0,717 | 17,305 | 0,162 | 1,008 | 213 | 110 | 25 | 3 |
| female 3mo C57BL/6J WT F19-F20 | Break contact | 563094 | 5,103E-10 | 20,088 | 0,992 | 17,864 | 0,239 | 1,124 | 175 | 96 | 24 | 1 |
| female 3mo C57BL/6J WT F13-F14 | Contact | 1260938,5 | 3,373E-92 | 143,351 | 12,715 | 61,235 | 2,731 | 2,341 | 219 | 122 | 22 | 4 |
| female 3mo C57BL/6J WT F15-F16 | Contact | 1089411,5 | 9,212E-113 | 208,560 | 26,430 | 66,089 | 3,235 | 3,156 | 192 | 82 | 22 | 3 |
| female 3mo C57BL/6J WT F17-F18 | Contact | 2682544,5 | 2,717E-230 | 226,002 | 21,955 | 50,777 | 2,269 | 4,451 | 217 | 105 | 23 | 1 |
| female 3mo C57BL/6J WT F19-F20 | Contact | 1101252,5 | 1,618E-126 | 269,733 | 38,791 | 77,977 | 4,114 | 3,459 | 213 | 109 | 20 | 2 |
| female 3mo C57BL/6J WT F13-F14 | Isolated | 2754402 | 3,242E-284 | 8,340 | 0,377 | 2,083 | 0,013 | 4,004 | 183 | 92 | 21 | 4 |
| female 3mo C57BL/6J WT F15-F16 | Isolated | 1616222 | 6,493E-237 | 7,599 | 0,370 | 2,123 | 0,017 | 3,580 | 201 | 100 | 15 | 2 |
| female 3mo C57BL/6J WT F17-F18 | Isolated | 5311084 | 0,000E+00 | 7,051 | 0,208 | 2,121 | 0,014 | 3,324 | 197 | 103 | 17 | 4 |
| female 3mo C57BL/6J WT F19-F20 | Isolated | 1907159 | 8,266E-302 | 9,436 | 0,393 | 2,024 | 0,012 | 4,662 | 185 | 96 | 15 | 1 |
| female 3mo C57BL/6J WT F13-F14 | Get away | 57034657 | 9,043E-197 | 6,424 | 0,196 | 4,189 | 0,016 | 1,533 | 198 | 101 | 23 | 5 |
| female 3mo C57BL/6J WT F15-F16 | Get away | 48725239 | 3,262E-235 | 6,303 | 0,179 | 3,976 | 0,017 | 1,585 | 178 | 101 | 26 | 3 |
| female 3mo C57BL/6J WT F17-F18 | Get away | 164832302 | 0,000E+00 | 5,795 | 0,115 | 4,503 | 0,016 | 1,287 | 196 | 98 | 17 | 2 |
| female 3mo C57BL/6J WT F19-F20 | Get away | 50793332 | 3,071E-306 | 7,007 | 0,230 | 3,740 | 0,016 | 1,873 | 198 | 106 | 29 | 3 |
| female 3mo C57BL/6J WT F13-F14 | longChase | 1500,5 | 3,653E-01 | 360,700 | 26,338 | 362,269 | 24,655 | 0,996 | 267 | 155 | 37 | 4 |
| female 3mo C57BL/6J WT F15-F16 | longChase | 936 | 1,914E-07 | 468,764 | 24,945 | 302,745 | 21,669 | 1,548 | 252 | 133 | 34 | 6 |
| female 3mo C57BL/6J WT F17-F18 | longChase | 2227,5 | 7,102E-04 | 403,548 | 17,462 | 317,000 | 21,642 | 1,273 | 255 | 140 | 39 | 9 |
| female 3mo C57BL/6J WT F19-F20 | longChase | 1461 | 5,228E-04 | 395,056 | 14,274 | 325,367 | 19,167 | 1,214 | 263 | 142 | 29 | 5 |
| female 3mo C57BL/6J WT F13-F14 | Contact | 7057443 | 2,656E-177 | 11,665 | 0,573 | 4,996 | 0,088 | 2,335 | 216 | 95 | 15 | 3 |
| female 3mo C57BL/6J WT F15-F16 | Contact | 5313944,5 | 1,860E-162 | 13,755 | 0,711 | 5,443 | 0,091 | 2,527 | 217 | 111 | 19 | 5 |
| female 3mo C57BL/6J WT F17-F18 | Contact | 15302825 | 1,308E-212 | 10,077 | 0,461 | 5,877 | 0,129 | 1,715 | 197 | 90 | 22 | 4 |
| female 3mo C57BL/6J WT F19-F20 | Contact | 7842947,5 | 2,650E-145 | 10,033 | 0,503 | 5,591 | 0,129 | 1,795 | 208 | 103 | 23 | 2 |
| female 3mo C57BL/6J WT F13-F14 | Train2 | 9433 | 1,617E-02 | 9,793 | 0,533 | 8,676 | 0,395 | 1,129 | 225 | 126 | 33 | 5 |
| female 3mo C57BL/6J WT F15-F16 | Train2 | 8251 | 1,396E-03 | 9,854 | 0,535 | 8,677 | 0,303 | 1,136 | 269 | 157 | 31 | 5 |
| female 3mo C57BL/6J WT F17-F18 | Train2 | 15998,5 | 2,428E-04 | 9,924 | 0,439 | 8,431 | 0,242 | 1,177 | 250 | 129 | 37 | 5 |
| female 3mo C57BL/6J WT F19-F20 | Train2 | 9437,5 | 9,488E-05 | 10,269 | 0,505 | 8,314 | 0,317 | 1,235 | 245 | 136 | 33 | 3 |

**Supplementary Table IV.** Simulation of the random distribution of the events to check whether long events are more likely to include USVs compared to short events in female WT pairs aged 7 months.

| | | | | | | | | | nb of cases for the given sign. level | | | |
|---|---|---|---|---|---|---|---|---|---|---|---|---|
| experiment | event | MW | p original | m_with_usv | sem_with_usv | m_no_usv | sem_no_usv | difference | 0.1 | 0.05 | 0.01 | 0.001 |
| female 7mo C57BL/6J WT F13-F14 | Approach contact | 46601580 | 0,000E+00 | 10,015 | 0,148 | 3,759 | 0,013 | 2,664 | 196 | 95 | 20 | 1 |
| female 7mo C57BL/6J WT F15-F16 | Approach contact | 9967619,5 | 6,203E-134 | 10,110 | 0,354 | 4,068 | 0,015 | 2,486 | 200 | 101 | 22 | 5 |
| female 7mo C57BL/6J WT F17-F18 | Approach contact | 144892175 | 0,000E+00 | 9,815 | 0,087 | 4,469 | 0,018 | 2,196 | 210 | 112 | 17 | 2 |
| female 7mo C57BL/6J WT F19-F20 | Approach contact | 13717518,5 | 3,763E-196 | 10,260 | 0,305 | 3,914 | 0,014 | 2,621 | 188 | 87 | 21 | 1 |
| female 7mo C57BL/6J WT F13-F14 | Break contact | 1093400,5 | 3,284E-07 | 17,916 | 0,678 | 18,178 | 0,247 | 0,986 | 206 | 93 | 22 | 3 |
| female 7mo C57BL/6J WT F15-F16 | Break contact | 157677,5 | 2,365E-02 | 19,931 | 2,132 | 18,900 | 0,248 | 1,055 | 208 | 98 | 18 | 1 |
| female 7mo C57BL/6J WT F17-F18 | Break contact | 3731687,5 | 4,054E-06 | 18,562 | 0,495 | 18,205 | 0,182 | 1,020 | 227 | 126 | 21 | 4 |
| female 7mo C57BL/6J WT F19-F20 | Break contact | 272302,5 | 9,110E-02 | 15,657 | 1,035 | 18,538 | 0,260 | 0,845 | 192 | 94 | 12 | 3 |
| female 7mo C57BL/6J WT F13-F14 | Contact | 1807688 | 8,328E-175 | 130,690 | 7,396 | 71,623 | 3,659 | 1,825 | 204 | 97 | 23 | 5 |
| female 7mo C57BL/6J WT F15-F16 | Contact | 391556 | 6,554E-42 | 170,167 | 20,733 | 76,898 | 3,781 | 2,213 | 196 | 100 | 21 | 2 |
| female 7mo C57BL/6J WT F17-F18 | Contact | 5463175,5 | 0,000E+00 | 132,430 | 6,161 | 45,167 | 2,111 | 2,932 | 219 | 108 | 22 | 4 |
| female 7mo C57BL/6J WT F19-F20 | Contact | 493327,5 | 1,092E-62 | 194,088 | 20,966 | 80,518 | 4,072 | 2,411 | 206 | 110 | 27 | 0 |
| female 7mo C57BL/6J WT F13-F14 | FollowZone Isolated | 2869211,5 | 0,000E+00 | 7,625 | 0,265 | 2,086 | 0,014 | 3,656 | 197 | 91 | 21 | 1 |
| female 7mo C57BL/6J WT F15-F16 | FollowZone Isolated | 388873,5 | 2,863E-76 | 7,365 | 0,720 | 1,973 | 0,015 | 3,733 | 207 | 99 | 12 | 0 |
| female 7mo C57BL/6J WT F17-F18 | FollowZone Isolated | 17836497,5 | 0,000E+00 | 7,184 | 0,131 | 2,162 | 0,013 | 3,323 | 221 | 109 | 29 | 8 |
| female 7mo C57BL/6J WT F19-F20 | FollowZone Isolated | 1001142,5 | 6,926E-156 | 9,562 | 0,576 | 2,232 | 0,018 | 4,284 | 192 | 89 | 15 | 0 |
| female 7mo C57BL/6J WT F13-F14 | Get away | 90165187,5 | 0,000E+00 | 6,626 | 0,156 | 4,270 | 0,019 | 1,552 | 188 | 92 | 20 | 2 |
| female 7mo C57BL/6J WT F15-F16 | Get away | 16193472,5 | 2,022E-79 | 6,495 | 0,382 | 3,820 | 0,016 | 1,701 | 204 | 107 | 21 | 3 |
| female 7mo C57BL/6J WT F17-F18 | Get away | 368443480 | 0,000E+00 | 6,396 | 0,090 | 4,399 | 0,017 | 1,454 | 222 | 106 | 24 | 2 |
| female 7mo C57BL/6J WT F19-F20 | Get away | 20425935,5 | 4,077E-126 | 5,927 | 0,222 | 3,797 | 0,018 | 1,561 | 206 | 97 | 21 | 2 |
| female 7mo C57BL/6J WT F13-F14 | longChase | 840 | 7,914E-02 | 407,333 | 17,029 | 377,250 | 50,768 | 1,080 | 248 | 145 | 40 | 5 |
| female 7mo C57BL/6J WT F15-F16 | longChase | 274,5 | 1,655E-03 | 489,364 | 45,804 | 330,844 | 24,398 | 1,479 | 240 | 126 | 32 | 4 |
| female 7mo C57BL/6J WT F17-F18 | longChase | 3048 | 3,536E-05 | 434,621 | 12,999 | 308,439 | 20,871 | 1,409 | 254 | 141 | 37 | 5 |
| female 7mo C57BL/6J WT F19-F20 | longChase | 1008 | 2,405E-04 | 453,667 | 21,312 | 348,702 | 21,249 | 1,301 | 247 | 151 | 36 | 6 |
| female 7mo C57BL/6J WT F13-F14 | Oral-genital Contact | 8646383,5 | 2,180E-212 | 9,820 | 0,377 | 5,288 | 0,102 | 1,857 | 198 | 102 | 18 | 0 |
| female 7mo C57BL/6J WT F15-F16 | Oral-genital Contact | 1502113,5 | 9,671E-57 | 13,895 | 1,460 | 5,104 | 0,129 | 2,722 | 200 | 106 | 27 | 4 |
| female 7mo C57BL/6J WT F17-F18 | Oral-genital Contact | 31184996,5 | 0,000E+00 | 8,818 | 0,199 | 4,730 | 0,086 | 1,864 | 193 | 101 | 26 | 2 |
| female 7mo C57BL/6J WT F19-F20 | Oral-genital Contact | 2481969,5 | 1,678E-110 | 12,440 | 0,751 | 4,977 | 0,130 | 2,499 | 198 | 83 | 16 | 3 |
| female 7mo C57BL/6J WT F13-F14 | Train2 | 10645 | 2,456E-02 | 8,973 | 0,409 | 8,192 | 0,399 | 1,095 | 264 | 158 | 42 | 14 |
| female 7mo C57BL/6J WT F15-F16 | Train2 | 1157,5 | 1,454E-01 | 11,647 | 2,491 | 8,534 | 0,351 | 1,365 | 225 | 136 | 32 | 3 |
| female 7mo C57BL/6J WT F17-F18 | Train2 | 56511,5 | 2,705E-11 | 9,906 | 0,245 | 7,653 | 0,240 | 1,294 | 242 | 141 | 42 | 8 |
| female 7mo C57BL/6J WT F19-F20 | Train2 | 8181 | 3,089E-02 | 10,391 | 0,775 | 9,032 | 0,297 | 1,150 | 247 | 133 | 34 | 3 |

**Supplementary Table V.** Simulation of the random distribution of the events to check whether long events are more likely to include USVs compared to short events in female Shank3-/- pairs aged 3 months.

| | | | | | | | | | nb of cases for the given sign. level | | | |
|---|---|---|---|---|---|---|---|---|---|---|---|---|
| experiment | event | MW | p original | m_with_usv | sem_with_usv | m_no_usv | sem_no_usv | difference | 0.1 | 0.05 | 0.01 | 0.001 |
| female 3mo Shank3 KO 4417496_4419810 | Approach contact | 44324525 | 0,000E+00 | 9,177 | 0,157 | 4,120 | 0,012 | 2,227 | 197 | 97 | 16 | 0 |
| female 3mo Shank3 KO 4724356-4724113 | Approach contact | 23659868 | 2,533E-301 | 10,339 | 0,254 | 3,860 | 0,012 | 2,678 | 192 | 89 | 23 | 2 |
| female 3mo Shank3 KO 4849069-4849197 | Approach contact | 17701431 | 1,061E-214 | 9,195 | 0,233 | 3,954 | 0,013 | 2,326 | 217 | 109 | 17 | 1 |
| female 3mo Shank3 KO 4849144-4849294 | Approach contact | 75390671 | 0,000E+00 | 9,614 | 0,148 | 3,655 | 0,011 | 2,630 | 202 | 105 | 23 | 2 |
| female 3mo Shank3 KO 4849155_4849457 | Approach contact | 22951613 | 2,492e-310 | 9,718 | 0,248 | 3,416 | 0,010 | 2,845 | 215 | 105 | 14 | 0 |
| female 3mo Shank3 KO 4849484_4849378 | Approach contact | 71699455 | 0,000E+00 | 10,257 | 0,141 | 3,513 | 0,010 | 2,920 | 204 | 106 | 19 | 3 |
| female 3mo Shank3 KO 4417496_4419810 | Break contact | 878670,5 | 1,975E-16 | 17,510 | 0,690 | 14,172 | 0,144 | 1,236 | 213 | 107 | 20 | 5 |
| female 3mo Shank3 KO 4724356-4724113 | Break contact | 431980 | 1,184E-09 | 16,307 | 1,048 | 13,132 | 0,155 | 1,242 | 197 | 106 | 23 | 5 |
| female 3mo Shank3 KO 4849069-4849197 | Break contact | 380647,5 | 1,468E-01 | 14,865 | 0,875 | 17,198 | 0,187 | 0,864 | 202 | 95 | 17 | 2 |
| female 3mo Shank3 KO 4849144-4849294 | Break contact | 1106371,5 | 2,417E-14 | 17,155 | 0,724 | 14,231 | 0,167 | 1,205 | 198 | 108 | 32 | 2 |
| female 3mo Shank3 KO 4849155_4849457 | Break contact | 118758 | 6,480E-10 | 18,588 | 1,631 | 12,210 | 0,179 | 1,522 | 229 | 104 | 20 | 2 |
| female 3mo Shank3 KO 4849484_4849378 | Break contact | 1494891 | 1,375E-42 | 18,680 | 0,696 | 12,589 | 0,146 | 1,484 | 216 | 110 | 25 | 5 |
| female 3mo Shank3 KO 4417496_4419810 | Contact | 1817302 | 3,556E-133 | 183,534 | 14,278 | 63,579 | 2,376 | 2,887 | 203 | 103 | 19 | 2 |
| female 3mo Shank3 KO 4724356-4724113 | Contact | 744948,5 | 7,089E-132 | 274,905 | 39,932 | 67,131 | 3,277 | 4,095 | 205 | 103 | 27 | 2 |
| female 3mo Shank3 KO 4849069-4849197 | Contact | 497752,5 | 7,416E-85 | 382,148 | 50,282 | 69,430 | 3,059 | 5,504 | 205 | 91 | 12 | 3 |
| female 3mo Shank3 KO 4849144-4849294 | Contact | 1937683 | 7,018E-192 | 249,412 | 26,054 | 75,514 | 3,686 | 3,303 | 201 | 110 | 33 | 4 |
| female 3mo Shank3 KO 4849155_4849457 | Contact | 364730,5 | 8,206E-94 | 388,640 | 56,701 | 84,372 | 4,419 | 4,606 | 198 | 107 | 20 | 0 |
| female 3mo Shank3 KO 4849484_4849378 | Contact | 3142082 | 1,669E-270 | 254,353 | 27,430 | 52,134 | 2,545 | 4,879 | 170 | 89 | 21 | 2 |
| female 3mo Shank3 KO 4417496_4419810 | FollowZone Isolated | 3777611 | 0,000E+00 | 8,457 | 0,298 | 2,225 | 0,014 | 3,801 | 192 | 88 | 15 | 0 |
| female 3mo Shank3 KO 4724356-4724113 | FollowZone Isolated | 1012911,5 | 1,265E-174 | 9,216 | 0,539 | 2,252 | 0,019 | 4,092 | 221 | 110 | 33 | 4 |
| female 3mo Shank3 KO 4849069-4849197 | FollowZone Isolated | 951229 | 5,961E-171 | 7,762 | 0,381 | 2,070 | 0,012 | 3,749 | 193 | 88 | 14 | 2 |
| female 3mo Shank3 KO 4849144-4849294 | FollowZone Isolated | 3153138,5 | 0,000E+00 | 7,280 | 0,245 | 2,029 | 0,012 | 3,588 | 200 | 109 | 21 | 0 |
| female 3mo Shank3 KO 4849155_4849457 | FollowZone Isolated | 640640 | 6,784E-101 | 5,228 | 0,282 | 2,023 | 0,015 | 2,584 | 193 | 91 | 18 | 2 |
| female 3mo Shank3 KO 4849484_4849378 | FollowZone Isolated | 3485305 | 0,000E+00 | 8,550 | 0,266 | 2,033 | 0,012 | 4,205 | 184 | 94 | 14 | 0 |
| female 3mo Shank3 KO 4417496_4419810 | Get away | 68692201 | 2,857E-277 | 6,668 | 0,170 | 3,887 | 0,014 | 1,715 | 206 | 95 | 21 | 0 |
| female 3mo Shank3 KO 4724356-4724113 | Get away | 27204842 | 3,974E-169 | 6,312 | 0,207 | 3,710 | 0,015 | 1,701 | 209 | 120 | 25 | 2 |
| female 3mo Shank3 KO 4849069-4849197 | Get away | 33237709 | 9,555E-100 | 6,123 | 0,252 | 4,393 | 0,017 | 1,394 | 195 | 92 | 20 | 0 |
| female 3mo Shank3 KO 4849144-4849294 | Get away | 87571698 | 0,000E+00 | 6,587 | 0,165 | 3,704 | 0,015 | 1,778 | 231 | 108 | 24 | 1 |
| female 3mo Shank3 KO 4849155_4849457 | Get away | 22867601 | 2,464E-155 | 5,138 | 0,228 | 3,121 | 0,013 | 1,646 | 214 | 105 | 24 | 5 |
| female 3mo Shank3 KO 4849484_4849378 | Get away | 90206325 | 0,000E+00 | 7,060 | 0,165 | 3,625 | 0,015 | 1,948 | 206 | 115 | 25 | 2 |
| female 3mo Shank3 KO 4417496_4419810 | longChase | 5210,5 | 4,074E-02 | 360,201 | 11,650 | 327,821 | 14,977 | 1,099 | 281 | 165 | 50 | 8 |
| female 3mo Shank3 KO 4724356-4724113 | longChase | 2456 | 2,524E-02 | 463,705 | 22,573 | 428,597 | 30,932 | 1,082 | 254 | 149 | 40 | 9 |
| female 3mo Shank3 KO 4849069-4849197 | longChase | 370,5 | 2,877E-02 | 360,356 | 20,934 | 302,435 | 28,567 | 1,192 | 244 | 145 | 39 | 5 |
| female 3mo Shank3 KO 4849144-4849294 | longChase | 837 | 1,534E-01 | 347,714 | 11,414 | 347,750 | 38,979 | 1,000 | 242 | 150 | 40 | 5 |
| female 3mo Shank3 KO 4849155_4849457 | longChase | 212 | 3,791E-01 | 396,389 | 49,528 | 388,520 | 29,240 | 1,020 | 261 | 148 | 33 | 3 |
| female 3mo Shank3 KO 4849484_4849378 | longChase | 1251,5 | 3,620E-03 | 397,606 | 15,006 | 313,000 | 29,499 | 1,270 | 268 | 144 | 40 | 6 |
| female 3mo Shank3 KO 4417496_4419810 | Oral-genital Contact | 8341606 | 8,778E-112 | 9,686 | 0,421 | 6,514 | 0,103 | 1,487 | 176 | 78 | 12 | 1 |
| female 3mo Shank3 KO 4724356-4724113 | Oral-genital Contact | 4162072,5 | 2,796E-98 | 15,068 | 0,869 | 7,561 | 0,270 | 1,993 | 205 | 92 | 16 | 3 |
| female 3mo Shank3 KO 4849069-4849197 | Oral-genital Contact | 2959622,5 | 2,022E-47 | 9,113 | 0,683 | 6,377 | 0,109 | 1,429 | 217 | 120 | 27 | 2 |
| female 3mo Shank3 KO 4849144-4849294 | Oral-genital Contact | 9991739 | 1,251E-166 | 10,394 | 0,492 | 6,465 | 0,120 | 1,608 | 208 | 104 | 24 | 3 |
| female 3mo Shank3 KO 4849155_4849457 | Oral-genital Contact | 2441714 | 2,136E-76 | 13,737 | 1,258 | 7,279 | 0,365 | 1,887 | 216 | 100 | 18 | 3 |
| female 3mo Shank3 KO 4849484_4849378 | Oral-genital Contact | 12855142 | 5,398E-289 | 14,274 | 0,562 | 6,898 | 0,154 | 2,069 | 189 | 94 | 20 | 3 |
| female 3mo Shank3 KO 4417496_4419810 | Train2 | 26701 | 2,298E-04 | 9,182 | 0,359 | 7,975 | 0,197 | 1,151 | 263 | 135 | 39 | 6 |
| female 3mo Shank3 KO 4724356-4724113 | Train2 | 12287,5 | 3,252E-03 | 11,727 | 0,908 | 8,854 | 0,230 | 1,325 | 216 | 107 | 29 | 1 |
| female 3mo Shank3 KO 4849069-4849197 | Train2 | 1859 | 2,938E-04 | 9,353 | 0,693 | 7,440 | 0,358 | 1,257 | 262 | 152 | 41 | 5 |
| female 3mo Shank3 KO 4849144-4849294 | Train2 | 4263 | 1,424E-05 | 9,430 | 0,404 | 7,409 | 0,430 | 1,273 | 247 | 142 | 37 | 9 |
| female 3mo Shank3 KO 4849155_4849457 | Train2 | 373,5 | 1,081E-01 | 7,000 | 0,705 | 8,726 | 0,533 | 0,802 | 209 | 114 | 26 | 3 |
| female 3mo Shank3 KO 4849484_4849378 | Train2 | 17834,5 | 8,482E-05 | 9,758 | 0,365 | 8,073 | 0,311 | 1,209 | 247 | 138 | 33 | 6 |
